# Transcriptome-wide analysis of circRNA and RBP profiles and their molecular and clinical relevance for GBM

**DOI:** 10.1101/2024.06.12.598692

**Authors:** J Latowska-Łysiak, Ż Zarębska, MP Sajek, A Grabowska, A Buratin, JO Misiorek, K Kuczyński, S Bortoluzzi, M Żywicki, JG Kosiński, A Rybak-Wolf, R Piestrzeniewicz, AM Barciszewska, K Rolle

**Author notes:** These authors contributed equally to this work (J Latowska-Łysiak, Ż Zarębska, MP Sajek).

## Abstract

Glioblastoma (GBM) is the most aggressive and lethal type of glioma, characterized by aberrant expression of non-coding RNAs including circular RNAs (circRNAs). They might impact cellular processes by interacting with other molecules – like microRNAs or RNA-binding proteins (RBPs). The diagnostic value of circRNAs and circRNAs/RBPs complexes is still largely unknown. To explore circRNAs and RBPs transcripts expression in GBM, we performed and further analyzed RNA-seq data from GBM patients’ primary and recurrent tumor samples. We identified circRNAs differentially expressed in primary tumors, the circRNA progression markers in recurrent GBM samples as well as the expression profile of RBP transcripts. Subsequent analysis allowed us to generate a comprehensive catalog of circRNA-RBP interactions regarding both the RBPs sequestration by circRNA as well as the RBPs involvement in circRNA biogenesis. Furthermore, we demonstrated the clinical potential of circRNAs and RBPs in GBM and proposed them as the stratification markers in the de novo assembled tumor subtypes. Therefore, our transcriptome-wide study specified circRNA-RBP interactions that could play a significant regulatory role in gliomagenesis and GBM progression.

## 1. Introduction

Gliomas are brain neoplasms originating from the glia and constitute the majority of all primary tumors in the central nervous system. They are classified into four grade groups, based on histological, genetic, and prognostic features. Low-grade gliomas (I and II grades) exhibit minor metastasis and recurrence potential, while high-grade gliomas (III and IV grades) present significant recurrence and progression prognosis [1]. Glioblastoma (GBM) - an IV-grade glioma is the most lethal, aggressive, and malignant among brain tumors in adults, with a median survival of 14.6 months post-treatment [2]. Current therapies, which are not curative, include radiotherapy and temozolomide-based chemotherapy [3]. Due to the lack of effective treatments, high internal molecular heterogeneity among GBM tissues, and high rate of recurrence, there is an urgent need to better understand the molecular basis of GBM and improve patient stratification, defining signatures useful as potential diagnostic and prognostic markers and most importantly to identify new therapeutic targets.

The most common molecular stratification parameter of GBM is the presence of mutations in isocitrate dehydrogenase - *IDH1* and *IDH2* genes. These mutations are most frequent in secondary GBM and predict a favorable disease outcome with prolonged median survival [4, 5]. An additional molecular classification of GBM comprises of 4 submolecular groups: proneural, classical, mesenchymal, and neural which are distinguished by specific gene expression patterns, mutations and associated with different tumor aggressiveness and prognosis for treatment response [6].

In recent years, non-coding RNAs (ncRNAs) and particularly circular RNAs (circRNAs) have gained attention as key players in cancer and potential therapeutic targets. CircRNAs are single-stranded RNA molecules produced by “back-splicing”, which joins a downstream 5’ donor splice site and an upstream 3’ acceptor splice site [7]. Due to their intrinsic resistance to exonuclease cleavage, circRNAs have a longer half-life in comparison to their linear counterparts making them promising cancer biomarkers [8–15].

Recent studies have highlighted the role of RNA binding proteins (RBPs) in the biogenesis and maintenance of circRNAs, influencing the transcriptome’s molecular balance [16, 17]. CircRNAs can act also as miRNA and RBPs sponges potentially impacting gene expression and cellular pathways significantly [12, 18–24]. The interactions between circRNAs and RBPs rely on binding motifs and contextual features, such as secondary structure, flanking nucleotide composition, or short non-sequential motifs [25]. Apart from the RBPs binding motifs, the biogenesis of circRNAs can be also facilitated by the presence of the complementary sequences in both flanking regions, like *Alu* elements [26]. Understanding nature and function of such interactions is crucial for understanding GBM, including tumor biology, progression, treatment response and survival rates [27–29].

In this study, we provide the comprehensive atlas of circRNAs and RBP transcripts differentially expressed in GBM. By utilizing deep total RNA-sequencing (RNA-seq) we established the expression profile of circRNAs and extensively studied their interaction with RBPs, including the potential impact on circRNA formation. First, we identified the differentially expressed circRNA in primary GBM patients’ samples (GBM-PRM) as well as in the recurrent GBM samples (GBM-REC). We further experimentally validated the expression of selected circRNAs, their unique back-splice junction sequences and structures. Finally, based on the same dataset we have identified the profiles of RBPs transcripts expression in GBM. The study of circRNA and RBPs transcripts highlighted the potential binding sites within circRNAs and the RBPs possibly involved in circRNA biogenesis. Further analysis correlated circRNA and RBP expression with established GBM subtypes, leading to a new subtype classification that enhances patient stratification and identification of prognostic markers. Our global analysis pointed circRNAs significantly deregulated in GBM, offering new insights into potential diagnostic and therapeutic targets.

## 2. Results

### 2.1 circRNAs and RBPs transcripts are differentially expressed in GBM

#### 2.1.1. Expression profile of circRNA in GBM

To define circRNA and RBPs mRNA expression profiles in primary (GBM-PRM, n=23) and recurrent GBM (GBM-REC, n=3) (Suppl. Tab. 1) in comparison to the healthy brain control (HB, n=52) (Suppl. Tab. 2), RNA sequencing was performed. Transcriptome sequencing approach was designed to simultaneously profile both circRNA and messenger RNA transcripts. We identified a total number of 29,141 circRNAs in all samples (Fig.1A) including more than 7,600 not previously annotated. The majority of identified circRNA were derived from exons (Suppl. Fig. 1A) and their length was similar among all analyzed samples with the median equal: 740, 631 and 555 nucleotides for HB, GBM-PRM and GBM-REC, respectively (Suppl. Fig. 1B-3D). The circRNA transcripts were distributed within all human chromosomes, including the chromosomes 1 and 2 particularly enriched in circRNAs (Suppl. Fig. 2). CircRNA expression profiles showed clear discrimination between GBM-PRM, GBM-REC and HB samples (Suppl. Fig. 3). The differential circRNA expression (DCE) analysis revealed 1270 circRNAs expressed between GBM-PRM and HB tissues with a log2 fold change in the range between -5.6 and 8.9 (adjusted p-value <0.05) where 1132 of circRNAs were downregulated and only 138 were upregulated (Fig. 1C). To search for circRNAs potentially involved in GBM progression we performed further DCE analysis between GBM-PRM and GBM-REC where we detected 3 differentially expressed circRNAs which were subsequently validated (Fig. 1D). To determine relationship between deregulated circRNAs and their linear counterparts we calculated the circular-to-linear ratio (CLR) based on the formula [circ / (circ + lin)]. A CLR value>0.5 indicates that the number of circular RNA molecules is higher than the corresponding mRNA, while a CLR value<0.5 indicates the opposite. Interestingly, the obtained results showed that in the most cases the number of circRNAs was higher than that the number of their parental linear counterparts (CLR>0.5). Moreover, both circular and linear molecules were frequently upregulated, indicating a positive correlation between their expression levels (Fig. 1E, upper right quadrant of the figure).

**Figure 1.**
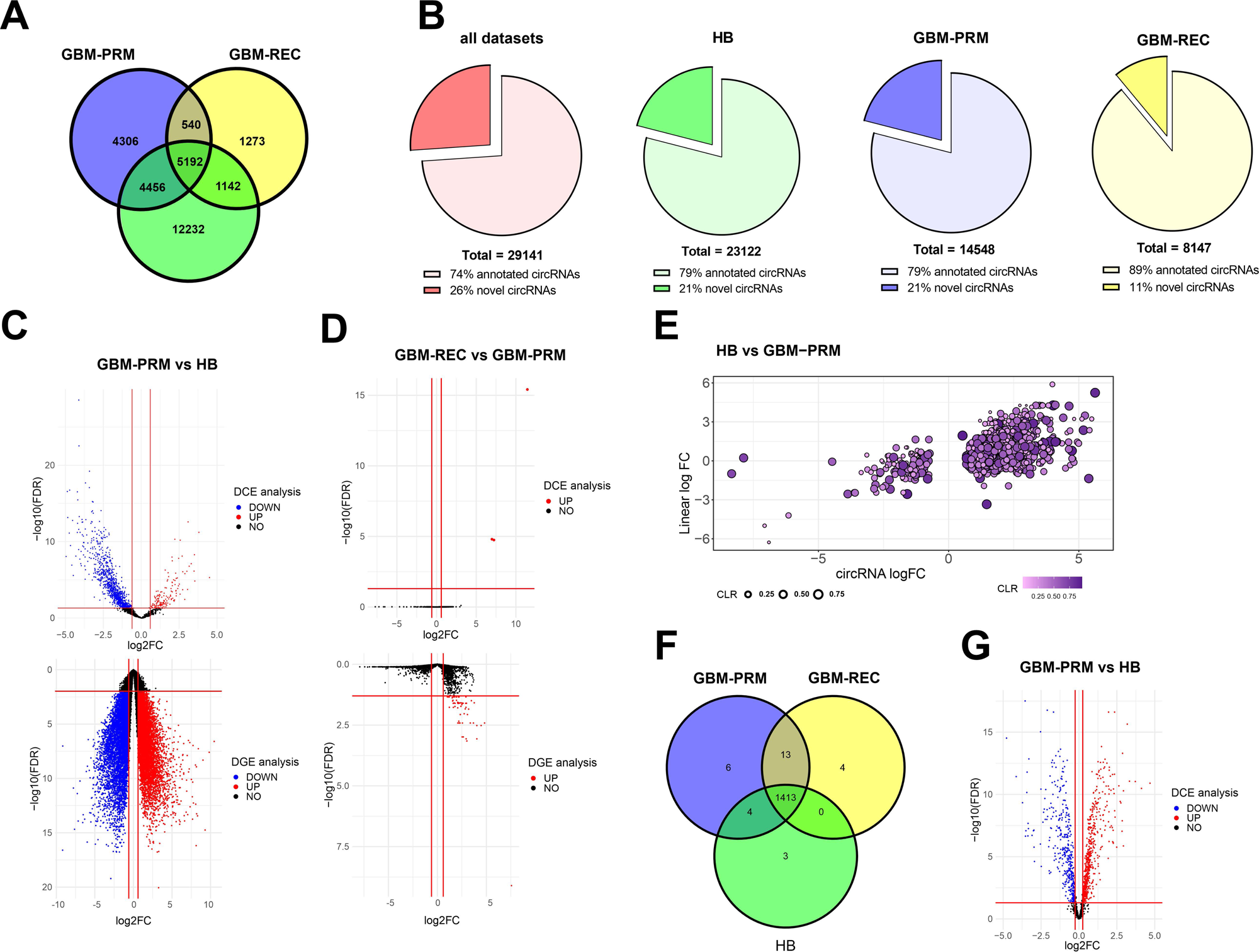
Overview of the identified circRNAs and RBPs in GBM. **A.** Venn diagram illustrates an overlap between circRNAs that were found expressed in analyzed GBM-PRM, GBM-REC tissues and HB control. **B.** Distribution of novel and annotated circRNAs in selected samples. **C.** Volcano plot of differentially expressed circRNAs and mRNAs in GBM-PRM vs HB. The blue and red dots indicate downregulated and upregulated circRNAs, respectively. **D**. Volcano plot of differentially expressed circRNAs in GBM-REC vs GBM-PRM. The blue and red dots indicate downregulated and upregulated circRNAs, respectively. **E.** The graph presents the circular-to-linear ratio (CLR) of circRNAs and their linear counterparts. The log2fC of circRNAs is marked on the X-axis, and the log2FC values of the corresponding mRNAs are marked on the Y-axis. **F.** Venn diagram illustrates an overlap between RBPs expressed in analyzed GBM-PRM, GBM-REC tissues, and HB control. **G.** Volcano plot of differentially expressed RBPs in GBM-PRM vs HB. The blue and red dots indicate downregulated and upregulated circRNAs, respectively.

To have a better insight into the role of differentially expressed circRNAs, we performed Gene Set Enrichment Analysis (GSEA) on their linear counterparts. The analysis revealed 22 GO terms with negative enrichment scores with most of them related to the neurotransmitter secretion, transport and regulation of their levels, signal release from synapse or ion transport. Although no enriched KEGG terms were found below adjusted p-value threshold (0.05), it is worth to note, that without multiple test correction among 13 enriched KEGG terms like ‘Neuroactive ligand-receptor interaction’, ‘Synaptic vesicle cycle’ and ‘Glutamatergic synapse’ were identified with negative enrichment scores (Suppl. Fig. 4).

Obtained results thus clearly indicate global deregulation of circRNAs in GBM, which may suggest their role in processes related to the development of this tumor type.

#### 2.1.2. Expression profile of RBPs transcript in GBM

As mentioned previously, RBPs play a pivotal role in circRNA biogenesis, thus they might be involved in regulating the balance between the expression of circRNAs and their linear counterparts. Therefore, we performed the analysis of the expression profile of RBP transcripts based on RNA-seq obtained from the same data set. We have identified 1420 RBP transcripts in HB, 1436 in GBM-PRM, and 1430 in GBM-REC samples (Fig. 1F). Differential gene expression (DGE) analysis of RBPs transcripts resulted in the finding that 817 from 1542 known human RBPs [30] are characterized by the significantly altered expression in GBM-PRM (469 up- and 348 downregulated) with a log2 fold change in the range between -4.8 and 4.7 (Fig. 1G, Suppl. Fig. 5).

### 2.2 Characteristics and validation of circRNAs dysregulated in GBM-PRM and GBM-REC

The DCE analysis revealed a list of differentially expressed circRNA among GBM-PRM and HB samples and indicated their possible involvement in processes related to GBM development. The potential progression markers were additionally identified by the comparative expression analysis between GBM-PRM and GBM-REC samples. 11 circRNAs were selected for further validation based on the list of mostly downregulated and upregulated molecules in GBM.

Based on the analysis of GBM-PRM versus HB control, we selected 4 downregulated circRNAs (circCADPS2, circEPB41L5, circUNC13C and circUSP45) and 4 upregulated ones (circARID1A, circGUSBP1, circPLOD2 and circVCAN). The DCE analysis between GBM-REC and GBM-PRM allowed us to distinguish 3 circRNAs specifically upregulated in recurrent samples: circEGFR, and 2 new, not previously annotated, circRNAs coming from *HLA-B* gene, and intergenic circRNA originating from chromosome 6p22.1 named for convenience as circInter6p22 (Suppl. Tab. 3).

Further comparison of circular to linear read ratio of selected circRNAs based on RNA-seq data, showed that in almost all cases the circular transcript is less abundant than the linear form in GBM, with the exception of VCAN circRNA transcript more expressed than its linear counterpart. Interestingly, we also perceived that the relation between circular and linear transcripts might differ between HB and GBM. For instance, circEPB41L5 is more abundant than its parental transcript in healthy brain, but this interplay is reversed in GBM where the linear form is more abundantly expressed (Fig. 2A-2B).

**Figure 2.**
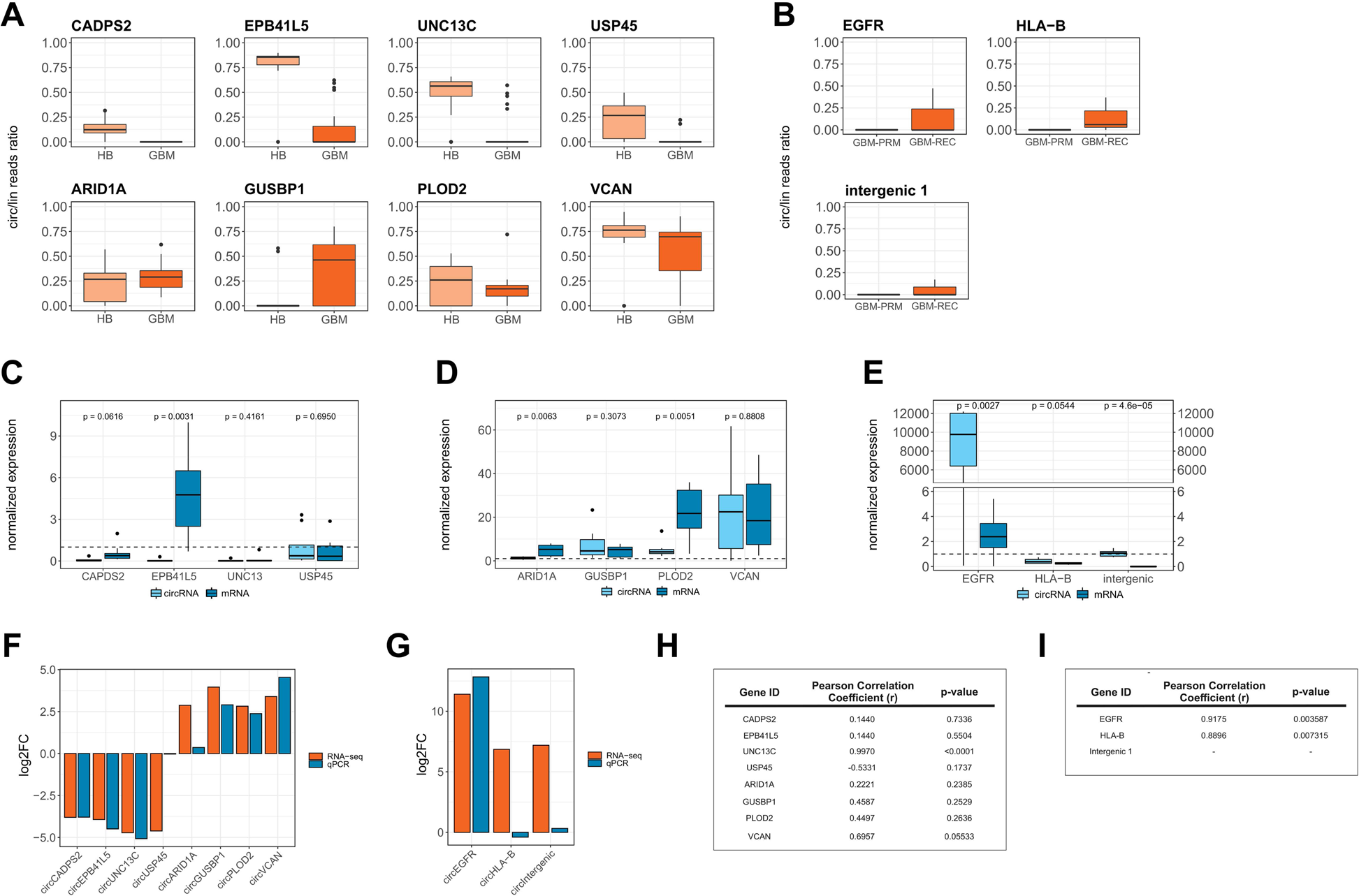
Expression of selected circRNA dysregulated in GBM and their linear counterparts. **A.** Boxplots illustrating the ratio of circular to linear RNA expression levels in RNA-seq analysis for analyzed transcripts dysregulated in GBM-PRM in comparison to HB. **B.** Boxplots illustrating the ratio of circular to linear RNA expression levels in RNA-seq analysis for analyzed transcripts dysregulated in GBM-REC in comparison to GBM-PRM. **C.** qRT-PCR results of circular and linear RNA expression levels of selected downregulated gene candidates in GBM-PRM normalized to HB control. Dashed horizontal lines on panels A-C indicatess HB expression level. **D.** qRT-PCR results of circular and linear RNA expression levels of selected upregulated gene candidates in GBM-PRM normalized to HB control. Dashed horizontal lines on panels A-C indicatess HB expression level. **E.** qRT-PCR results of circular and linear expression level of selected upregulated gene candidates in GBM-REC normalized to GBM-PRM. Dashed horizontal line indicates the expression level in GBM-PRM. **F.** Log2 fold change comparison of selected circRNAs dysregulated in GBM-PRM based on qRT-PCR and RNA-seq analysis. **G.** Log2 fold change comparison of selected circRNAs dysregulated in GBM-REC based on qRT-PCR and RNA-seq analysis. **H.** Pearson correlation of validated circRNAs and their linear counterparts dysregulated in GBM-PRM. **I.** Pearson correlation of validated circRNAs and their linear counterparts dysregulated in GBM-REC.

In order to confirm the circular structure of the identified molecules, the RNase R treatment was performed (Suppl. Fig. 6). The obtained data clearly support the RNase R resistance and high RNA stability of the verified circRNA molecules. Subsequently, we used the same 8 GBM-PRM samples that were utilized in RNA-seq, to validate the RNA-seq results with RT-qPCR. Overall, we confirmed previous results of selected upregulated and downregulated circRNAs. Regarding downregulated circRNAs, we observed simultaneous downregulation of their linear transcripts with one exception – circEPB41L5 in which the linear form was upregulated in GBM-PRM (Fig. 2C). Among all upregulated candidates VCAN was distinguished as the most upregulated one (Fig. 2D). Regarding circular RNAs upregulated in GBM-REC in comparison to GBM-PRM, we noted the upregulation of circRNAs expression in comparison to their linear counterparts, with a strikingly elevated level of circEGFR reaching ∼12000-fold change (Fig. 2E). Comparison of RNA-seq results with RT-qPCR supported obtained data in most cases (Fig. 2H). We also verified the correlation of the expression of circular and linear transcripts in analyzed samples and confirmed the significant positive correlation for UNC13C (r=0.997, p-value=<0,001), VCAN (r=0.6957, p-value=0.05533) in GBM-PRM vs. HB. Regarding circRNAs upregulated in GBM-REC we have also noticed high positive correlation between circular and linear isoforms for EGFR (r=0.9175, p-value=0.003587) and HLA-B (r=0.8896, p-value=0.007315) (Fig. 2I).

### 2.3 Interplay between circRNAs and RBPs

It is already known that RBPs regulate circRNAs biogenesis via binding to flanking introns and thus promote back-splicing process. They can also be sequestered by circRNAs acting as molecular sponges making them direct interactors in molecular processes. This interplay could be significant for the function of both circRNAs and RBPs, therefore we aimed to comprehensively investigate the interplay between those molecules [17, 31–34]

#### 2.3.1 RBPs might be sequestered by circRNAs

In the first step, we verified potential probability of circRNAs to sequester RBPs. To examine which differentially expressed RBPs are able to bind to deregulated circRNAs, we calculated k-mers (k=6) enrichment in differentially expressed circRNAs in GBM-PRM vs. HB (both up- and downregulated) and compared them to each binding motif’s position in RBPs. 6-mers enriched in differentially expressed circRNAs were mapped to the RBPs binding sites determined either by eCLIP or by RNA-bind and RNA-seq data [35]. As a result, we have obtained the list of 38 RBPs differentially expressed in GBM-PRM compared to HB tissue. We observed very distinct sequence patterns for RBP motifs enriched exclusively in downregulated vs upregulated circRNA. Namely, motifs enriched in downregulated circRNA were U-rich, whereas those for upregulated ones contained a lot of G and C nucleotides (Fig. 3, Suppl. Fig. 7). This observation indicated a distinct difference in the recognition sites of upregulated and downregulated circRNAs which might explain the discrimination mechanism between the binding of both types of molecules.

**Figure 3.**
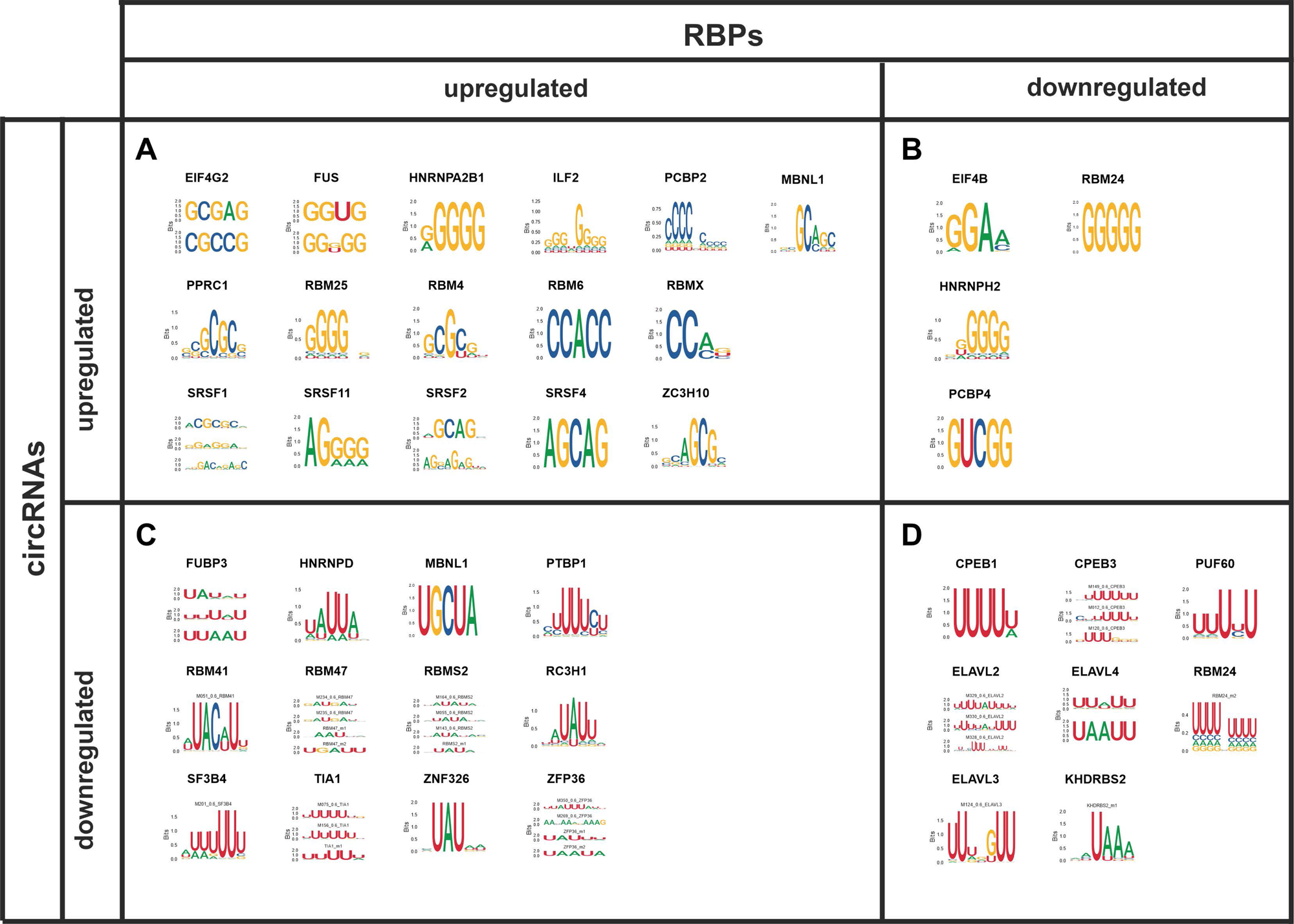
Differentially expressed RBPs with motifs enriched in differentially expressed circRNAs. **A.** Upregulated RBPs with motifs enriched in upregulated circRNAs. **B.** Downregulated RBPs with motifs enriched in upregulated circRNAs. **C.** Upregulated RBPs with motifs enriched in downregulated circRNAs. **D.** Downregulated RBPs with motifs enriched in downregulated circRNAs. Sequence logos represent motifs enriched in circRNAs.

#### 2.3.2 RBPs are correlated with circRNAs and their host gene expression

Next, we calculated correlation coefficients between the expression of previously identified differentially expressed RBPs and deregulated circRNA. We observed that expression levels of some RBPs significantly correlates with the expression levels of several hundred differentially expressed circRNAs (Fig. 4A, Suppl. Fig. 8). The vast majority exhibited a positive correlation, notably, the ELAV and CPEB families, which were downregulated in GBM (Fig. 4A, marked as red bars). Conversely, ZFP36, RBM47, PTBP1, RBM5S and SF3B4 mostly showed negative correlation with circRNAs (Fig. 4A, marked as green bars). Then, we assessed whether the expression of these RBPs correlates with expression of circRNAs’ linear counterparts. For this purpose, we used the subset of circRNAs parental mRNAs and calculated coefficients. Furthermore, we confirmed whether the correlation with RBPs have the same or opposite direction considering simultaneously circRNAs and mRNAs. Most of them showed the same direction of correlation in both circRNAs and corresponding mRNAs (Fig. 4B). Interestingly, RBPs positively correlated with circRNA expression (Fig. 4A marked as red bars) also showed positive correlation with mRNA expression in ∼50% of cases. In contrast, RBPs with expression negatively correlated with circRNAs (ZFP36, PTBP1, RBM47, RBMS2 and SF3B4) mostly did not show significant correlation with mRNA expression. Moreover, we investigated which of the deregulated RBPs might be associated with circRNAs subjected to validation. We observed that there are RBPs with altered expression levels which correlates with the expression level of selected circRNAs - regarding upregulated circARID1A and circGUSP1 there were over 20 positively correlated RBPs (Fig. 4C).

**Figure 4.**
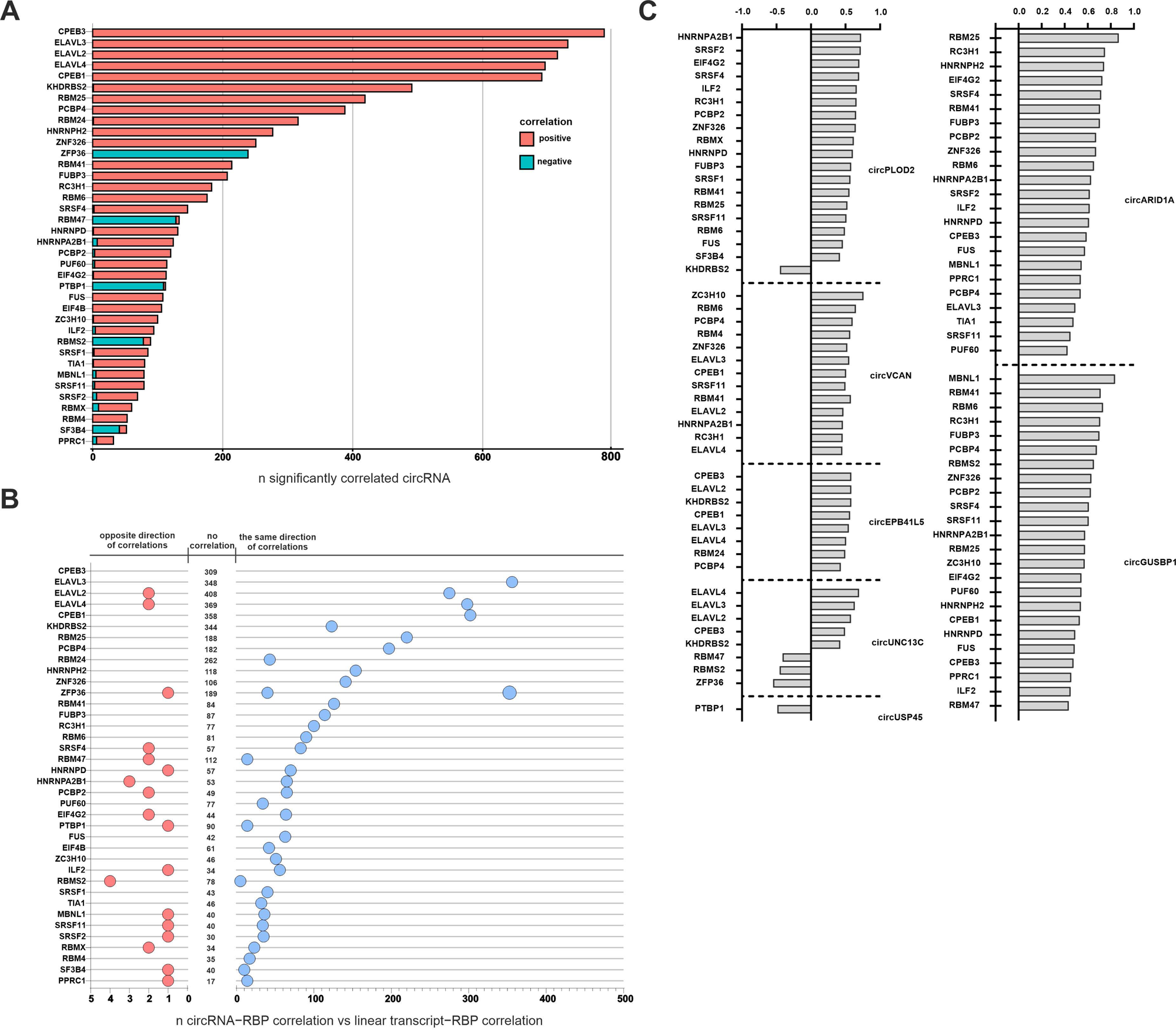
Expression of RBPs is correlated with circRNA and circRNA host genes expression. **A.** Counts of circRNA significantly correlated with differentially expressed RBPs. **B.** Comparison of correlation direction between RBPs (from figure A) and circRNAs vs RBPs and circRNAs linear counterparts. RBPs highlighted as blue show the same direction of correlation and RBPs highlighted as red show opposite direction. **C.** Correlation of the expression level of selected validated circRNAs with the expression level of dysregulated RBPs.

#### 2.3.3 RBPs might bind with flanking regions thus impact circRNAs biogenesis

As mentioned before, RBPs are important players in the circRNA biogenesis [17, 33, 34]. To identify the RBPs potentially involved in circRNAs biogenesis, we analyzed RBP 6-mers binding motifs enriched in flanking introns (up to 1000 nt upstream and downstream from the backsplice junction) of differentially expressed circRNAs in GBM vs. HB. We identified 14 differentially expressed RBP genes with binding motifs in the flanking introns across all the analyzed samples (Figure 5A). Correlation analysis between RBPs and circRNAs expression in GBM samples highlighted significant correlation with CPEB3, ELAVL2, ELAVL3, and KHDRBS2 showing a strong positive correlation with both up- and downregulated circRNAs - except one – ELAVL3 which correlated only with downregulated circRNAs. We also noted RBPs with a strong negative correlation with the expression level of upregulated circRNAs like IGF2BP2, and IGF2BP3. In search of potential circularization enhancers, we identified lowly expressed RBPs, which showed positive correlation with downregulated circRNAs (Fig. 5B-5C). CBEB3 was highly correlated with circRLF (rho = >0.8), circLRBA (rho = 0.7) and selected before as one of the most downregulated circRNAs in GBM - circEPB41L5 (rho = 0.65) (Fig. 5B), while KHDRBS2 showed strong positive correlation with circLRBA (rho >0.8) and circVMP1 (rho = 0.7). These findings might indicate RBPs exclusively involved in the regulation of circRNAs biogenesis.

**Figure 5.**
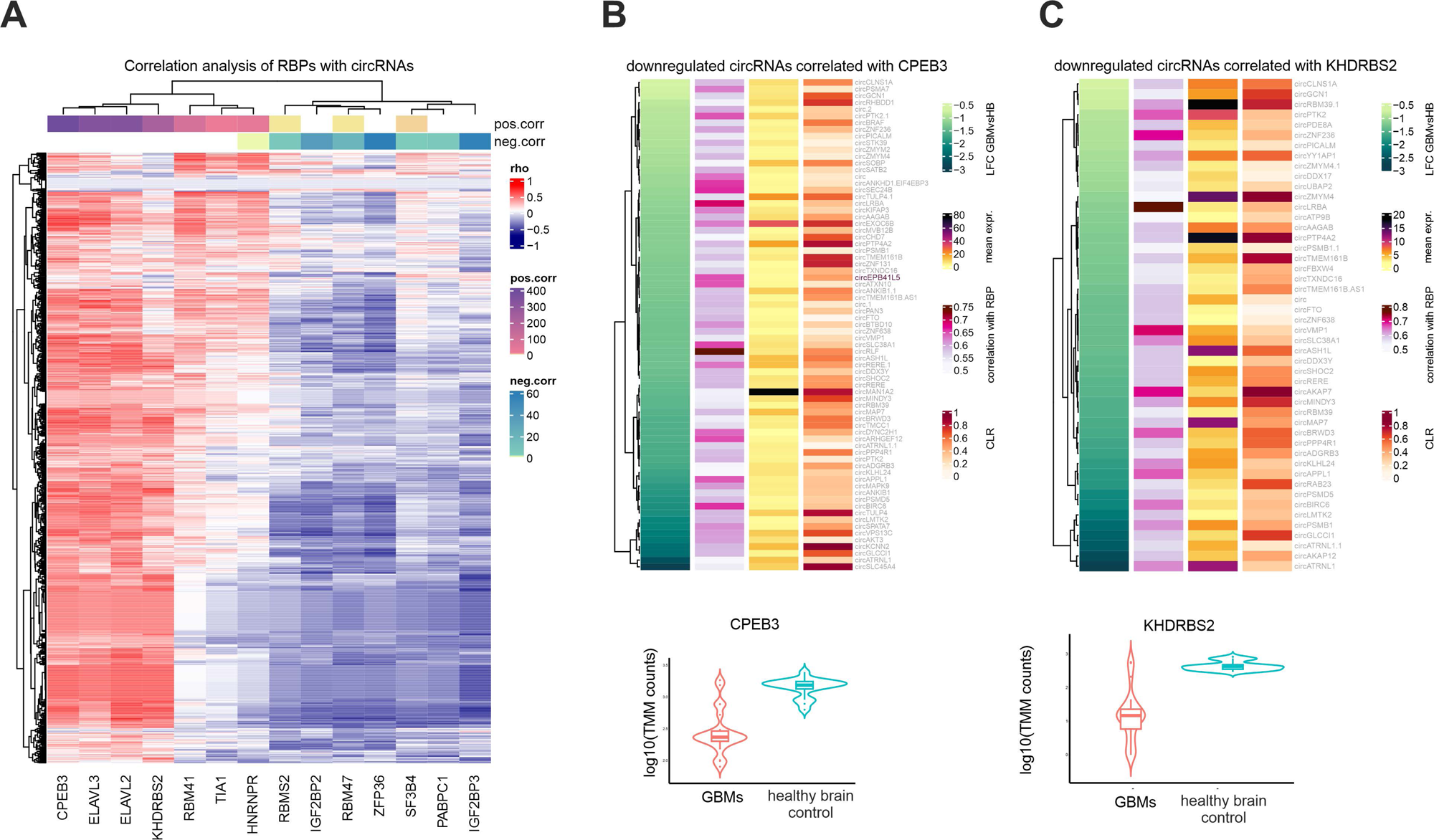
RBPs with motifs enriched in introns flanking circRNAs are correlated with circRNA expression. **A.** Heatmap of correlation (rho) between 14 RBPs with enriched motifs flanking backsplice junction of circRNAs differentially expressed in GBM vs HB. Only significant correlations (p values < 0.05) were reported. For each RBP number of positively and negatively correlated cicrRNAs were added in the upper bound of the heatmap. **B**-**C**. CBEP3 and KHDRBS2 normalized expression distribution in HB and GBM. For each RBP, correlated circRNAs are reported in terms of: I. average normalized expression in GBM samples; II. average circular to linear proportion (CLR) in GBM; III. log2fold change comparing circRNA expression in GBM vs HB and IV. correlation entity with the RBP - CBEP3 and KHDRBS2, respectively.

#### 2.3.4 RBPs could both impact circRNAs expression and be sequestered by them

We further investigated whether differentially expressed RBPs could both influence circRNAs biogenesis and be sequestered by the same circRNAs. We found a common set of 10 RBPs: CPEB3, ELAVL2, ELAVL3, KHDRBS2, RBM41, RBM47, RBMS2, SF3B4, TIA1, ZFP36 with motifs enriched in both - flanking introns and total sequence of circRNAs (Fig. 6A, Suppl. Fig. 9). Additionally, we calculated motif frequencies for these RBPs in the 100 nt regions encompassing the junction site (5’ boundary and 3’ boundary) and we defined intron-exon and exon-intron boundaries (Fig. 6B). For most RBPs, we observe similar motif frequencies in introns, boundaries and circRNA. However, an increased number of binding sites was observed in the sequence covering the 5’ boundary region for the CPEB3 protein. A similar dependence was noted in the case of the SF3B4, which was additionally characterized by an increased frequency of motifs in the 3’ boundary of the circRNA (Fig. 6C). Altogether, it suggests that RBPs might play a dual role-both influence circRNAs biogenesis and at the same time be regulated by these molecules.

**Figure 6.**
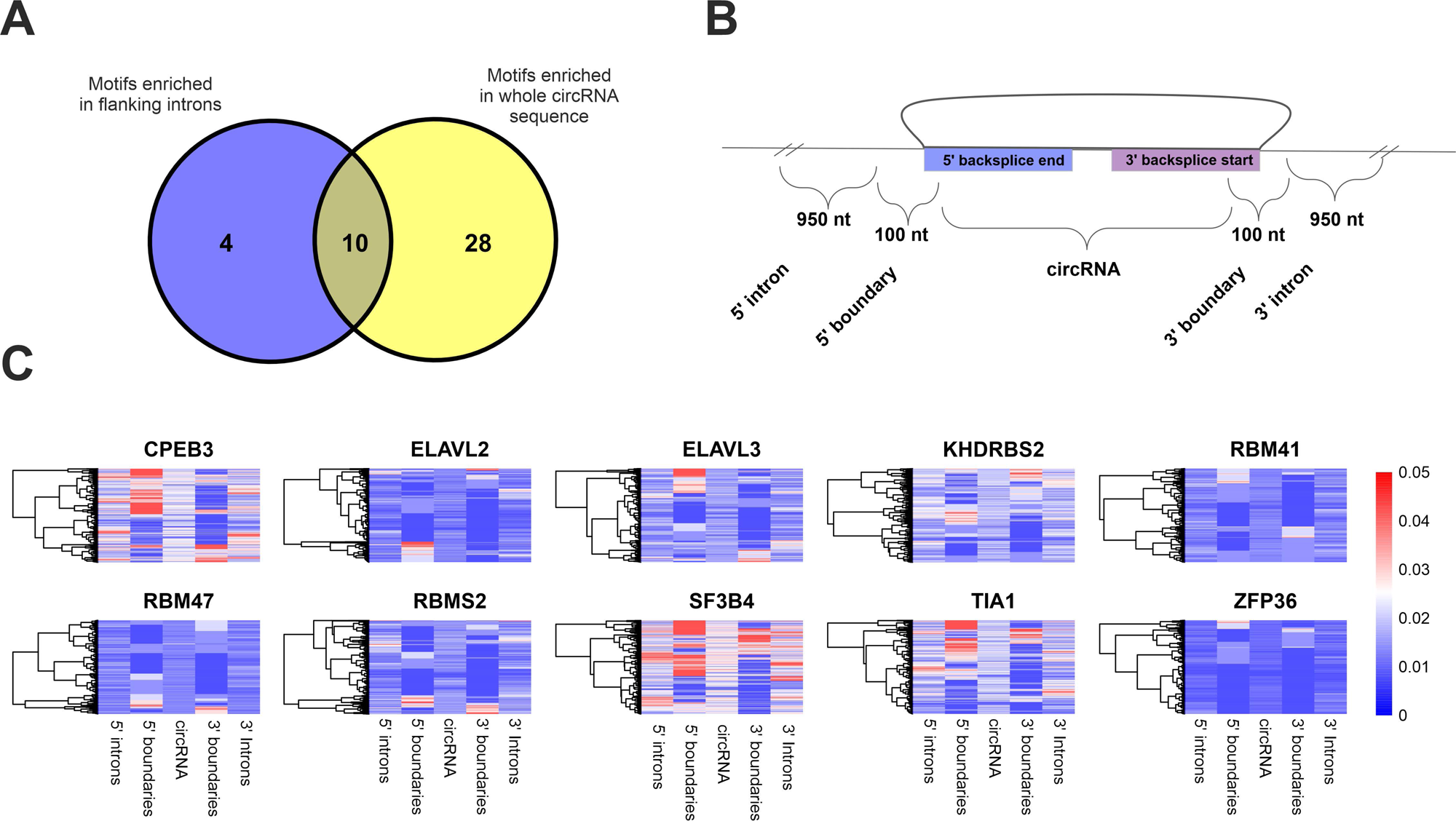
RBP motifs enriched in flanking introns and circRNA itself. **A.** Venn plot showing overlap between RBPs with motifs enriched in flanking introns and in total sequence of circRNA. **B.** Schematic representation of circRNA with 5’ and 3’ flanking introns (1000 nt upstream from the 5’ backsplice junction end, 1000 nt downstream from the 3’ backsplice junction end) and 5’ and 3’ boundaries (50 nt from intron and 50 nt from circRNA sequence). **C.** Normalized frequency of enriched motifs in RBPs within regions shown in panel B.

### 2.4 Combinatorial analysis of RBPs and circRNAs expression profiles suggest new patient stratification criteria

#### 2.4.1 CircRNA expression is related to the GBM subtypes

We utilized the list of 840 genes from the TCGA GBM dataset and literature [6] as it provides the knowledge for unifying GBM transcriptomic and genomic data allowing for GBM molecular stratification into 4 molecular subtypes: classical, mesenchymal, preneural and neural. The provided framework allowed us to cluster the analyzed 23 GBM study samples into classical (5 samples), mesenchymal (8), neural (5), and proneural (5) (Fig. 7A). Then we determined the presence of different circRNAs in each of the individual molecular subtypes (Fig. 7B). We identified 54 differentially expressed circRNAs in the neural subtype versus other subtypes, where circERC1 and circMLIP were characterized by the highest expression levels (Fig. 7C, Suppl. Tab. 4). The data show the high similarity of the neural subtype to the HB samples. Interestingly, we further distinguished overexpressed circCOL4A1 and circCOL1A2 which were upregulated in the mesenchymal subtype (Suppl. Tab. 5). Moreover, in this subtype, we observed decreased expression of circRBM39 and circRIMS2. Noteworthy, we did not find any circRNAs related exclusively to the classical and proneural subtypes.

**Figure 7.**
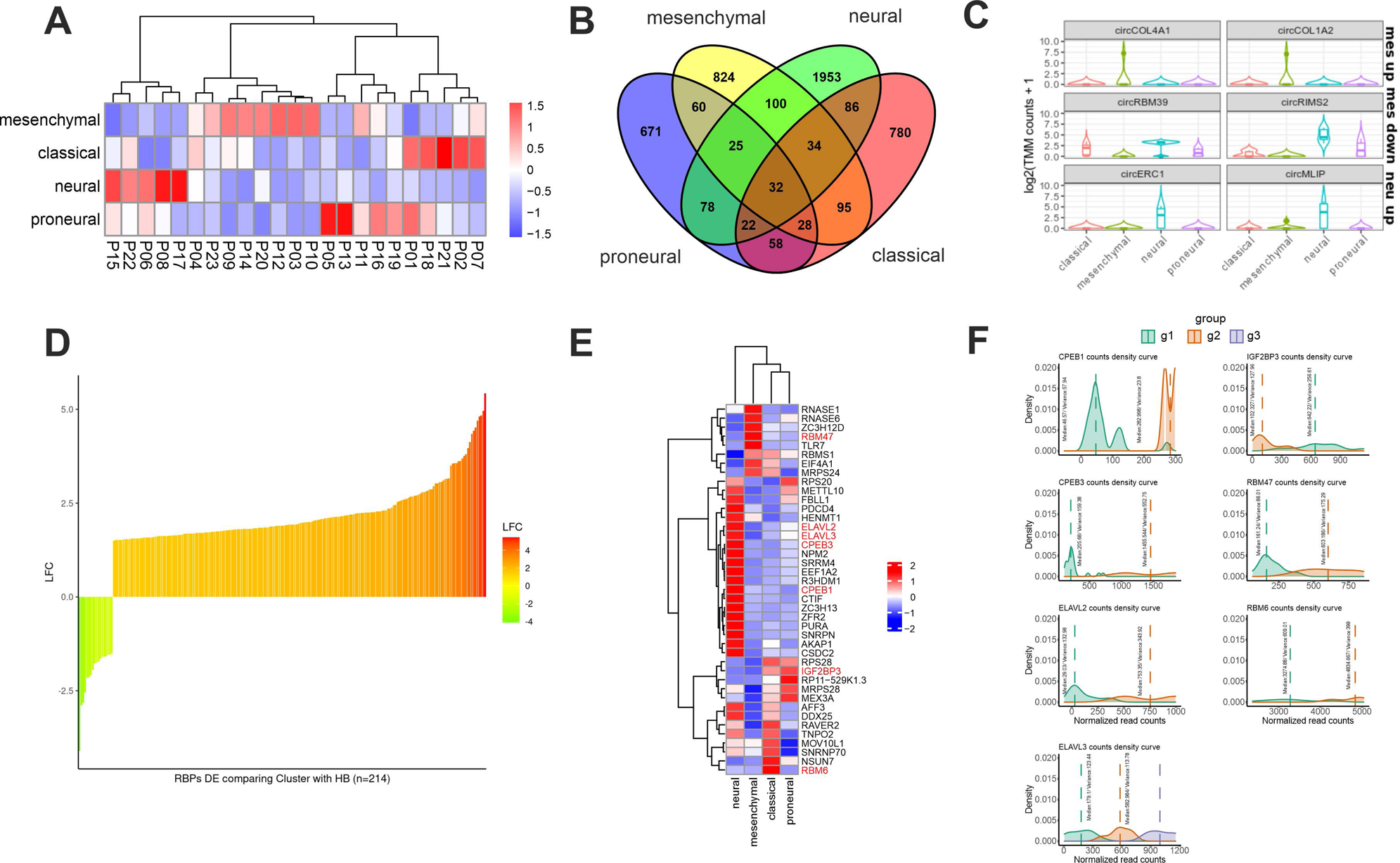
Representation of a subtype-specific circRNA and RBPSs clustering GBM into new subtypes. A. Heatmap presenting subtype classification of the GBM samples used in the study. B. Venn diagram shows the number of the identified circRNAs in detected GBM subtypes. C. Comparison of selected circRNA up- and downregulated in mesenchymal and upregulated in neural subtypes. D. Log2 fold changes of 214 differentially expressed RBPs that can be used for patients’ stratification. E. Heatmap showing mean expression differences between 41 RBPs within samples belonging to distinct GBM subtypes. RBPs highlighted red contain binding motifs enriched in differentially expressed circRNA sequences/flanking regions. F. RBPs clustering GBM samples into 2 groups with motifs enriched in differentially expressed circRNA sequences/flanking regions. G. ELAVL3 clusters GBM samples into 3 groups and has binding motifs enriched in differentially expressed circRNA sequences/flanking regions.

#### 2.4.2 RBPs in GBM can be used as the new potential stratification markers

To assess also the clinical value of RBPs in GBM we aimed to identify RBP-based signatures for patient stratification. At first, we subset RBPs able to discretize patients into two clusters according to their expression with no significant association with GBM molecular subtypes. We identified RBPs that cluster GBM samples into two or more groups (Suppl. Tab. 6). Next, we focused on RBPs whose expression in at least one of these newly defined clusters was differentially expressed in comparison to HB samples. We observed a prevalent upregulation of RBPs in the novel subgroups of patients (Fig. 7D). The overall expression of these RBPs was able to separate GBMs from HB samples, with the clusterization of patients in two major newly identified subgroups, independent from the GBM molecular subtypes. We were also looking for RBPs potentially related to the previously mentioned known 4 molecular subtypes and established that there are 41 of them (Suppl. Fig. 5). From these RBPs, we found 7 with motifs enriched in differentially expressed circRNA and/or flanking sequences (CPEB1, CPEB3, ELAVL2, ELAVL3, IGF2BP3, RBM47, RBM6) (Fig. 7E, highlighted in red). RBM47 was highly expressed in mesenchymal, RBM6 in classical, and IGF2BP3 in classical and proneural subtypes (Fig. 7E). Majority (6) of 7 selected RBPs further cluster GBM samples into 2 newly defined groups (Fig. 7F) and ELAVL3 cluster GBM samples into 3 groups (Fig. 7G). Thus, RBPs expression profile might be taken into consideration as an additional factor stratifying individual GBM tumors, which could enrich the current division.

#### 2.4.3 RBPs expression correlates with the overall survival rates of GBM patients

Finally, we checked the differentially expressed RBPs and their correlation with the overall survival rates (OVS) of GBM patients. We utilized available data from TCGA GBM dataset. We selected patients with high or low expression of the given RBPs and found that 30 RBPs differentially expressed in our 23 GBM samples were significantly correlated with OVS in the TCGA GBM cohort (Suppl. Fig. 10A - upregulated RBPs, 10B - downregulated ones). 22 of them were previously shown to be brain tumor markers, correlated with disease outcome or response to the treatment (Suppl. Tab. 7).

## 3. Discussion and future perspectives

In our study, we conducted an extensive RNA-seq screen focusing on circRNA and RBP expression profiles as well as their interactions. We detected circRNAs differentially expressed in GBM primary tumors, identified the circRNA progression markers in GBM recurrent ones, as well as we established the expression profile of RBP transcripts. Further analysis enabled us to generate a comprehensive catalog of circRNA-RBP interactions including the sequestration of RBPs by circRNA and their involvement in circRNA biogenesis. Additionally, we demonstrated also the clinical potential of circRNAs and RBPs in GBM and identified them as the stratification markers in the *de novo* assembled tumor subtypes.

Given that GBM is an aggressive, therapy-resistant brain tumor with high inter- and intra-tumoral diversity, there is an urgent need to understand the transcriptome profile of this tumor type to find new, distinctive molecular biomarkers and therapeutic targets for GBM. Due to their high stability and brain-specific expression pattern, circRNAs are considered remarkably interesting as potential cancer molecular signatures [36–38]. Additionally, their role in cancer-related processes namely proliferation, migration, invasion, and apoptosis in GBM have already been studied and verified [39]. Most of the studies concerning circRNA in GBM are focused on a single or a few molecules and their specific mechanism of action with only a few studies attended to high-throughput circRNA profiling in GBM [40–43]. One study reported up to now the GBM patients–derived data being however focused on general cancer-related mechanisms. The findings reveal that certain circRNAs are deregulated across multiple types of cancers, including GBM, esophageal squamous cell carcinoma, lung adenocarcinoma, thyroid cancer, colorectal cancer, gastric cancer, and hepatocellular carcinoma. Despite these observations, the study does not solely focus on GBM, nor validates the obtained results. Additionally, it did not provide any description of the complex network of interactions and dependencies between circRNAs, their linear counterparts, and RBPs in glioblastoma [44].

In our study, we identified 1270 circRNAs differentially expressed in GBM-PRM samples compared to HB controls with almost 90% being downregulated. This pattern aligns with previously published circRNA profiles in different cancer types, including GBM, indicating a potential competition in biogenesis between circRNAs and mRNAs utilized for the protein synthesis - essential for proliferating tumor cells [43, 45–48]. Further, common downregulation of circRNAs observed in cancer cells can be caused by their extensive proliferation. This could dilute the concentration of stable circRNAs as previously suggested [45, 49]. It is also supported by the opposite phenomena, namely the accumulation of circRNA in the non-proliferating aging mouse brain, mainly composed of post-mitotic cells [50]. Similar trends can be observed in *D. melanogaster* and *C. elegans* aging models [51, 52]. On the contrary, in T-cell acute lymphoblastic leukemia (T-ALL) the majority of deregulated circRNAs showed increased expression in comparison to normal thymocytes [53]. Tumor cells display a high rate of transcription, especially in aggressive cancers like GBM, while the increased incidence of back-splicing happens rather when co-transcriptional processing activities are inhibited or slowed down [54, 55]. This agrees with our analysis, where circRNA downregulation is in most cases independent of changes in their linear counterparts which suggests no impairment in the transcription process but rather the involvement of back-splicing.

Prior research has demonstrated that dysregulation of circRNAs might exert oncogenic functions in GBM, both downregulated in high-grade glioma circBRAF and upregulated in GBM circPITX1 are associated with poor patients’ prognosis [43, 56]. In our study, among downregulated circRNAs, circEBP41L5 was detected as the most decreased one in GBM and displays the biggest discrepancy between the expression of linear and circular transcripts. In the literature, circEBP41L5 is described as a GBM suppressor wthat acts through miR-19a sponging, thus it could serve as a prognostic or therapeutic molecule for new clinical approaches [57]. Remaining downregulated circRNAs might act as suppressors and require further studies to better understand the mechanism underlying GBM.

In the remaining 10% of upregulated circRNA in GBM, the most upregulated appeared to be circVCAN which high expression is observed also in gastric cancer [58] and radioresistant glioma tissues [59]. The knockdown of these circRNAs resulted in the inhibition of cell proliferation, migration, and invasion, and accelerated apoptosis by regulating miR-587 [58] and miR-1183 [59]. Another validated differentially expressed circRNA with elevated expression levels in our GBM tissues – circPLOD2 was also found upregulated in GBM in a previous study [60]. This finding indicates an immense potential of circPLOD2 as a biomarker of GBM. CircPLOD2 has also been described as a promoter of tumorigenesis and recurrence biomarker in colon cancer. [61, 62]. It is also frequently upregulated under hypoxia conditions in HeLa and MCF-7 cancer cell lines which mimic oxygen-deprived core of tumor mass [63]. There are no literature reports on selected circARID1A and circGUSPB1 which are overexpressed in GBM. The mRNA of *ARID1A* gene emerged as a cancer suppressor in different cancers. The absence of ARID1A in cancer can lead to widespread dysregulation of gene expression in cancer initiation, promotion, and progression [64, 65]. We also tried to distinguish potential molecular markers of GBM progression based on our analysis. 3 circRNAs: circEGFR, circHLA-B, circInter6p22 were found differentially expressed between GBM-PRM and GBM-REC tissue samples. The most overexpressed candidate, particularly in GBM-REC tumors, is circEGFR. CircEGFR is derived from the EGFR gene which is a well-established oncogene in various cancers [66, 67]. In contrast, circEFGR was also found as an inhibitor of the malignant progression of glioma by regulating the levels of miR-183-5p and TUSC2 [68]. Since there are different circEGFR isoforms originating from different genomic locations, the isoform defined in the mentioned report (hsa_circ_0080223) shows downregulation in tumor tissues, however circEGFR regarding our research (hsa_circ_0080229) is upregulated. Another study, consistent with our findings, indicates that hsa_circ_0080229 upregulates the expression of murine double minute-2 (MDM2) and promotes glioma tumorigenesis and invasion via the miR-1827 sponging mechanism [69]. This proposes that the expression of circEGFR is increasing with tumor progression. Our results, although observed in a depleted group of samples and should be interpreted with caution, supported this hypothesis and it could be intriguing to further examine additional glioma tissues of different histopathological grades in the future.

Previous studies revealed that the expression pattern of circRNAs and their circularization are significantly affected by RBPs which bind to the specific motifs in pre-mRNA [17, 34]. We observed significant changes in the expression of ∼half of annotated human RBPome in GBM, which suggests that some of the changes in circRNA expression may be caused by disruption of their biogenesis. Moreover, these direct interactions of circRNAs with RBPs can substantially affect molecular processes [70]. Additionally, circRNAs can act as protein sponges, decoys that regulate protein accessibility in cell compartments, and also as scaffolds that allow protein-protein interactions mediating both circRNA and RBP functions [33]. Based on these findings, we decided to analyze the transcriptome of RBPs which could potentially interact with our deregulated circRNAs. In our data, recognition motifs for 38 RBPs are enriched in differentially expressed circRNAs. In our study, upregulated circRNAs are mostly motifs enriched in G and C in differentially expressed RBPs. It was shown that RBPs enriched in G-rich motifs might be splicing activators, that might potentially explain the upregulation of circRNAs to which they bind [25]. On the other hand, downregulated circRNAs are in U-rich motifs. It was previously shown that U-rich motifs correlate with higher mRNA decay rates [71]. Further analyses showed that ELAV families, downregulated in GBM, are correlated with the highest number of downregulated circRNAs. ELAVL proteins recognize AU-rich elements in the 3’ UTRs of gene transcripts and thereby regulate gene expression post-transcriptionally stabilizing mRNA to avoid degradation [72]. It was also shown that circularizing exons were significantly more enriched with RBP binding sites compared to non-circularizing exons [17]. In our study, we found 14 RBP binding motifs enriched in flanking introns of differentially expressed circRNAs with the most promising HNRNPD and ZFP36 significantly correlated with highly expressed circRNAs. In our study, ELAVL2 and ELAVL3 showed a positive correlation with circRNAs enriched in flanking introns with the highest frequency in a region covering 100 nt upstream from the 5’ backsplice junction end. These observations indicate specific circRNA-RBP interactions that might be directly involved in the circRNA processing.

Another protein family that might impact circRNAs expression is CPEB. Both CPEB1 and CPEB3 are downregulated in GBM with U-rich motifs enriched in downregulated circRNAs and show the highest number of positive correlated downregulated circRNAs including validated by us circEPB41L5. We showed, that KHDRBS2 predicted to be involved in alternative splicing [73] is the most downregulated RBPs in GBM and highly correlated with exosome-transmitted circVMP1 which facilitates the progression and cisplatin resistance of non-small cell lung cancer [74] and promotes glycolysis and disease progression in colorectal cancer [75].

Besides the transcriptome-wide characterization of circRNAs and RBPs in GBM and their putative interactions, we extend our analysis to determine differentially expressed circRNAs and RBPs specific for known molecular GBM subtypes. GBM is a heterogeneous disease that can be classified into four known molecular subtypes according to mutation landscape and gene expression pattern [6].

According to the circRNA expression pattern, the neural subtype was the most similar to the healthy brain samples. The RBPs expression analysis revealed the high expression levels of CPEB and ELAVL families in neural GBM samples which are generally downregulated in GBM ([76, 77]. These findings confirm earlier reports that the neural subtype may be non-tumor margins contamination [78–80].

Among GBM subtypes, the mesenchymal one is known as the most aggressive, invasive, and resistant to treatment [81]. We found further that both circCOL4A1 and circCOL1A2 are highly expressed in this subtype. Moreover, circCOL1A2 was previously described as upregulated in gastric cancer enhancing the migration and invasion properties [82] and also promoting angiogenesis in diabetic retinopathy [83, 84], which can make circCOL1A2 a relevant marker of mesenchymal GBM subtype. For the first time, we managed to distinguish new GBM groups based on RBP expression patterns, independent of known molecular subtypes. As mentioned before, potentially involved in circRNAs biogenesis CPEB and ELAVL families stratify GBM into 2 or 3 groups. Altogether, both CPEB and ELAVL families appear to have an impact on the circRNAs.

In conclusion, we provided a comprehensive global-scale analysis of circRNAs and RBPs in glioblastoma based on the patient-derived samples.

Specifically, we established a list of circRNAs differentially expressed in primary GBM tumors, the circRNA progression markers in recurrent GBM samples, but also, indicated a global deregulation of RBP transcripts. Selected circRNAs were further verified experimentally by the qRT-PCR method. Subsequent analysis allowed us to generate a comprehensive catalog of circRNA-RBP interactions regarding both the RBPs sequestration by circRNA as well as the RBPs involvement in circRNA biogenesis. Furthermore, we demonstrated the clinical potential of circRNAs and RBPs in GBM and proposed them as the stratification markers in the de novo assembled tumor subtypes. Moreover, our findings suggest that both circRNAs and RBPs might be considered as clinical markers and tumor-subtyping factors in the future. The list of potential functions of distinct circRNAs as well as the circRNA-RBP interactions is long and their regulatory functions are complex. The mechanism of action and the mutual dependencies await still further in-depth investigation. However, with our comprehensive study, we provide a foundation for future research into the molecular mechanisms and clinical implications of circRNAs and RBPs interactions in glioblastoma.

## 4. Materials and Methods

### 4.1 Patients’ sample collection

Tumor tissues (n=26; Suppl. Tab. 1) were collected from GBM patients up to one hour after tumor excision. The material was obtained from the Department and Clinic of Neurosurgery and Neurotraumatology of the University of Medical Sciences in Poznan (Poland) and from the Department of Neurosurgery of Multidisciplinary City Hospital in Poznan (Poland). Prior to the surgery, the approval by the Bioethics Council of the Poznan University of Medical Science (Nr. 46/13) and the donors’ consent had been obtained. Four commercial samples (purchased from Ambion, Clontech and Takara companies) of pooled human brain (HB) total RNAs were used as healthy brain controls (Suppl. Tab. 2).

### 4.2 RNA extraction and quality check

Total RNA extraction from GBM patient tissue was performed using TRIzol reagent (Invitrogen) according to the manufacturer’s protocol. RNA samples have been subjected to DNase I treatment using a ready-to-use DNA-free™ DNA Removal Kit reagents following the manufacturer’s protocol (Ambion). The quality of purified RNA was measured by NanoDrop 2000 spectrophotometer (Thermo Fisher Scientific) followed by agarose gel electrophoresis. RNA integrity was verified on Agilent Bioanalyzer 2100 (Agilent Technologies). RNA samples with RNA Integrity Number (RIN) at least 7.4 were used for library preparation and RNA-seq analysis (Suppl. Fig. 11, 12).

### 4.3 Library preparation and RNA sequencing

300 ng of total RNA were ribosomal RNA-depleted using RNase H [23]. Ribosomal RNA-depleted libraries were constructed using Illumina’s TruSeq Total RNA Library Prep Kit. Adaptor ligations were performed according to manufacturer’s instructions and cDNA fragments were amplified in RT-qPCR for 8-15 cycles. After the purification with AMPure XP beads, the DNA concentration was measured with Qubit (Thermo Fisher Scientific) and the fragments length were defined with Screen Tape Assay Agilent D1000, 4200 Tape Station System. RNA sequencing was performed using Illumina Hi-seq 4000, average of ∼73 million of 150 paired-end reads per sample.

### 4.4 circRNAs identification

Quality control of raw sequencing reads was done with FastQC (https://www.bioinformatics.babraham.ac.uk/projects/fastqc/). Next, the adapters were removed using Trimmomatic [85], version 0.38 with the following parameters ILLUMINACLIP:2:30:10 SLIDINGWINDOW:10:25 MINLEN:35. After adapter trimming, a second quality check with FastQC was performed. A genomic index was created for GRCh37 human genome obtained from Gencode using Burrows-Wheeler aligner [86], version 0.7.17 (bwa –bwtsw). Then, reads were mapped with the following parameters: bwa-mem –T 19. CircRNAs were detected from alignment files with CIRI [87], version 2.0.6 using default parameters. CircRNAs annotation and differential expression analysis was performed using circMeta R package [88]. The edgeR [89] method was used for differential expression (DE) analysis. Heatmaps were prepared using the pheatmap R package and data normalized using Trimmede Mean of M-values (TMM) method from edgeR R package [89].

Gene Set Enrichment Analysis (GSEA), including Gene Onthology (GO) and Kyoto Encyclopedia of Genes & Genomes (KEGG) was performed using clusterProfiler R package [90] and IDs for linear transcripts ranked by circRNA log2 fold change between GBM and HB samples. In case of multiple circRNAs originating from one gene, mean log2 fold change was used. For both GO and KEGG, the following set of parameters was used: nPerm = 10000, minGSSize = 3, maxGSSize = 800, pvalueCutoff = 0.05, pAdjustMethod = “fdr”.

Data were visualized using the ggplot2 R package. Most of the analysis was done in R 4.1.3, except for KEGG enrichment for GSEA, which was prepared in R 3.6.1.

### 4.5 RBPs identification

To identify RBPs differentially expressed between HB and GBM samples, trimmed reads were mapped to the human genome (GRCh37 v30 from Gencode) using Burrows-Wheeler aligner [86], version 0.7.17, followed by mapping sequencing reads to genomic features using featureCounts function from Rsubread package with useMetaFeatures option and without counting multimapping and multi-overlapping reads. Differential gene expression analysis was performed using edgeR glmQLFTest [89]. RBPs were selected from differentially expressed genes based on the human RBP list, obtained from Gerstberger et al., 2014 [30].

### 4.6 circRNAs-RBPs interactions

To identify putative RBPs binding sites in differentially expressed circRNA, first, circRNA sequences were extracted using FcircSEC [91] followed by k-mer enrichment analysis between differentially expressed and non-differentially expressed circRNAs. K-mer enrichment analysis was performed with k = 6, using kmer_compare function from the FeatureReachR package. Then 6-mers enriched in differentially expressed circRNAs were mapped to the RBPs binding sites determined either by eCLIP [35] or by RNA bind and Seq experiments [25], using estimate_motif_from_kmer function from FeatureReachR and CISBPRNA_hs or RBNS position weighted matrices lists, respectively. Significantly enriched RBP binding motifs (adjusted p value <0.1) were visualized using ggseqlogo R package [92]. Analysis was then repeated for circRNA flanking regions (1000 nt flanks).

Correlation of expression of RBPs, which were differentially expressed between HB and GBM samples, and their binding motifs enriched in differentially expressed circRNA, was estimated with differentially expressed circRNA expression using cor.test function from R with Spearman rank correlation option. The Upset plot was prepared using the UpSetR package [93].

### 4.7 GBM subtypes analysis based on circRNAs

Raw sequencing data were subjected to the quality check using FastQC tool. The adaptors were trimmed and the reads were filtered to eliminate the low-sequencing-quality bases using Trimmomatic [85]. Furthermore, the RNA-seq reads were mapped to the human reference transcriptome (ENSEMBLE V.102) to quantify the expression level of the transcripts. The transcript-level estimates were summarized and associated with the gene IDs for gene-level analysis using tximport [94]. For the samples categorization we used genes indicated in TCGA GBM dataset and other genes that have been found to be significant for GBM molecular subtyping including *SLC12A5*, *SYT1*, *GABRA1*, *NEFL*, *CDKN1A*, *NF1*, *MET*, *PDGFRA*, *BOP1*, *ILR4* were included. CircRNAs differentially expressed within the subtypes were identified using circMeta R package [88] with edgeR [89] method for differential circRNAs expression (DCE).

### 4.8 GBM samples stratification based on RBPs expression

GBM samples stratification according to RBPs expression changes was performed using the CircIMPACT workflow [95]. At first, we subset RBPs able to discretize patients into two clusters according to their expression with no significant association with GBM molecular subtypes (χ² test, p-value > 0.01). We clustered patients in novel subgroups linked by common RBPs pattern expression. The unsupervised clustering was performed using the k-means algorithm with euclidean distance and automatic selection of the optimal number of k clusters of patients using the silhouette index. Finally, we tested which RBPs reach significant estimates of the expression changes between previously defined clusters (one-way ANOVA test p-values ≤ 0.05). After correction for multiple tests, the RBPs with significant expression variation were used to compose the list of discriminant RBPs for the corresponding sample clustering.

### 4.9 Survival analysis

Survival analysis was performed on the GBM dataset from TCGA. Log2 transformed HTseq counts were downloaded using UCSC Xena tool [96]. Log2 transformation was reversed and raw reads were normalized using TMM method from edgeR R library [89]. Then differentially expressed RBPs were selected (listed in Suppl. Tab. 8) and for each RBP quartile normalization was applied. Samples belonging to Q1 were marked as ‘low’, Q2 and Q3 - ‘medium’ and Q4 - ‘high’. Survival analysis was performed using survival and survminer R libraries between samples with ‘low’ and ‘high’ expression levels. Log-rank (Mantel-Haenszel) test was used to assess statistical significance (p value < 0.05). Data were visualized using ggsurvplot function from survminer library.

### 4.10 Reverse transcription and quantitative RT-qPCR

The reverse transcription reaction was proceeded using the Transcriptor High Fidelity cDNA Synthesis Kit (Roche) according to the manufacturer’s protocol. 500 ng of total RNA extracted from patient-derived tissues was used. The real-time qPCR reaction was performed using the CFX Connect Real-Time PCR Detection System (Bio-Rad) on the 96-well plates recommended by the manufacturer. Each sample was run in three technical replicates. RT-qPCR analysis was performed with LightCycler® 480 SYBR Green I Master (Roche) and primers designed using the Primer-BLAST tool [97] and purchased from Thermo Fisher Scientific. Primers’ sequences are listed in the Suppl. Tab. 9. The expression level of the studied genes was calculated inCFX Maestro 2.0 software. Hypoxanthine phosphoribosyltransferase (*HPRT*) was used as an endogenous control for calculating the expression levels of selected genes. As control, commercially available RNA from HB tissues was used (Suppl. Tab. 2).

### 4.11 RNase R treatment

2 μg of total RNA were treated with 4U of RNase R (Lucigen) at 37°C for 5 or 10 minutes in case of downregulated and at 37°C for 30 minutes for upregulated circRNAs followed by 20 minutes at 65°C. Next, total RNA was purified using NucAway Spin Columns (Invitrogen) and reverse transcribed with the Transcriptor High Fidelity cDNA Synthesis Kit (Roche) according to the manufacturer’s protocol. Then, PCR was performed with 35 cycles (15s denaturation at 95°C followed by 30s at the optimum annealing temperature for each pair of primers (Suppl. Tab. 9) and 20s extension) with a prior 3-minute denaturation. The depletion of linear RNA was confirmed by running the PCR products on agarose gel.

### 4.12 Statistical analysis of the results

Each experiment was performed in three biological replicates including three technical replicates each. Statistical significance of the differences observed in experimental data was calculated in R, using t-test (stat_compare_means function). P-values are presented in plots.

## Acknowledgements

We would like to thank Neelanjan Mukherjee, Monika Piwecka, and members of Rolle and Rajewsky lab for critical review of the manuscript.

## 5. Authors Contribution

Conceptualization, JLŁ, ZZ, MPS, KK. JOM, AG and KR; Sample collection, RP, and AMB; Formal analysis, JLŁ, ZZ, MPS, JOM, AG, KK, AB, MZ, JK, AR-W and KR; Investigation, JLŁ, ZZ, MPS, KK, AG, AB and KR; Methodology, JLŁ, ZZ, MPS, AB, SB, AR-W and MZ; Resources, KR, NR; Supervision, KR; Funding acquisition, KR; Visualization, MPS, AB; Writing - original draft, JLŁ, ZZ, MPS, JOM; Writing - review & editing, JLŁ, ZZ, MPS, JOM, AG, KK, AB, SB, AR-W, and KR. All authors have read and approved the final version of the manuscript.

## 6. Conflict of Interest

The authors declare no conflict of interest.

## 7. Funding

Żaneta Zarębska, Konrad Kuczyński, Julia O. Misiorek and Katarzyna Rolle were supported by the Polish National Science Centre grant (NCN; 2017/25/B/NZ3/02173). Julia Latowska-Łysiak, Marcin P. Sajek, Adriana Grabowska, Julia O. Misiorek and Katarzyna Rolle were supported by the Polish National Science Centre (NCN; 2017/26/E/NZ3/01004^)^ Konrad Kuczyński was supported by the NCBR program (POWR.03.02.00-00-I032/16). Marcin P. Sajek was supported by the Polish National Agency for Academic Exchange Bekker program (PPN/BEK/2019/1/00173).

## 8. Ethics approval and consent to participate

The GBM samples (*n* = 26) were obtained from the Clinic of Neurosurgery and Neurotraumatology, Karol Marcinkowski University of Medical Sciences in Poznan, Poland, and Department of Neurosurgery, Jozef Strus Hospital in Poznan, Poland during 2015-2020 based on the approval from the Ethical Committee (Nr. 46/13), and individuals signed an informed consent form.

## 9. Data availability

Sequencing data have been deposited in NCBI GEO repository under accession code GSE196695. Scripts used for data analysis and processed data are available at https://github.com/MSajek/GBM_circs.

## Supplementary tables legends

**Supplementary Table 1.**
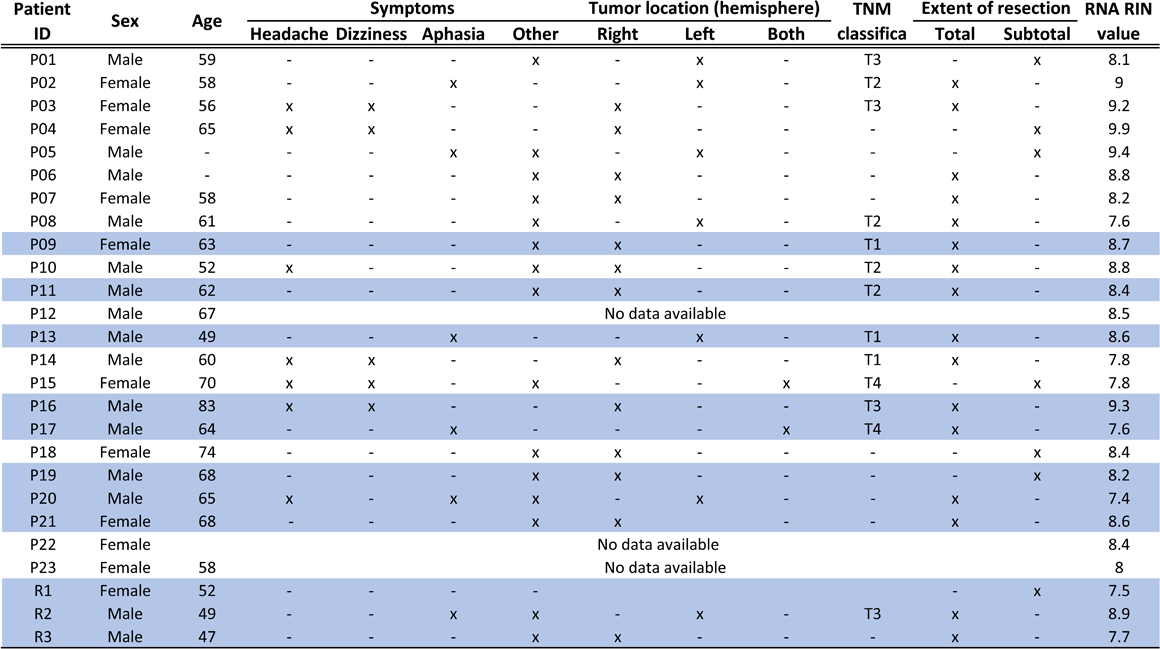
The characteristics of GBM tissue donors, who donated their tissue to the study. P – primary GBM, R – recurrent GBM; Classification of Malignant Tumors (TNM) according to <u>Union for International Cancer Control</u> (UICC) - T1, T2, T3, T4 represent size and/or extension of the primary tumor; RNA Integrity Number (RIN); samples subjected to experimental validation marked in blue.

**Supplementary Table 2.**
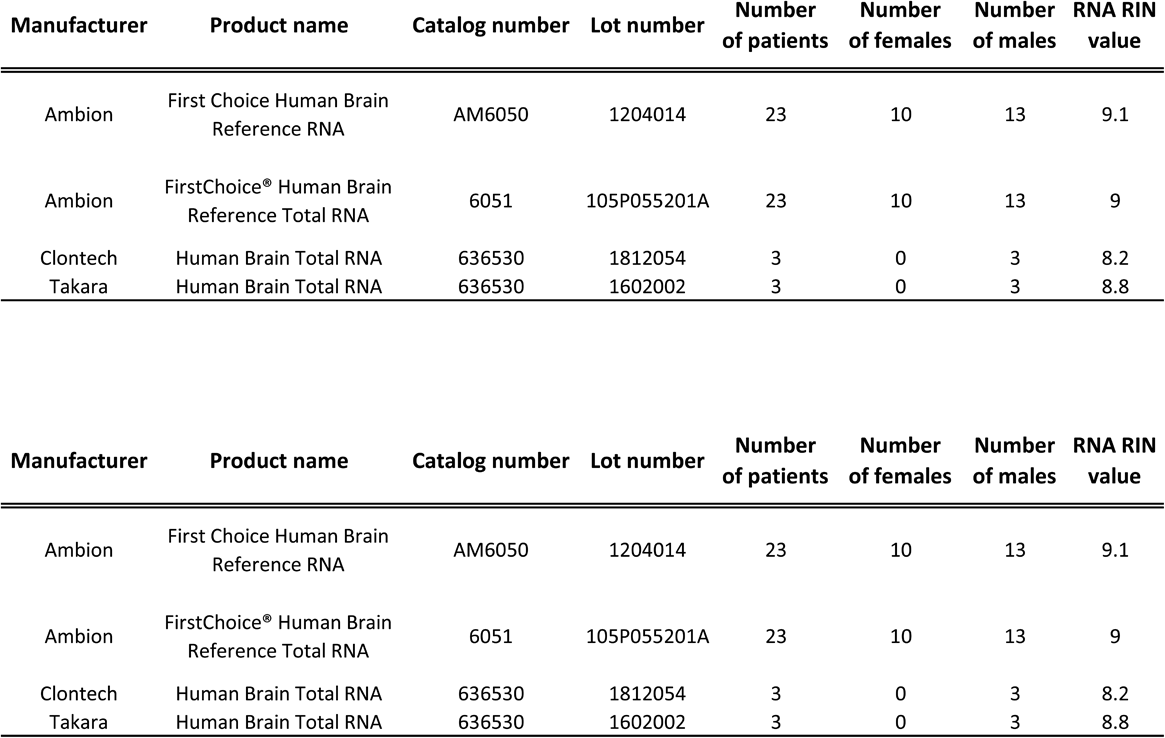
The characteristics of HB controls used in the study. RNA Integrity Number (RIN).

**Supplementary Table 3.**
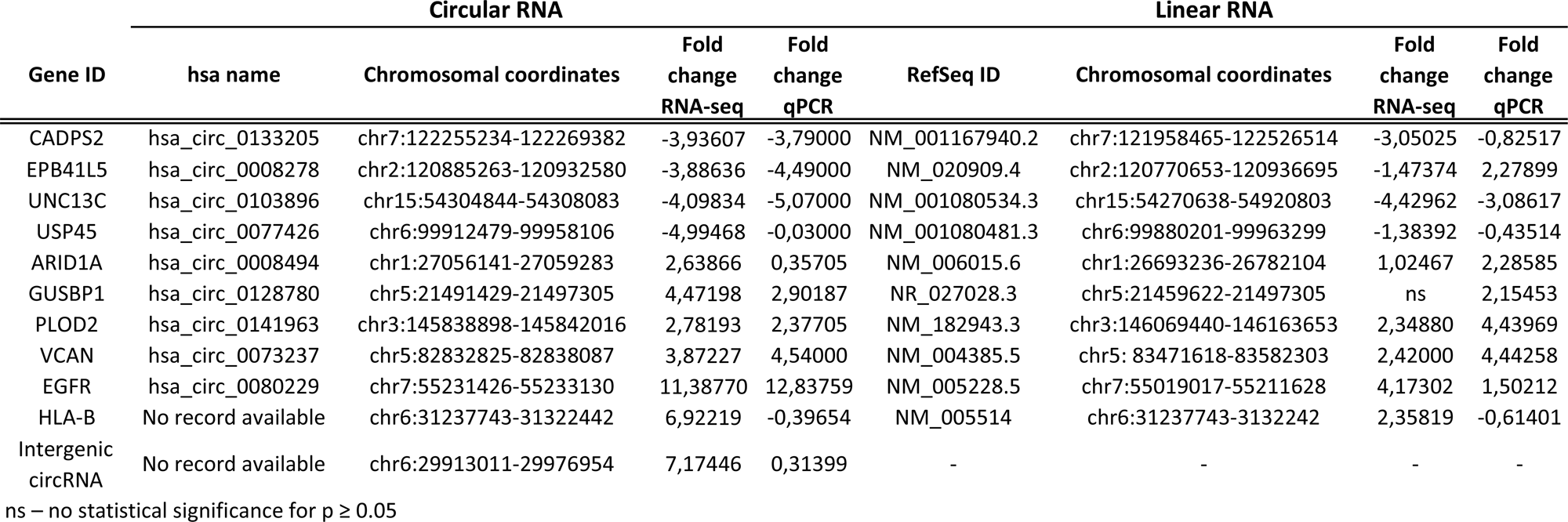
Characteristics of the circular and linear transcripts subjected to the validation. ns – no statistical significance for p ≥ 0.05.

**Supplementary Table 4.**
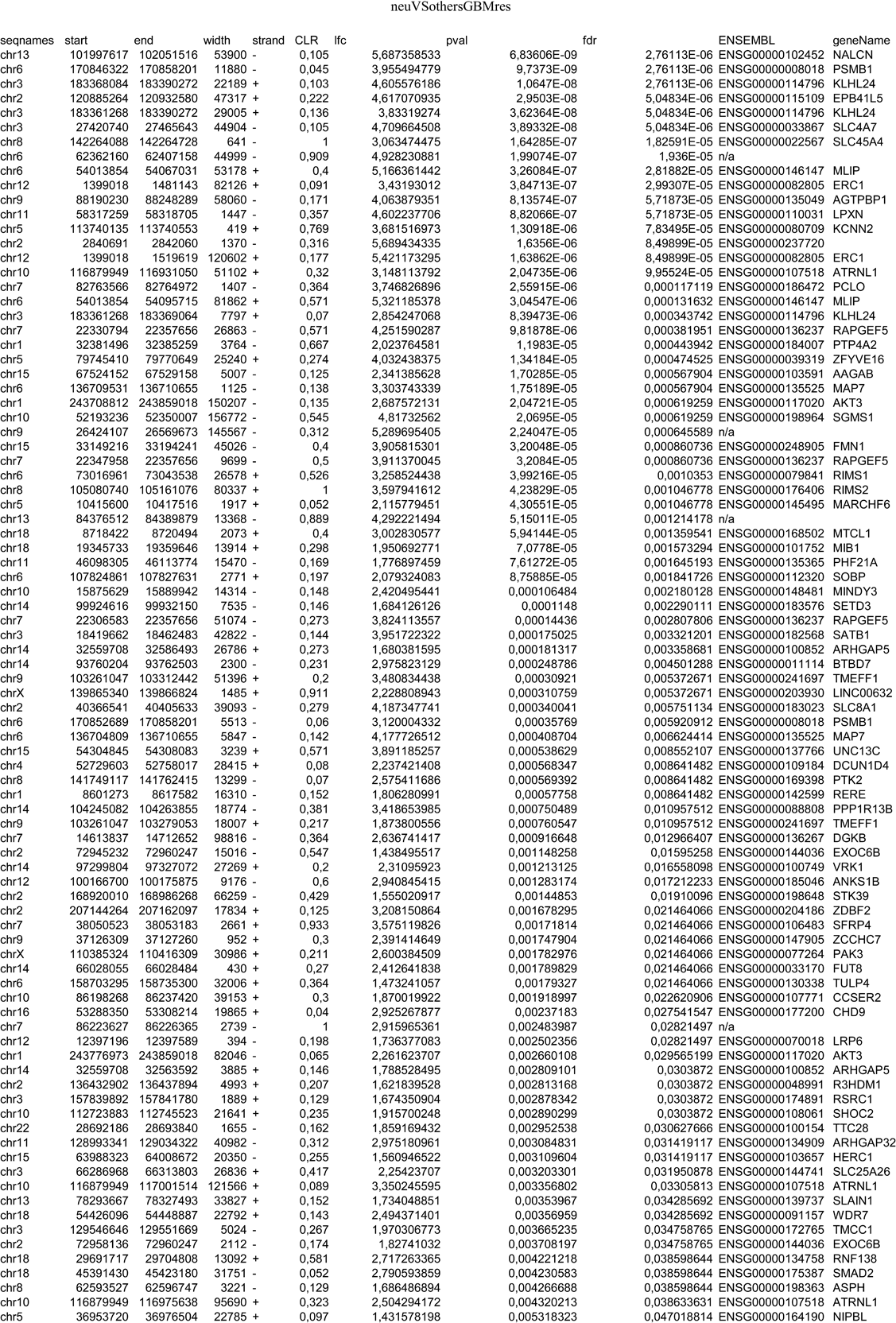
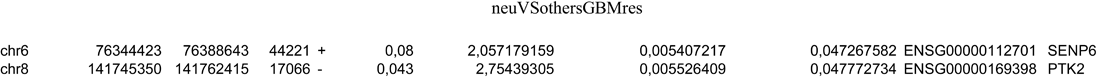
CircRNAs differentially expressed in neural subtype versus other subtypes.

**Supplementary Table 5.**
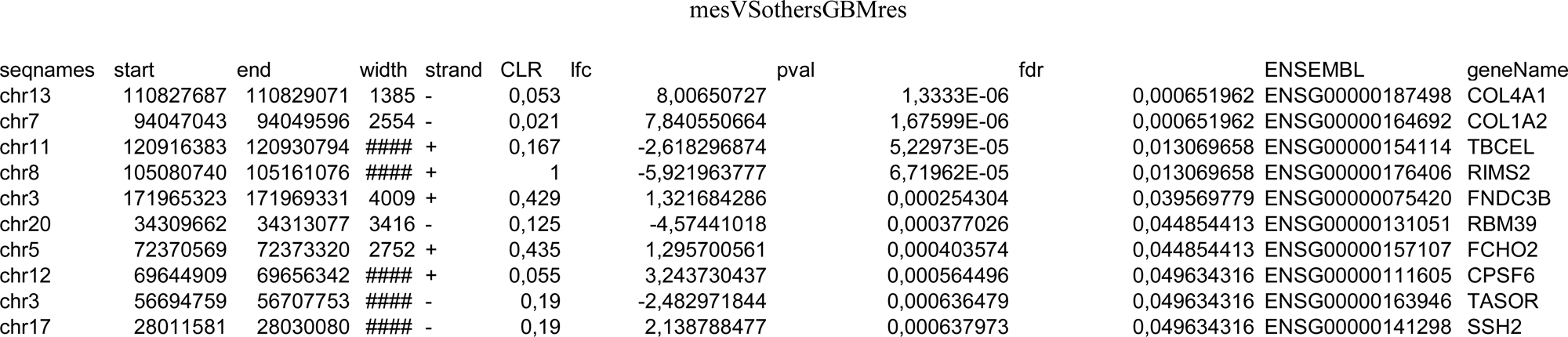
CircRNAs differentially expressed in mesenchymal subtype versus other subtypes.

**Supplementary Table 6.**
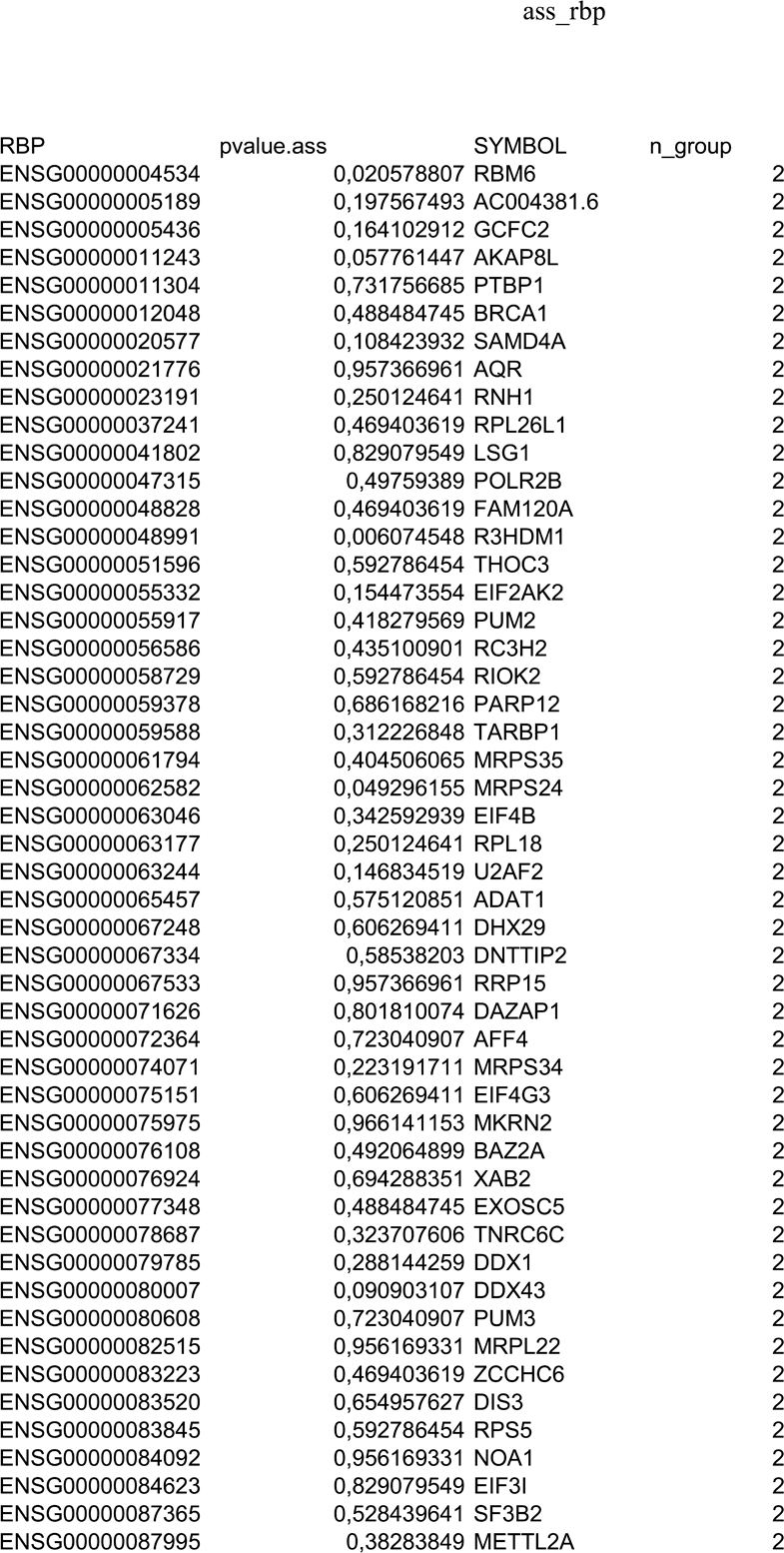
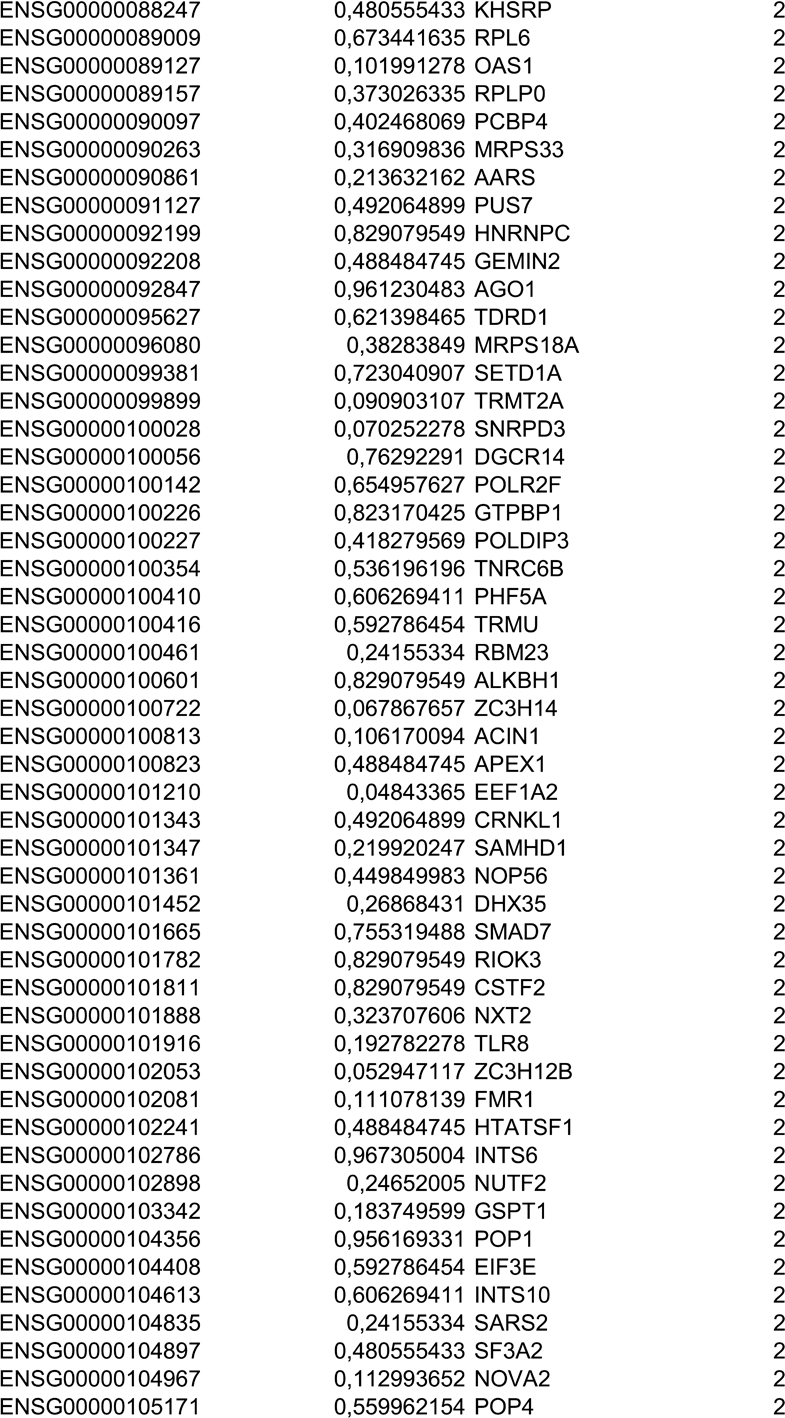
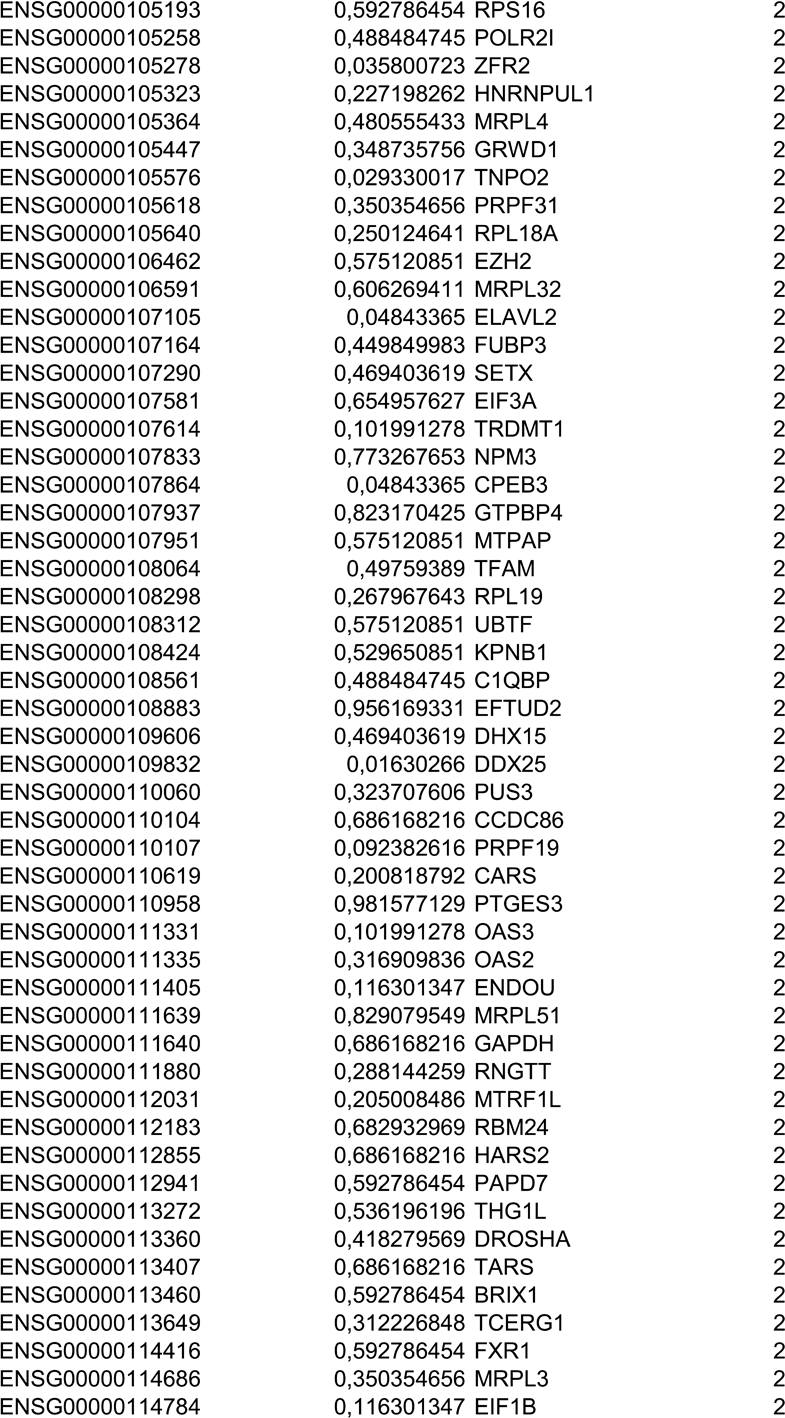
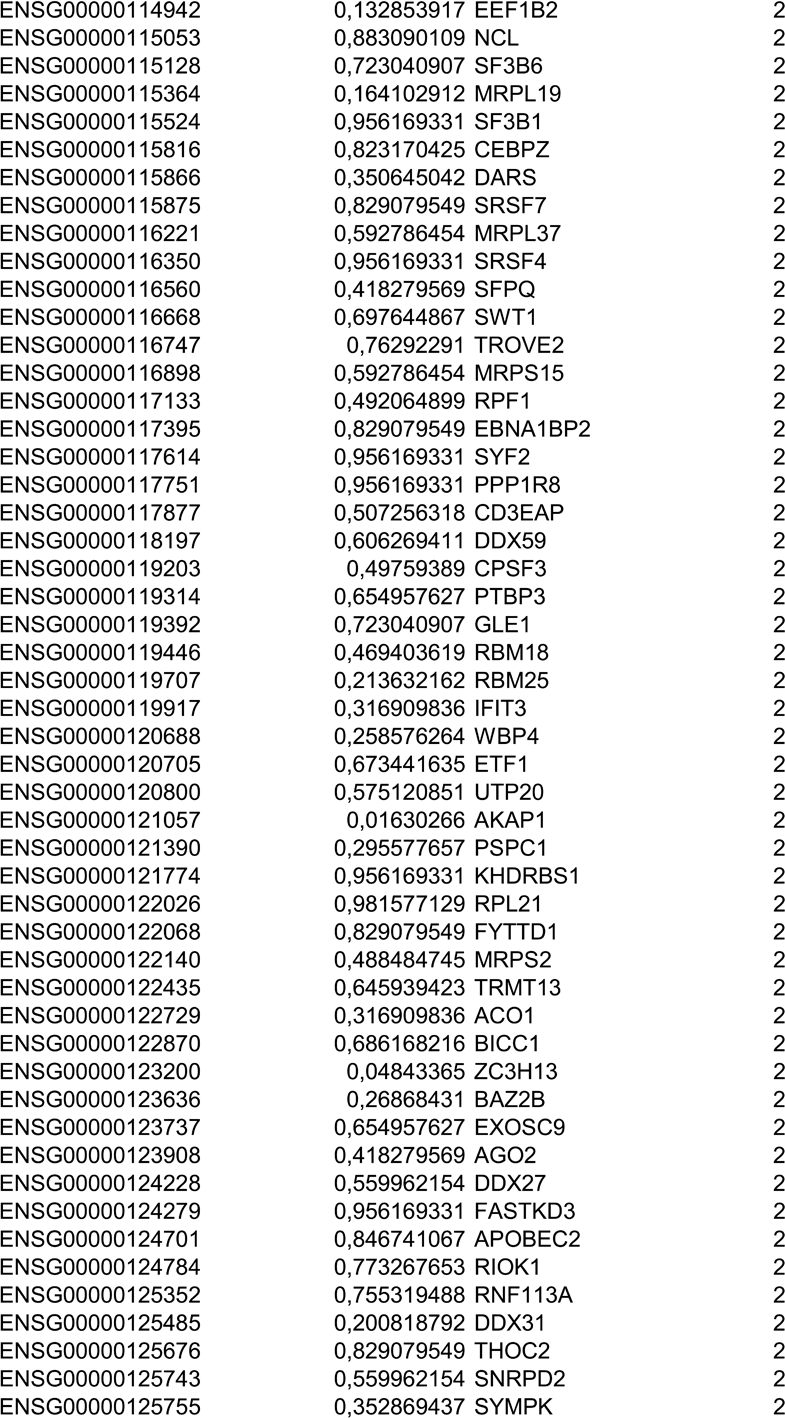
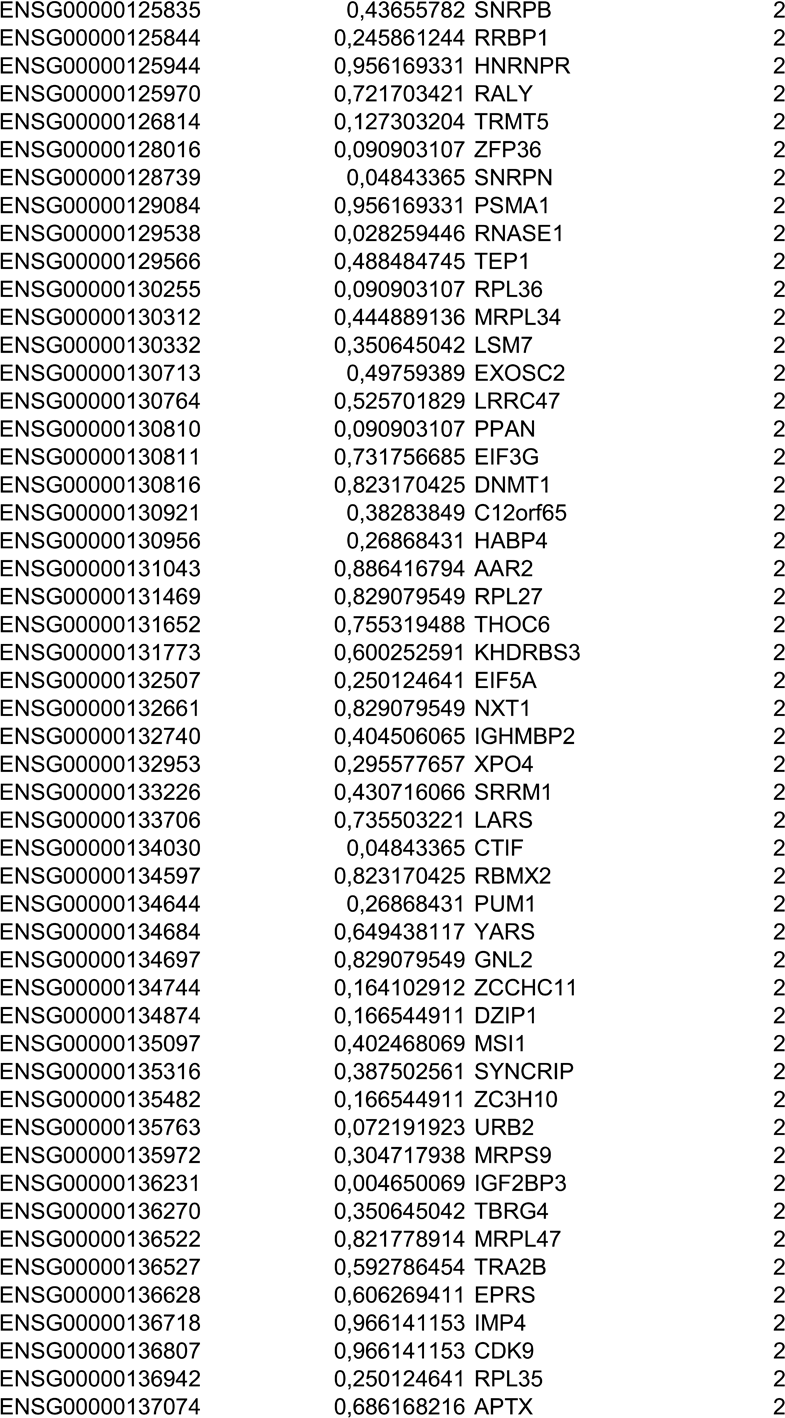
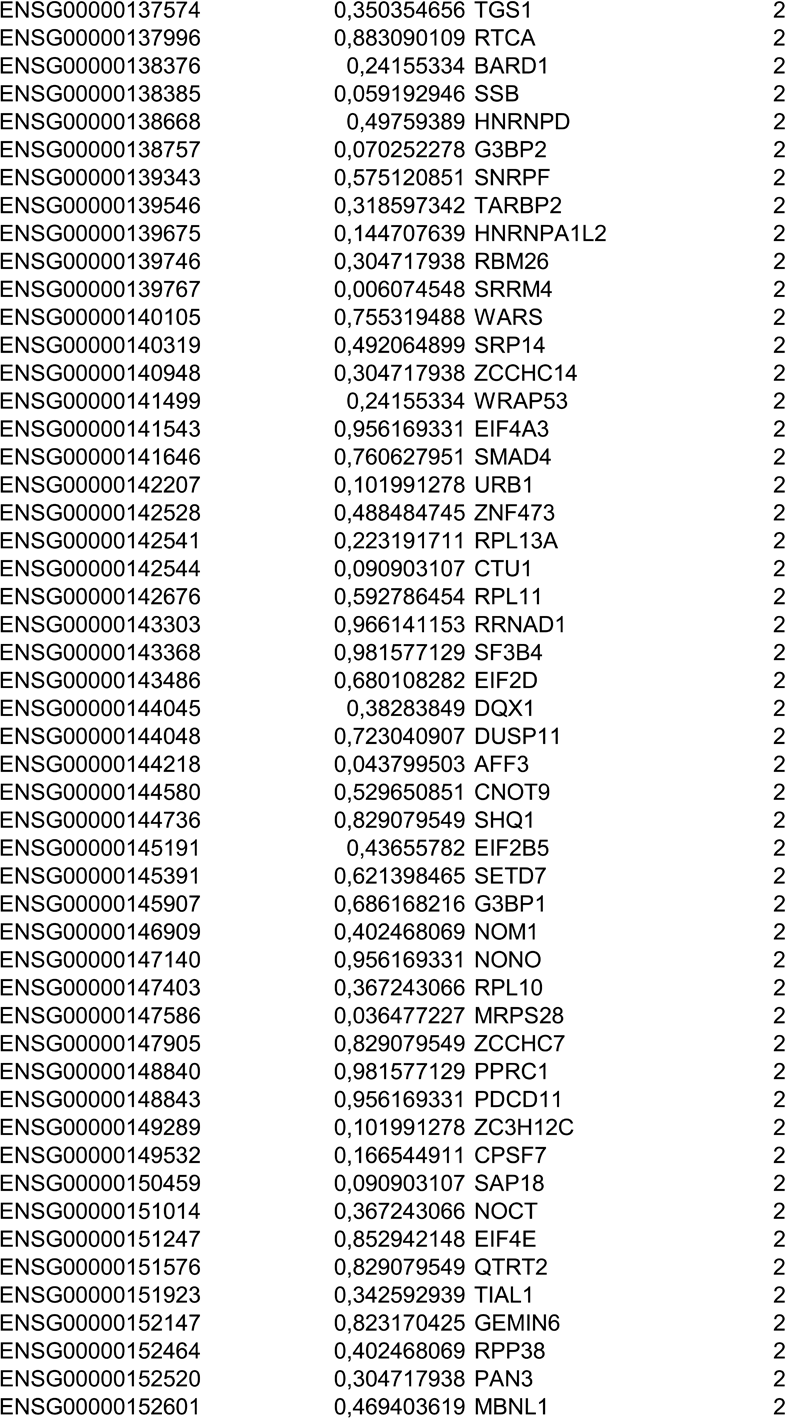
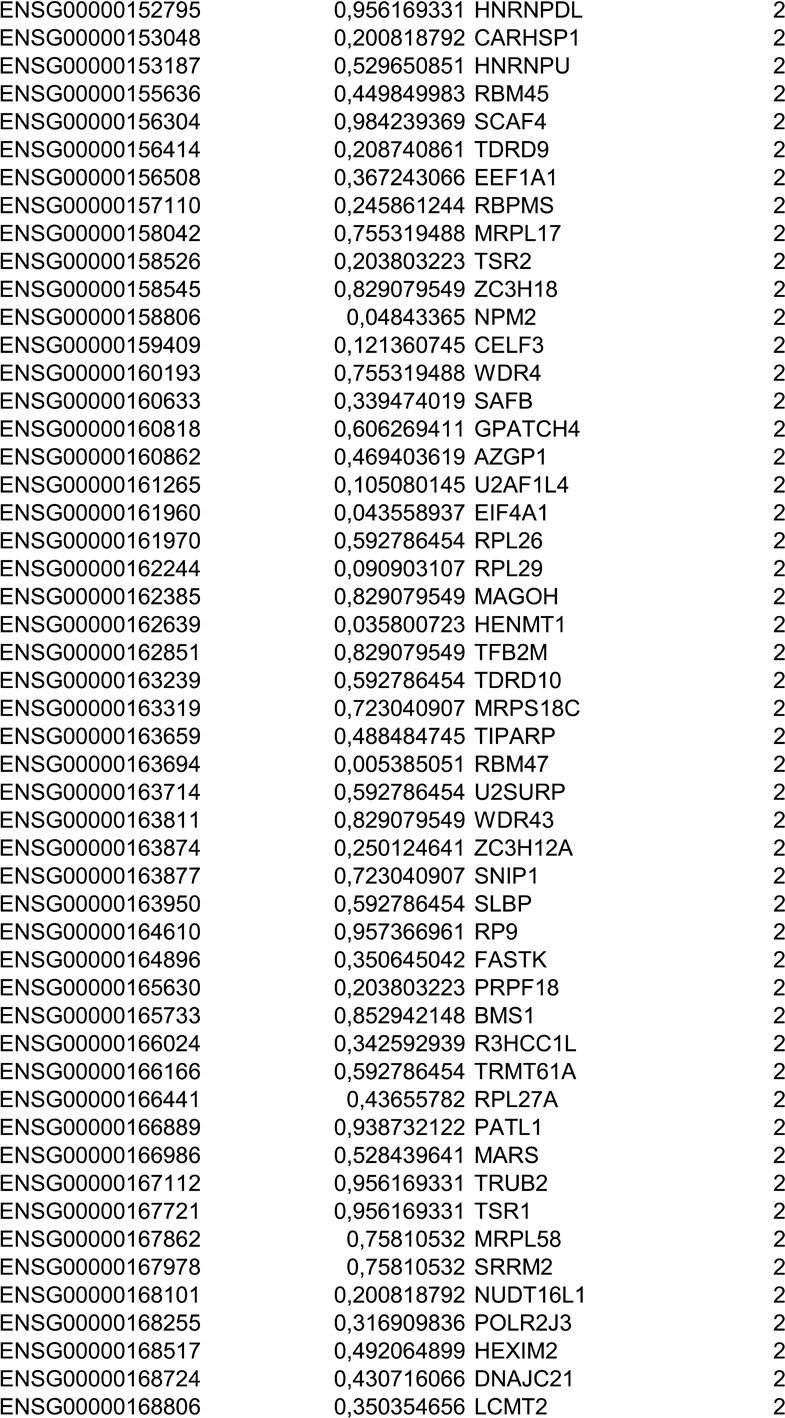
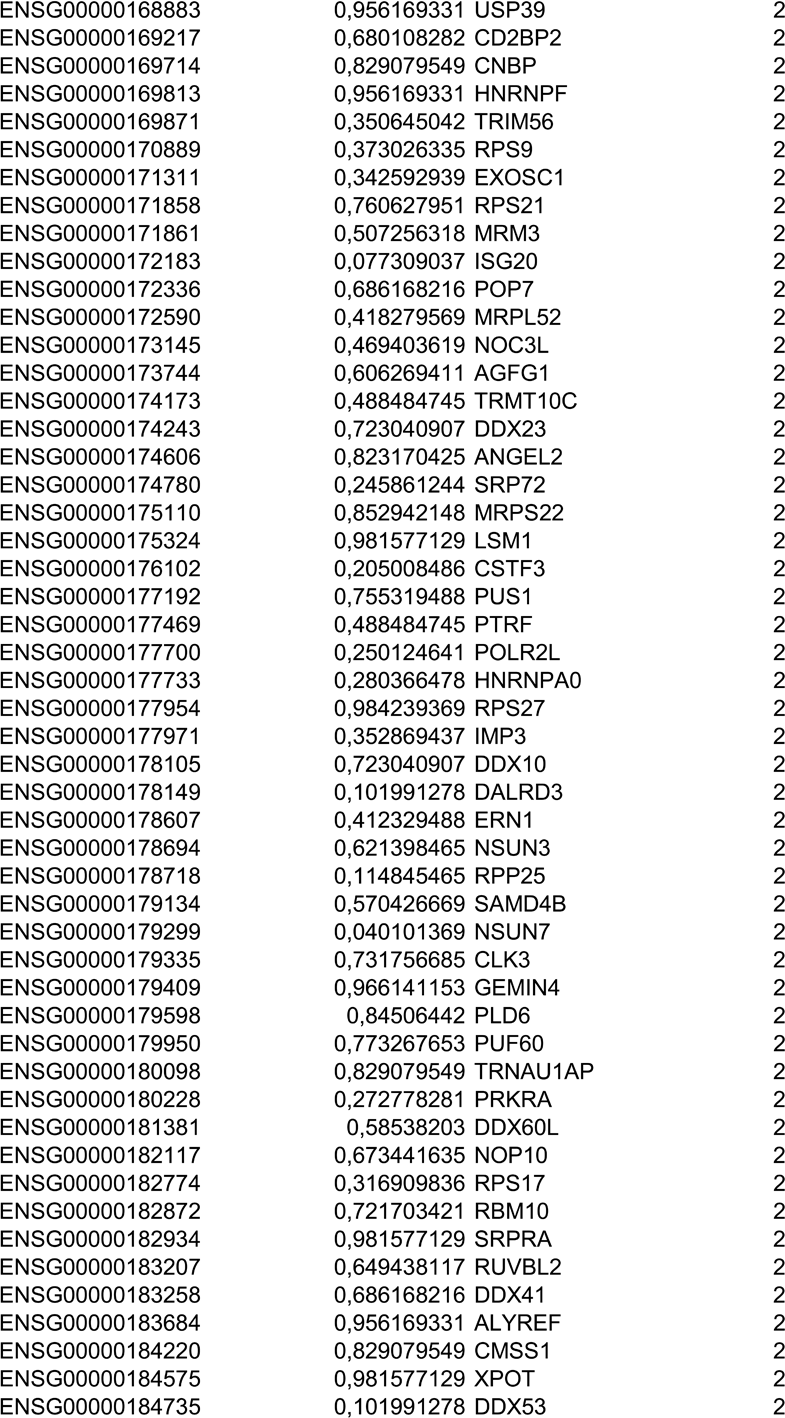
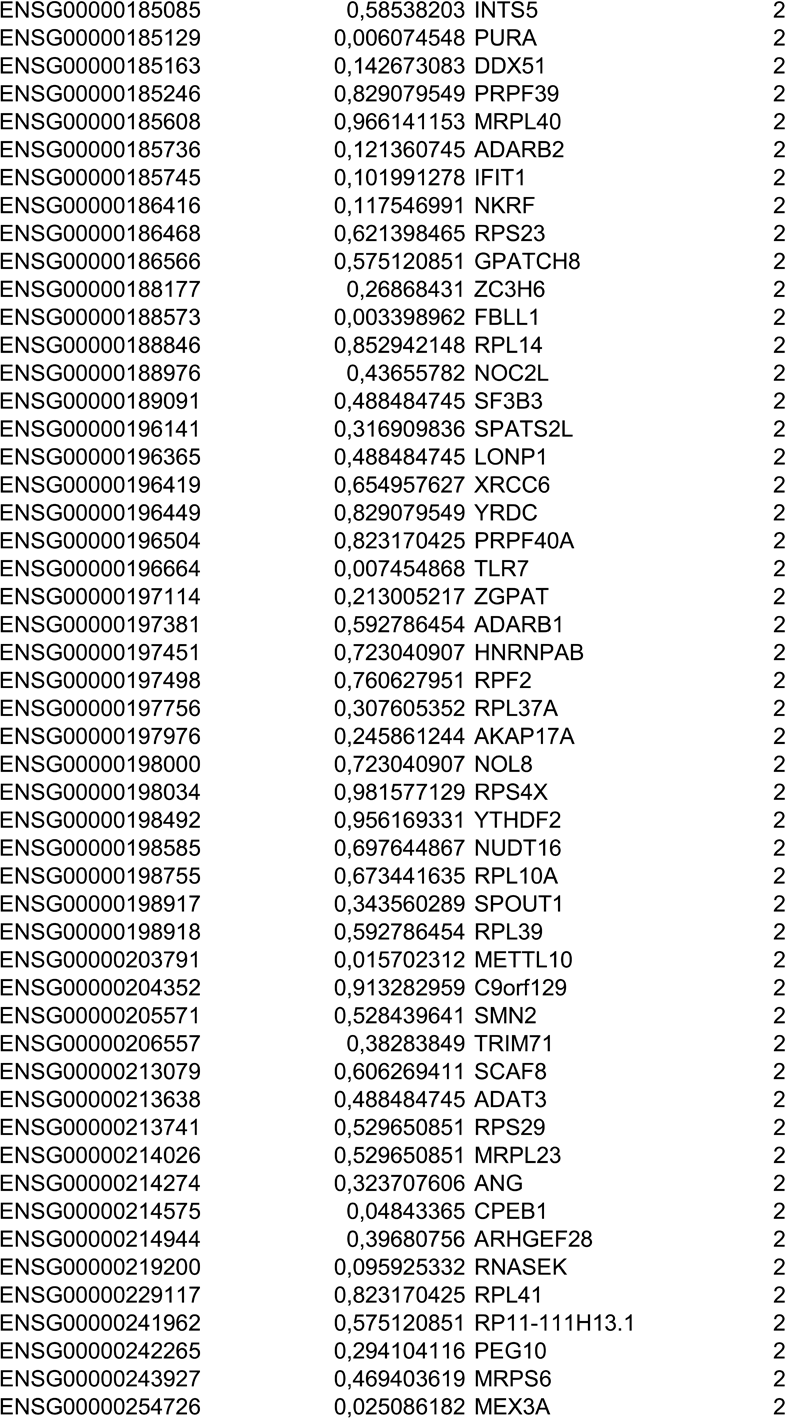
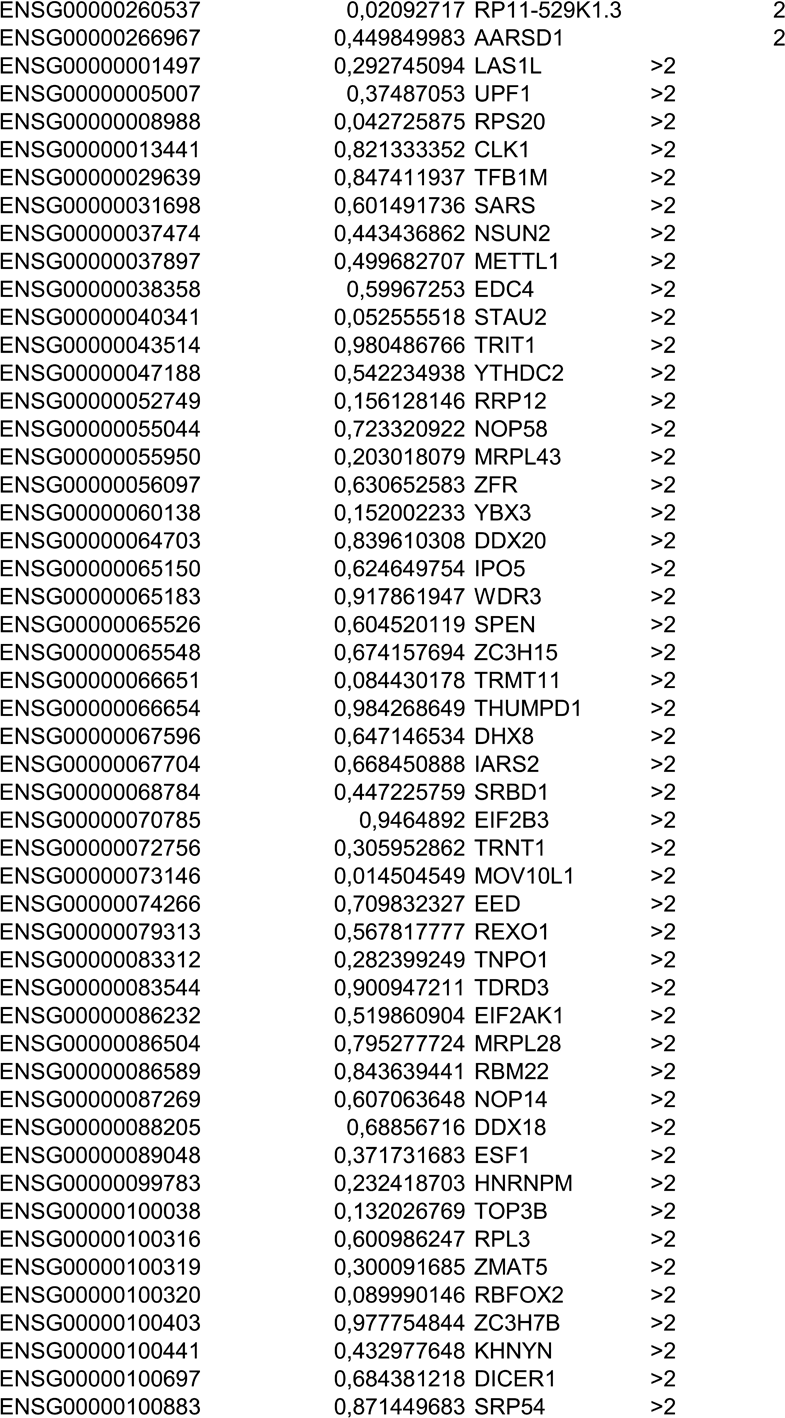
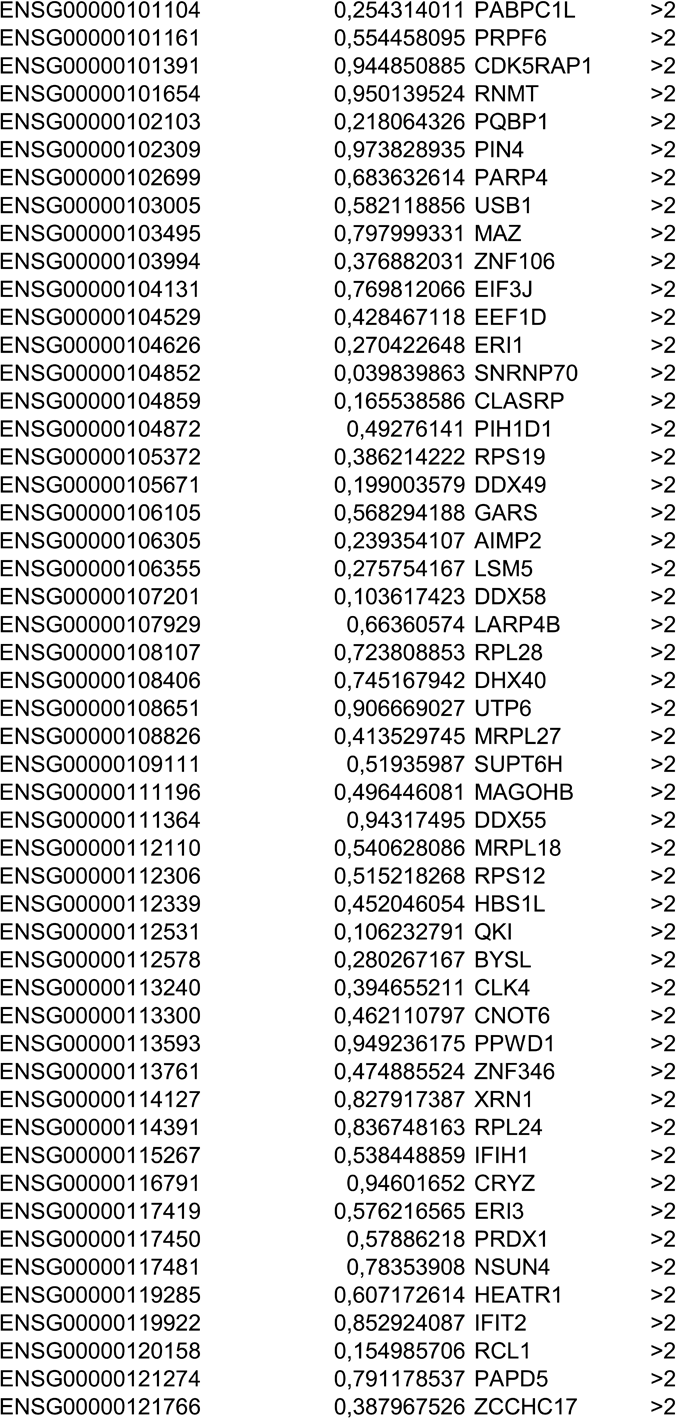
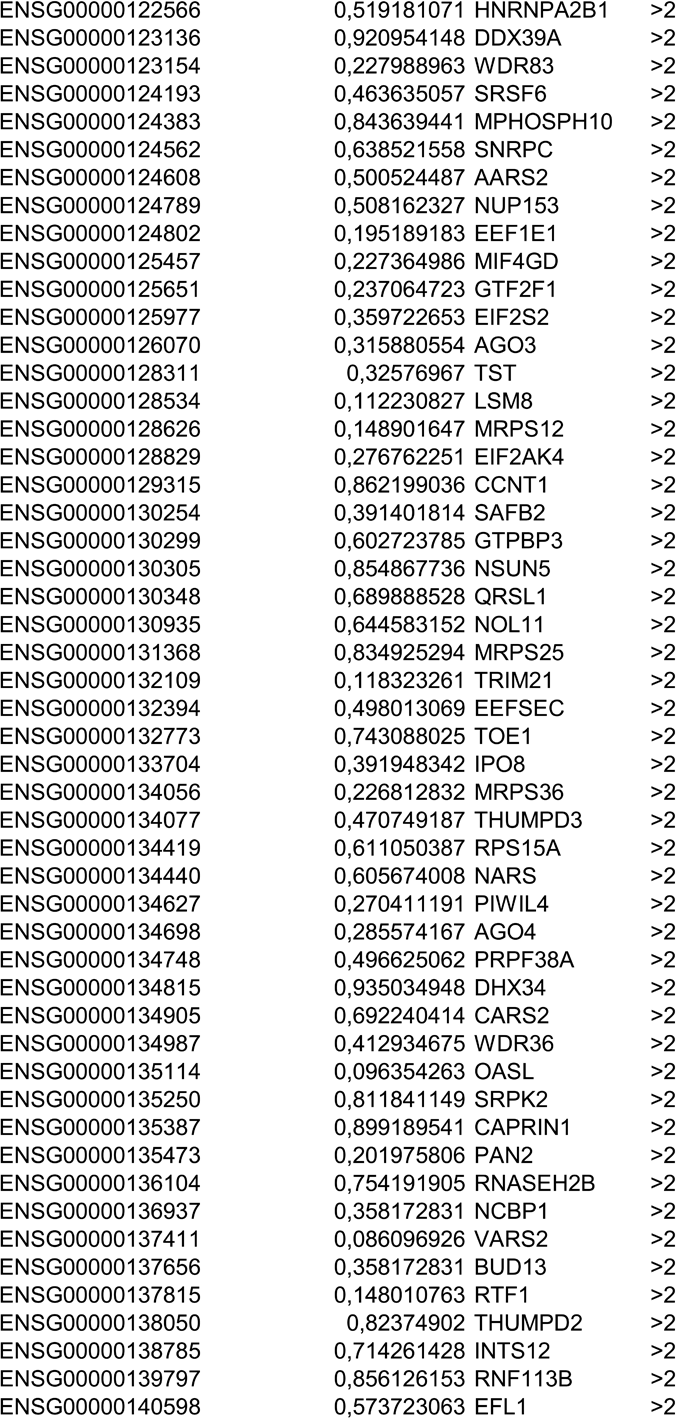
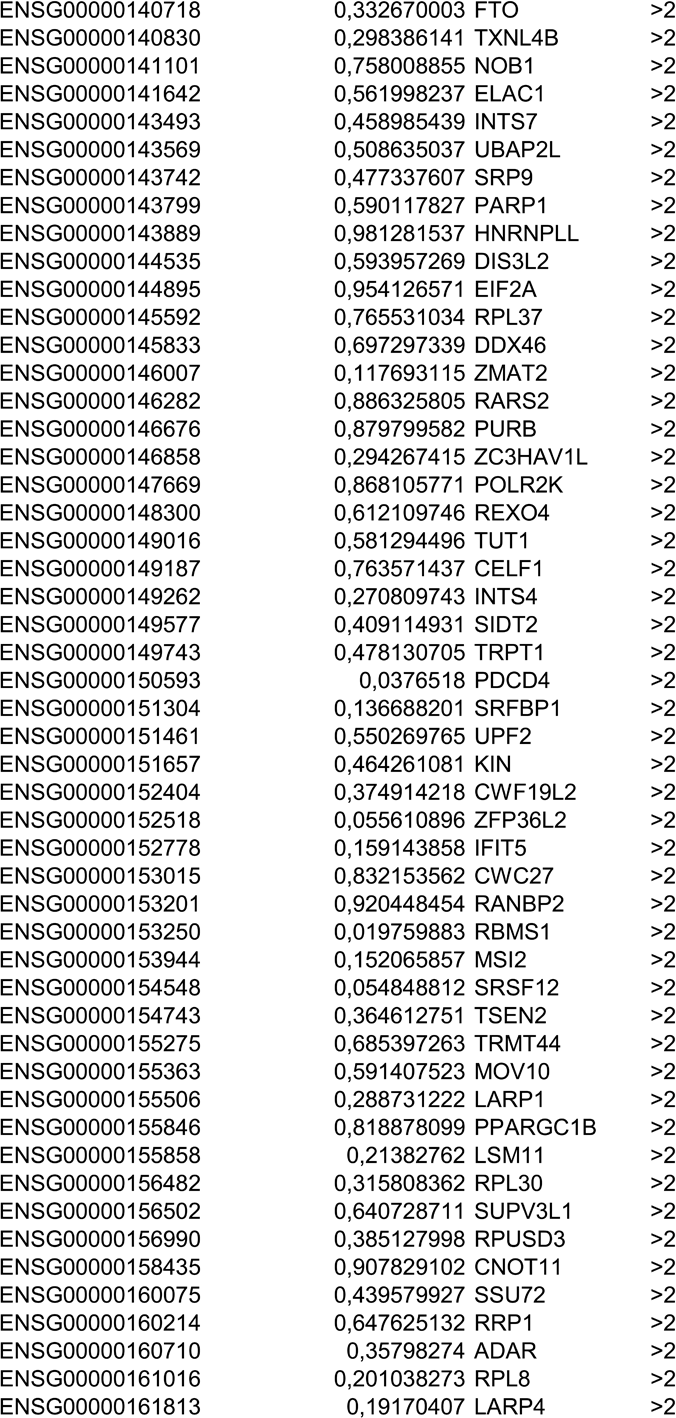
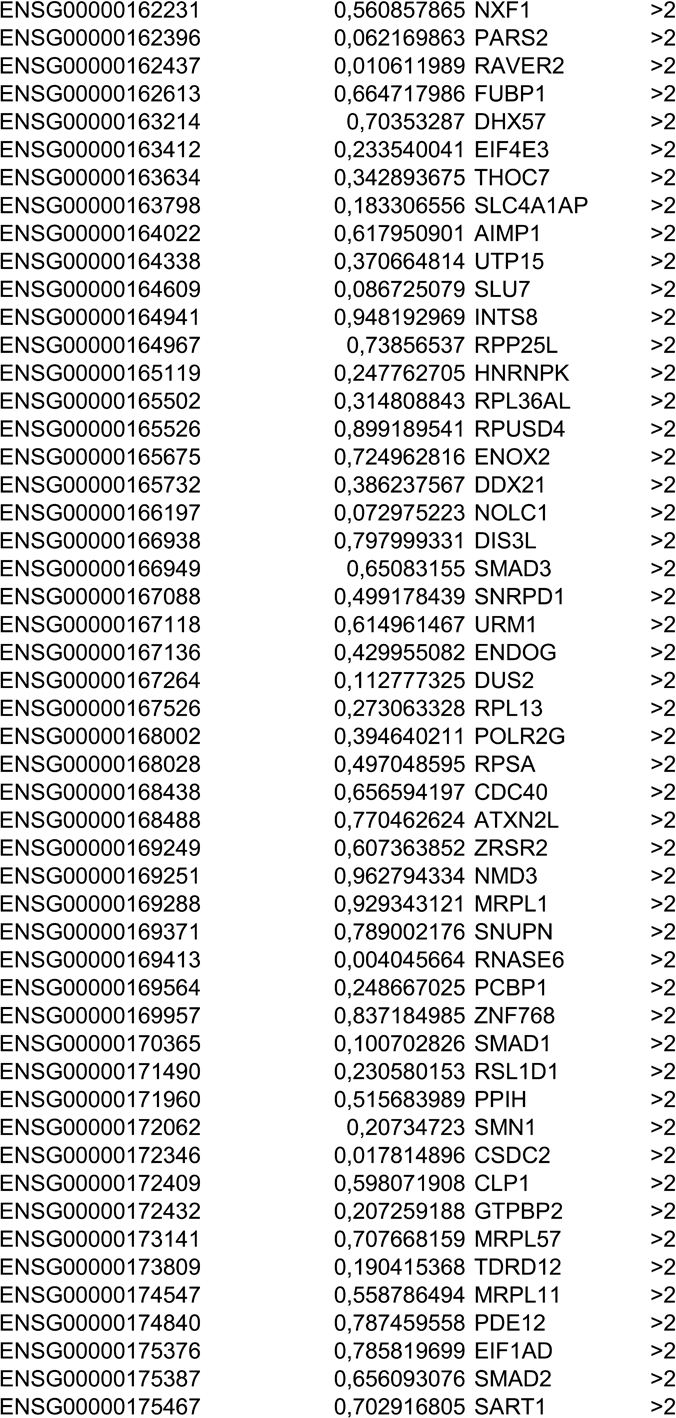
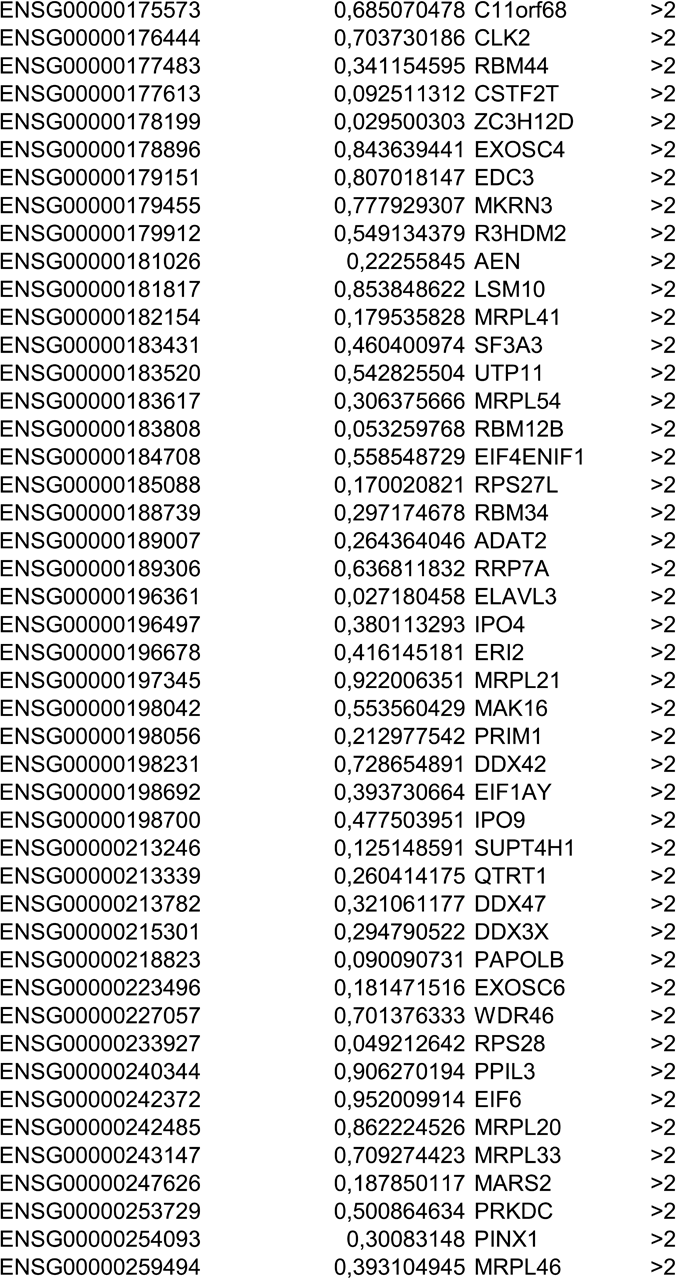
Differentially expressed RBPs used for stratification of GBM samples into different groups.

**Supplementary Table 7.**
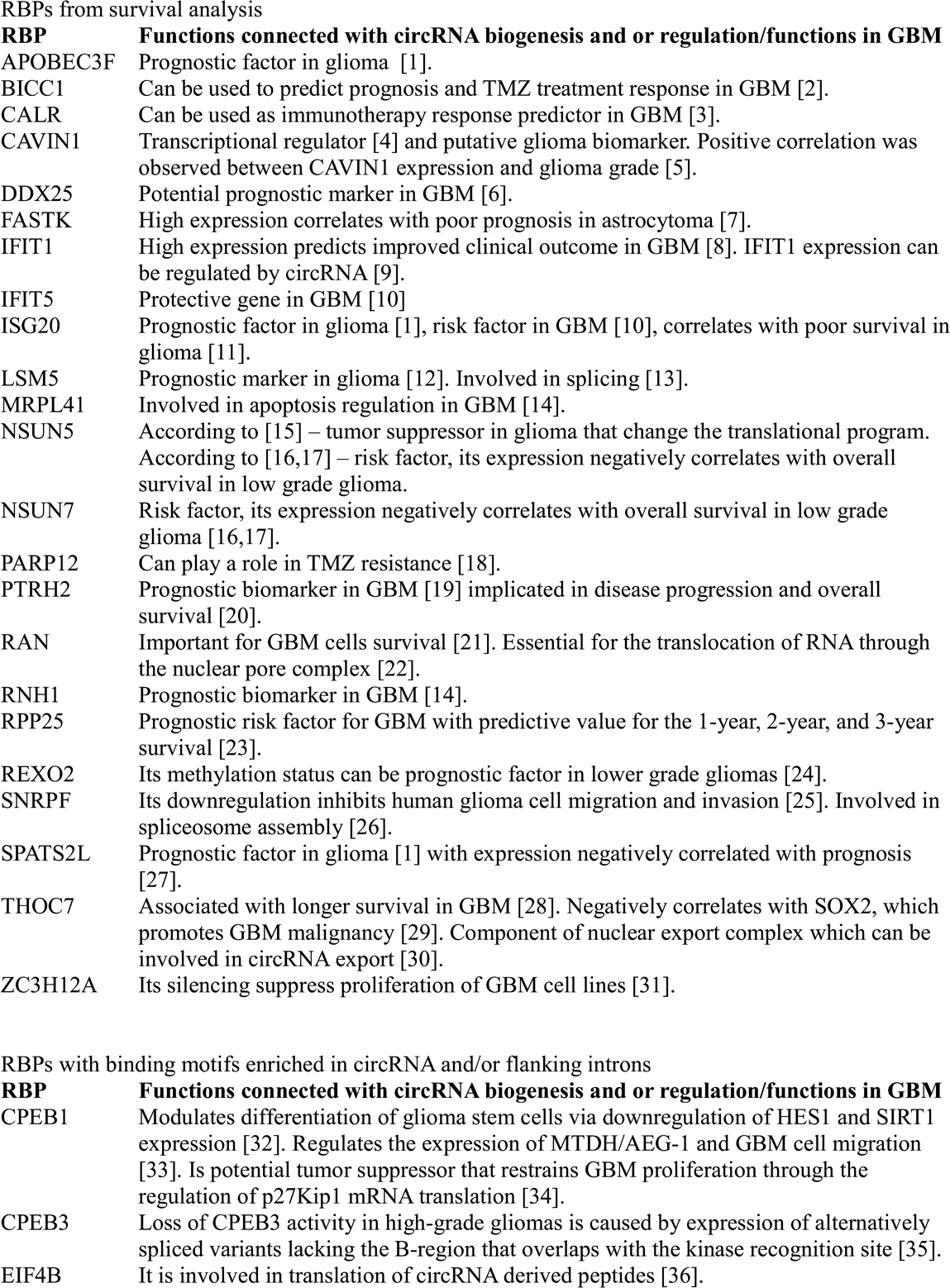
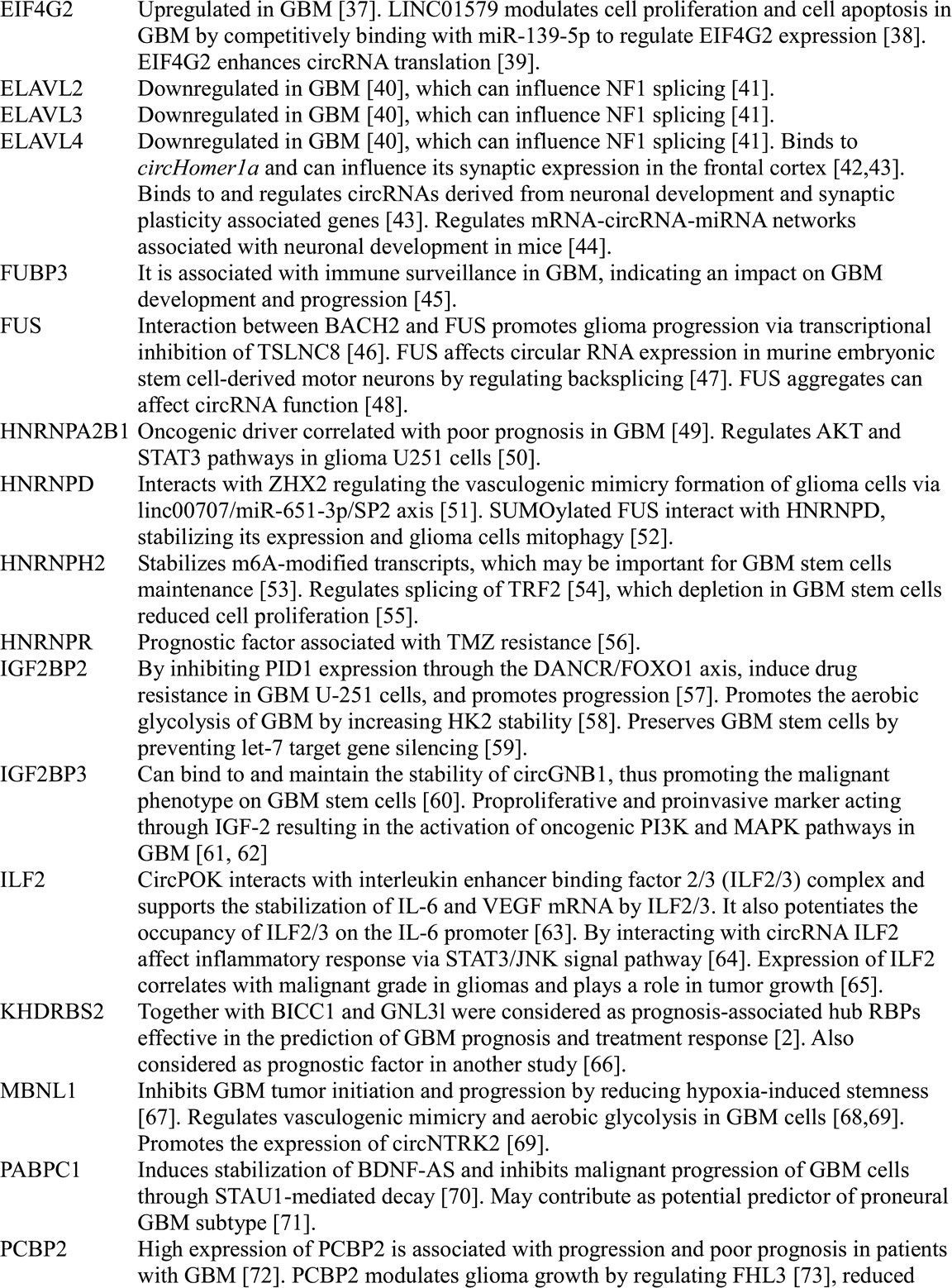
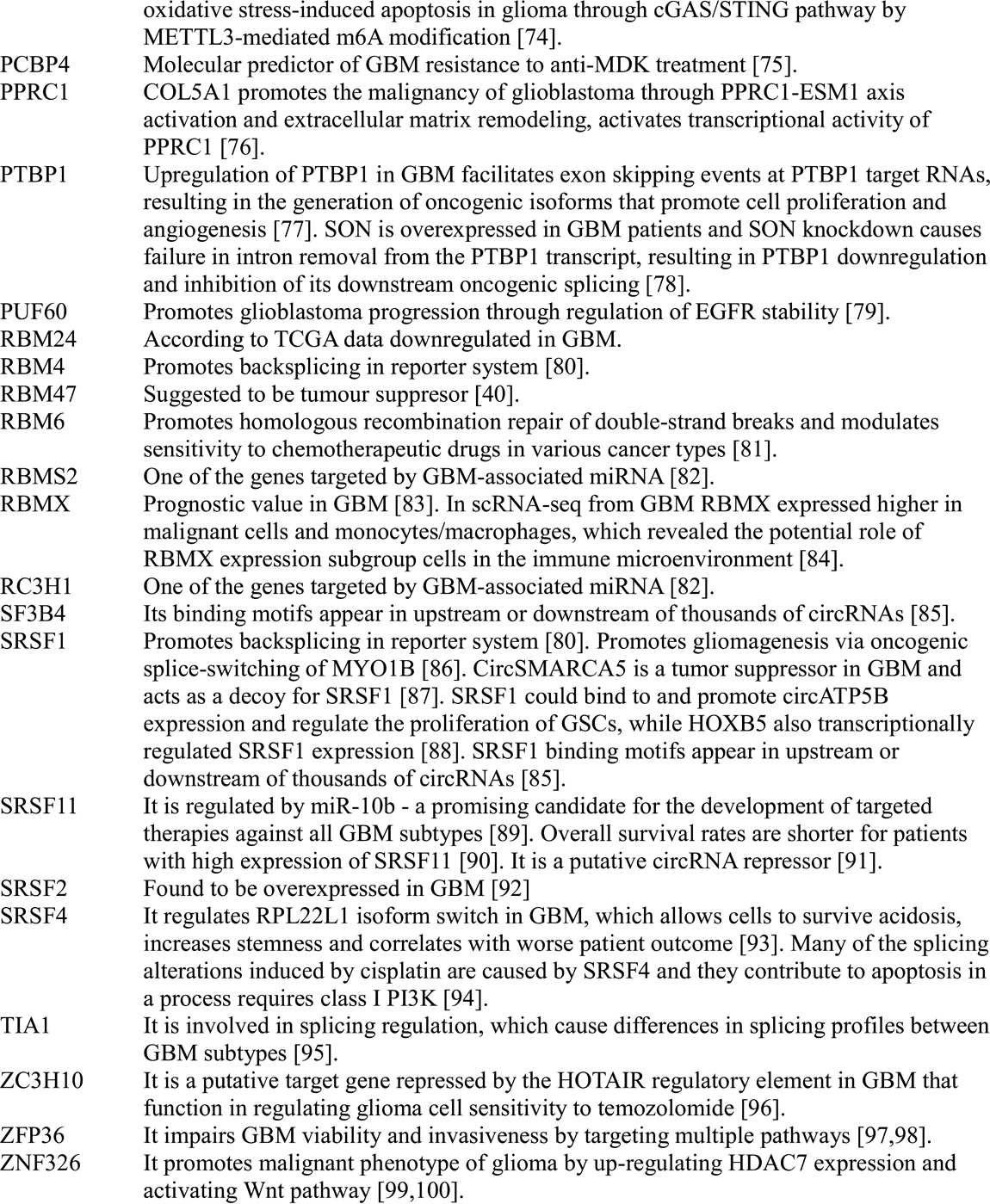
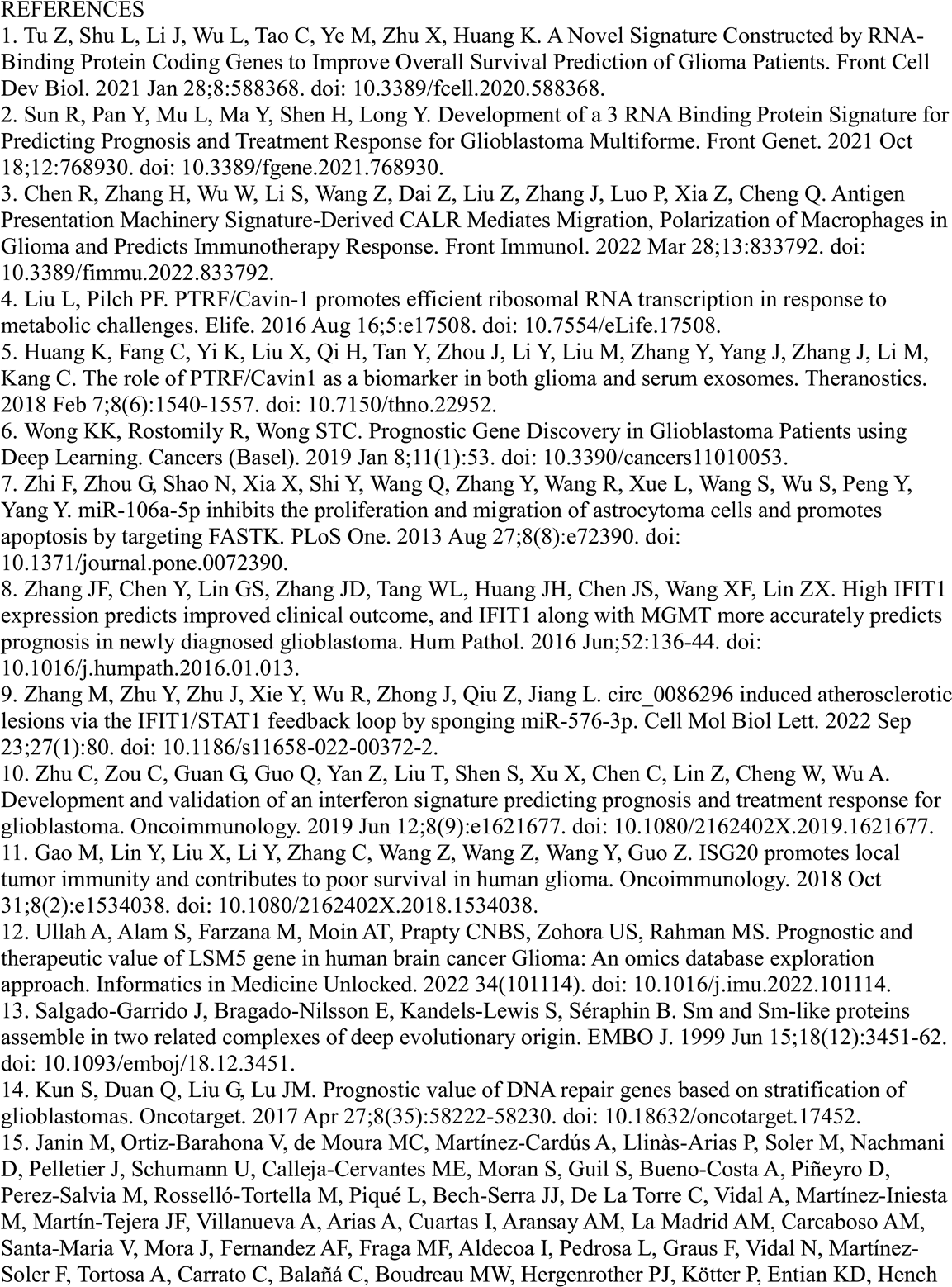
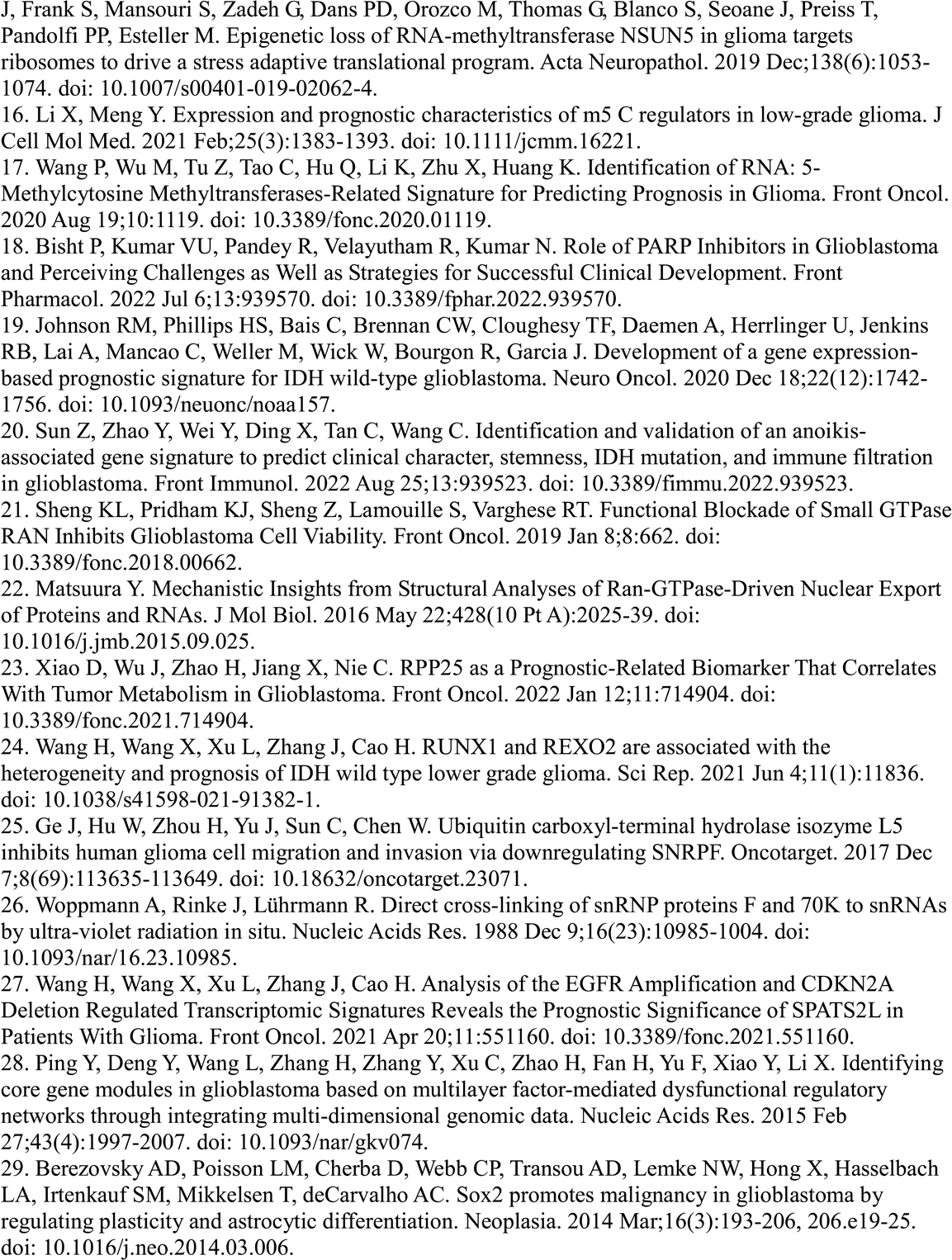
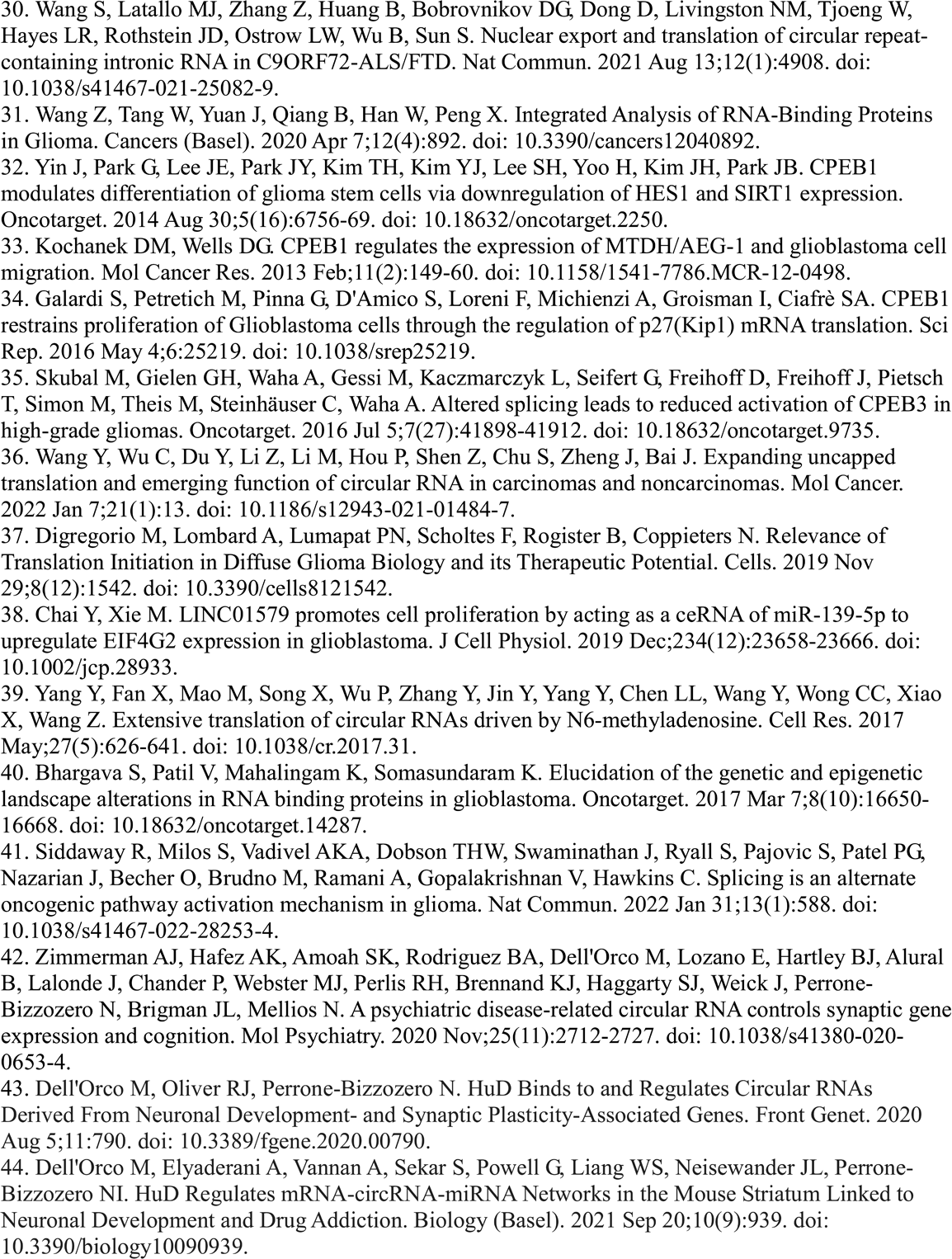
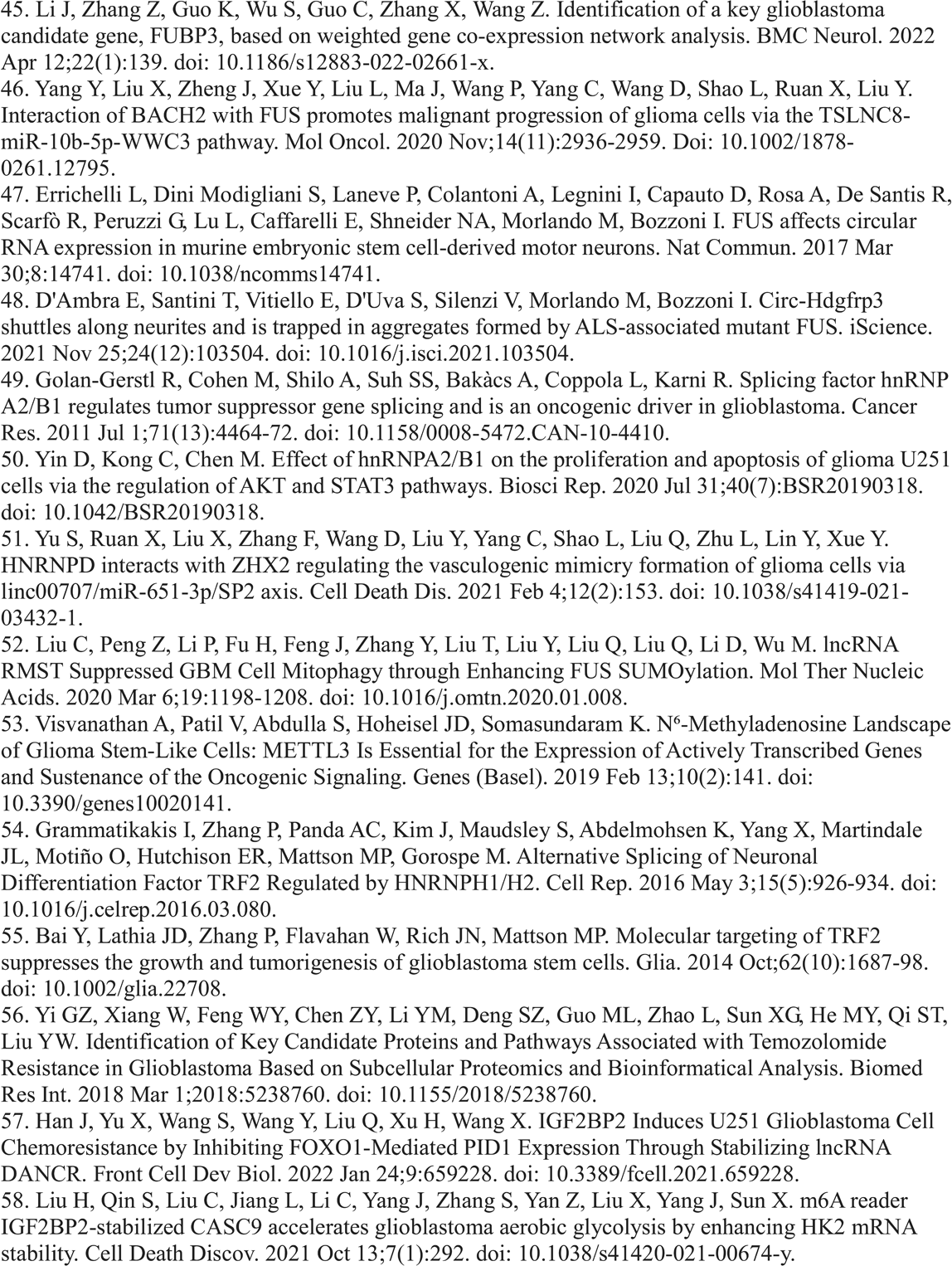
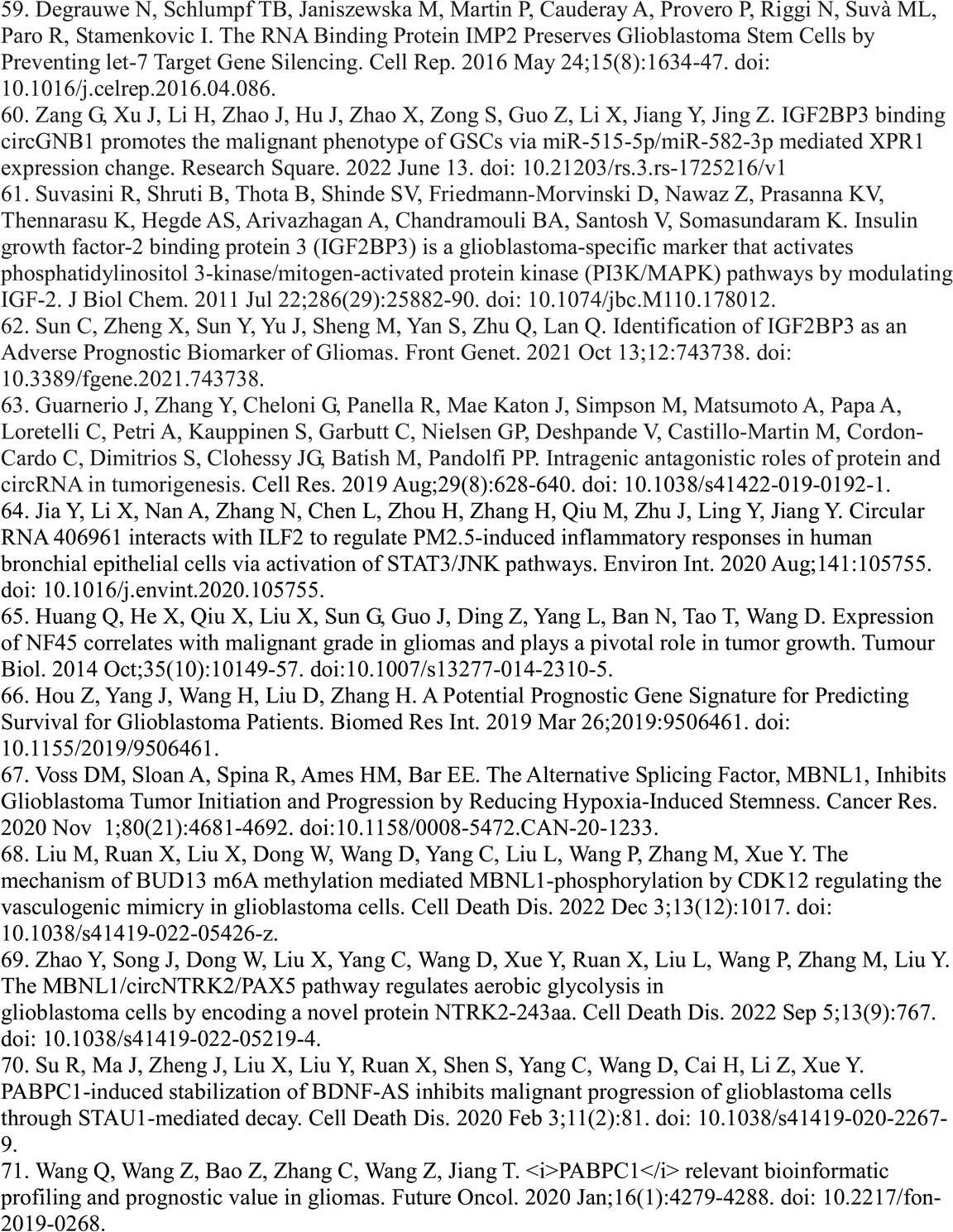
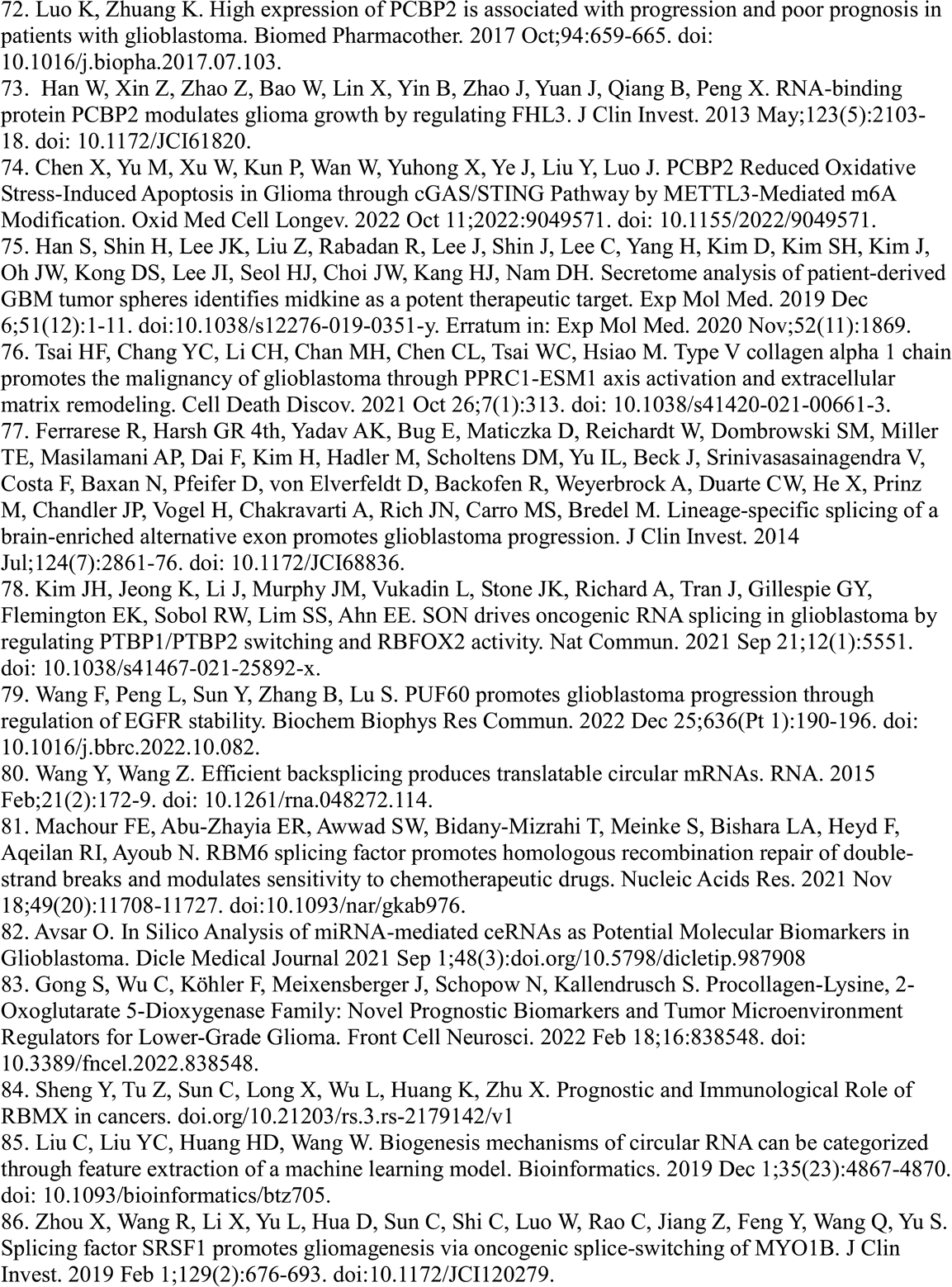
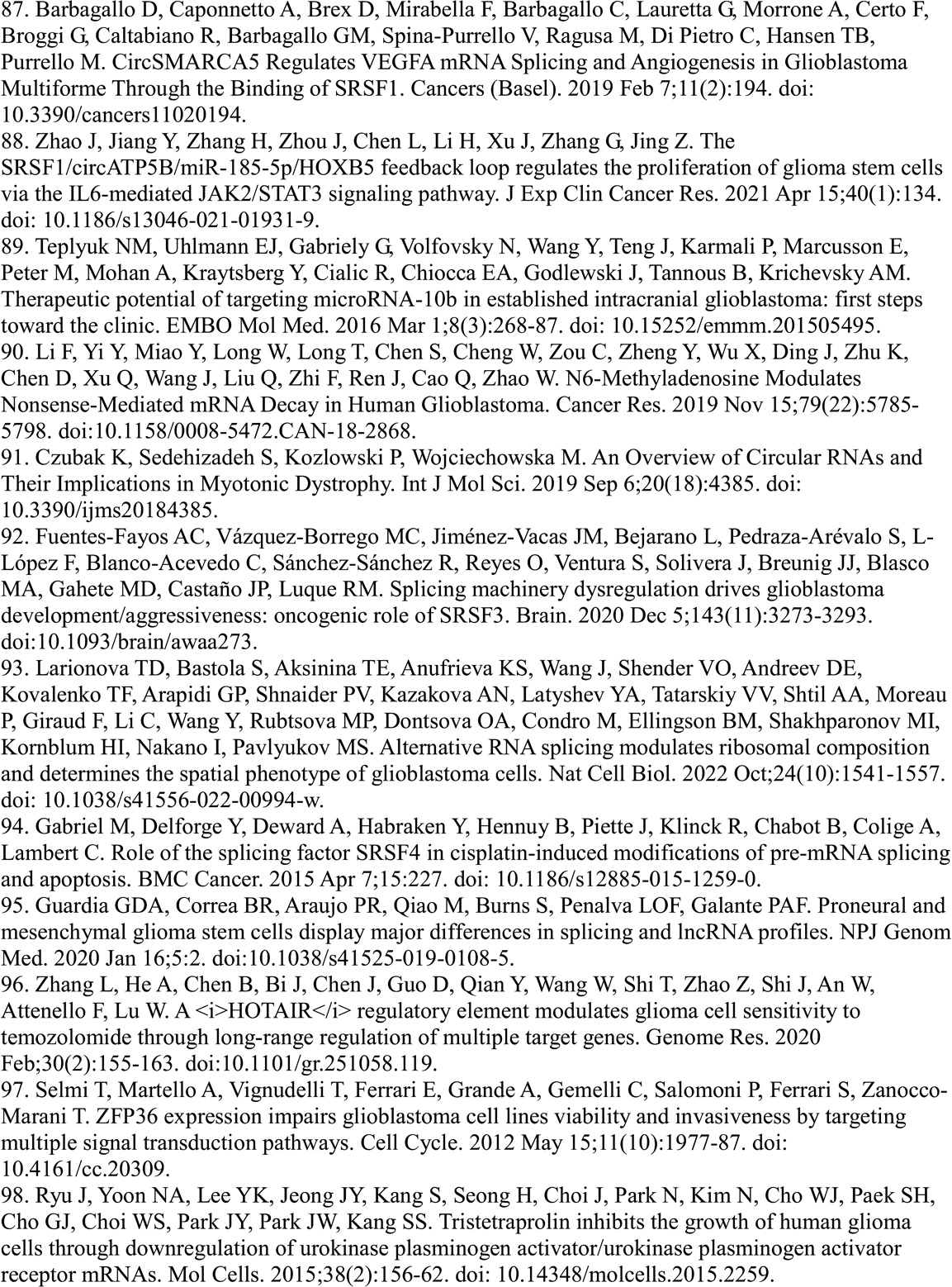
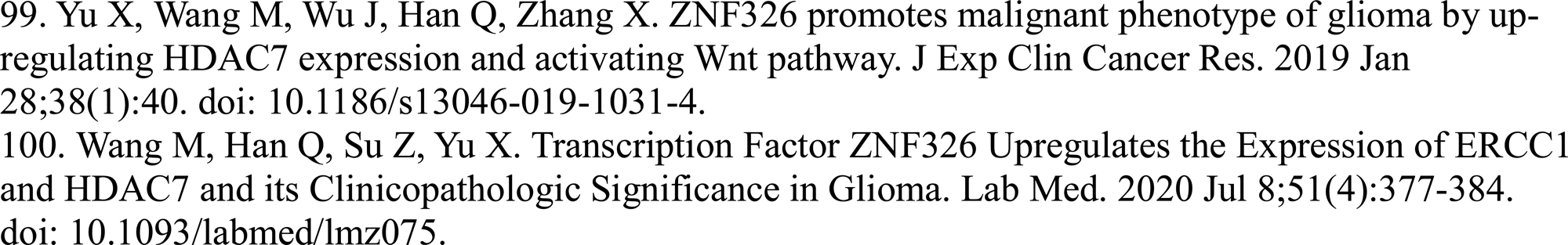
RBPs shown to be brain tumor markers, correlated with disease outcome or response to treatment.

**Supplementary Table 8.**
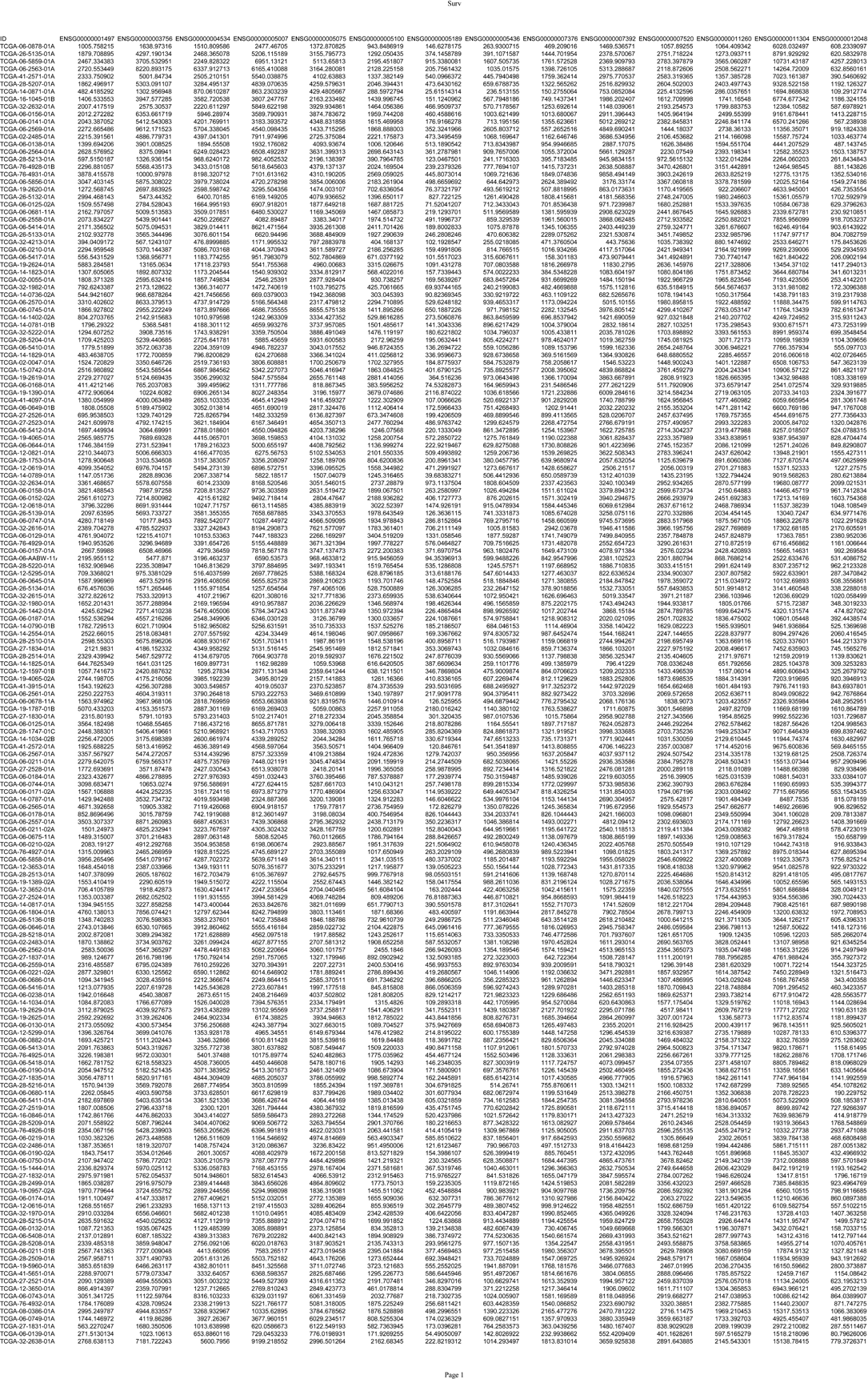
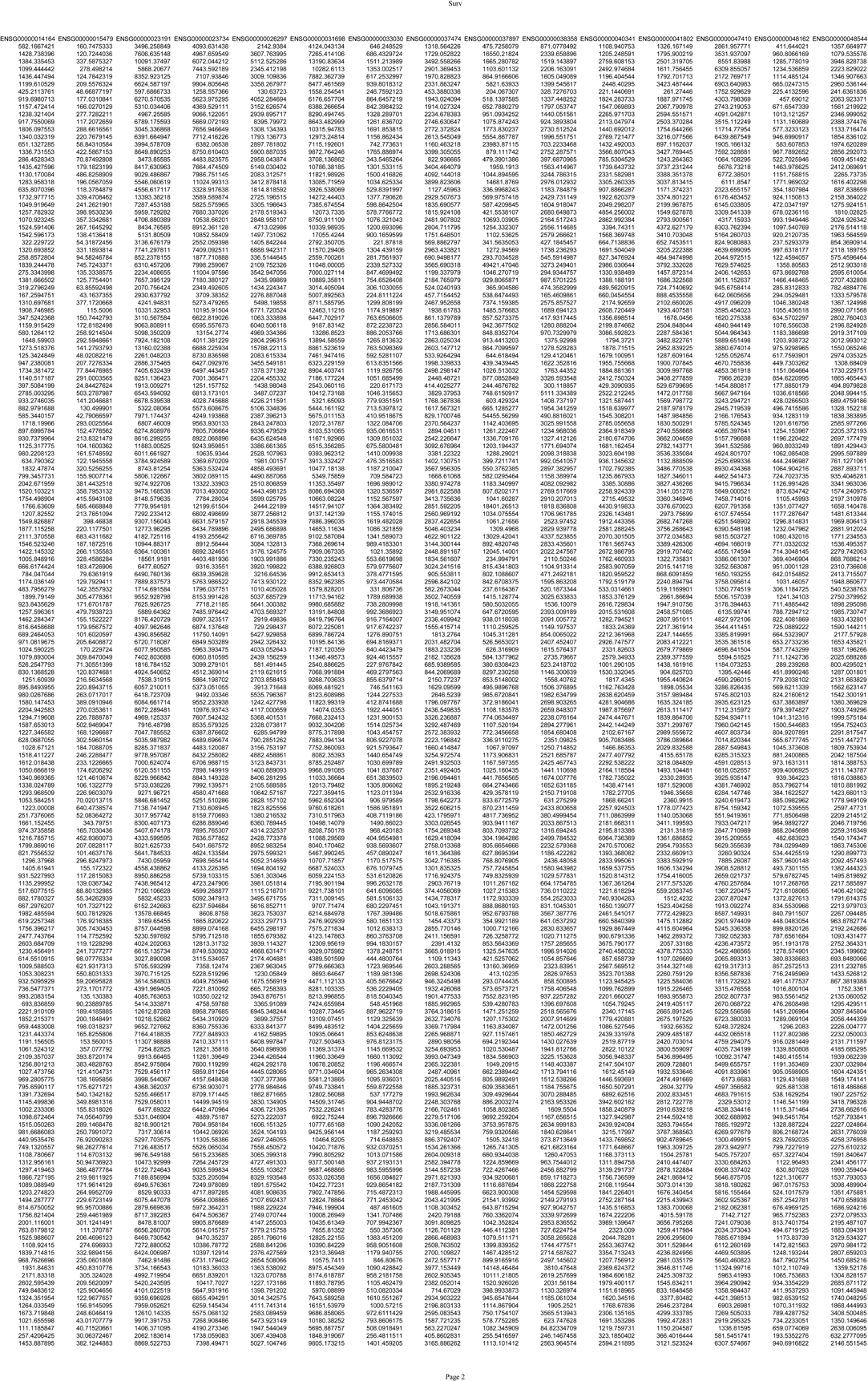
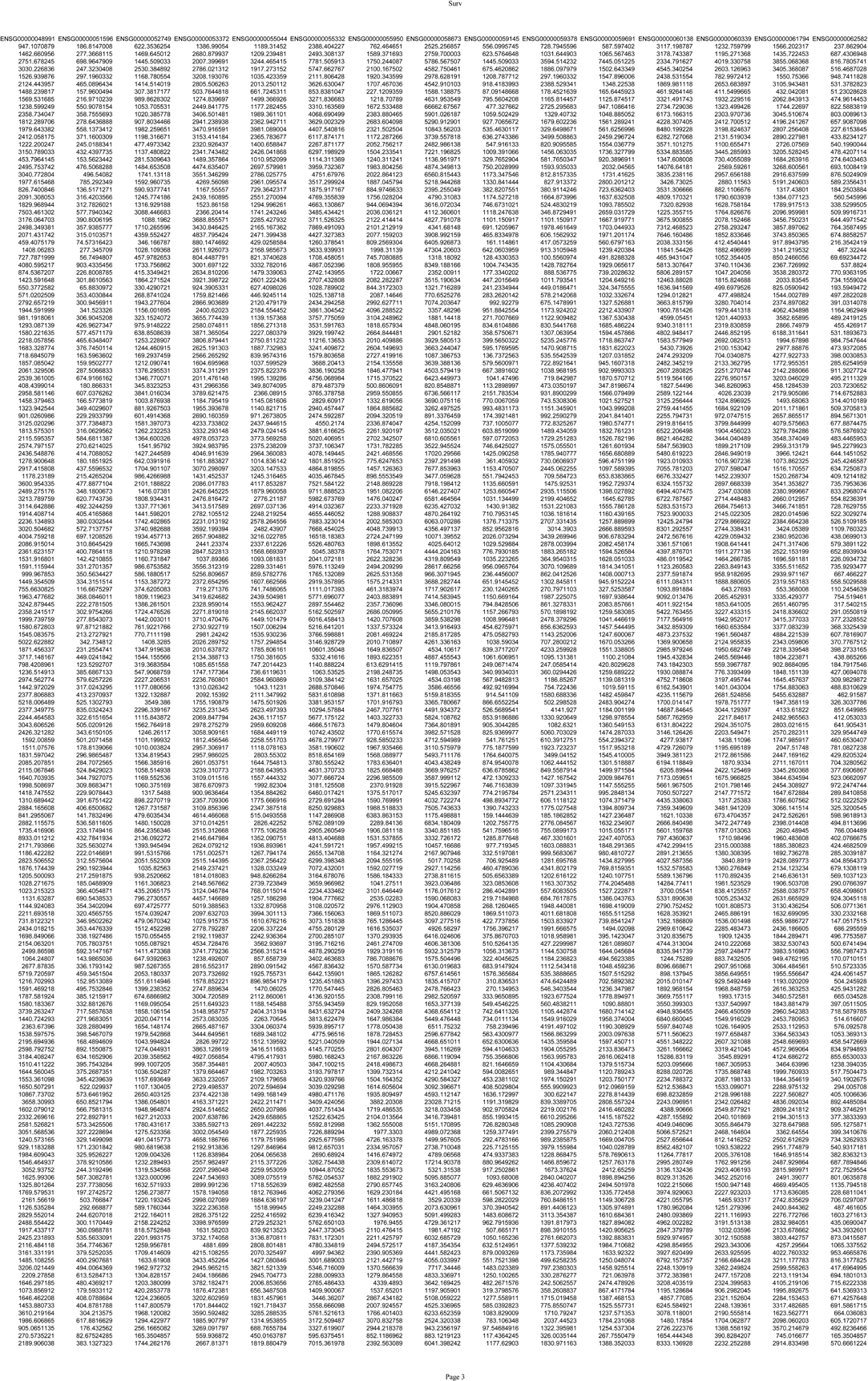
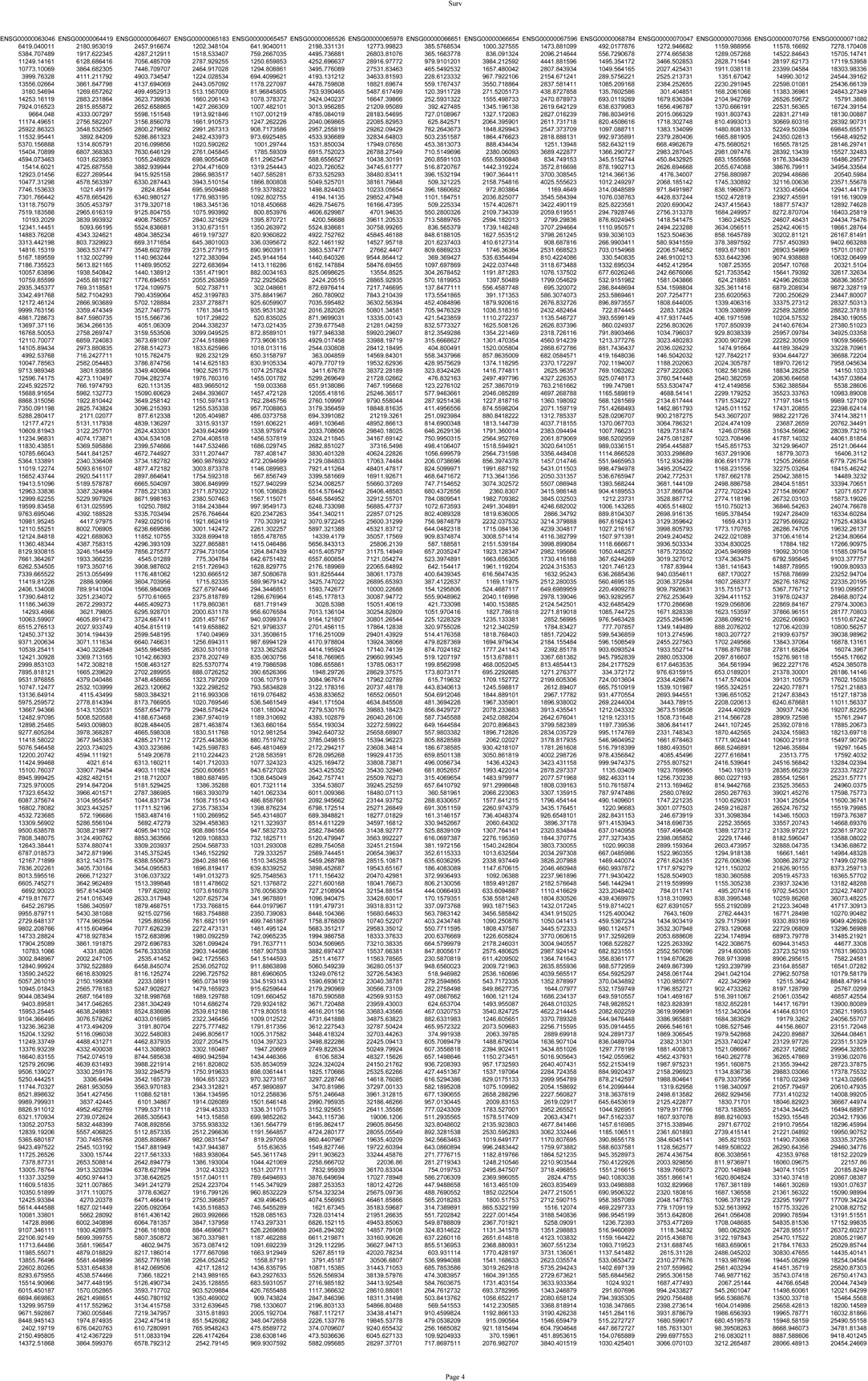
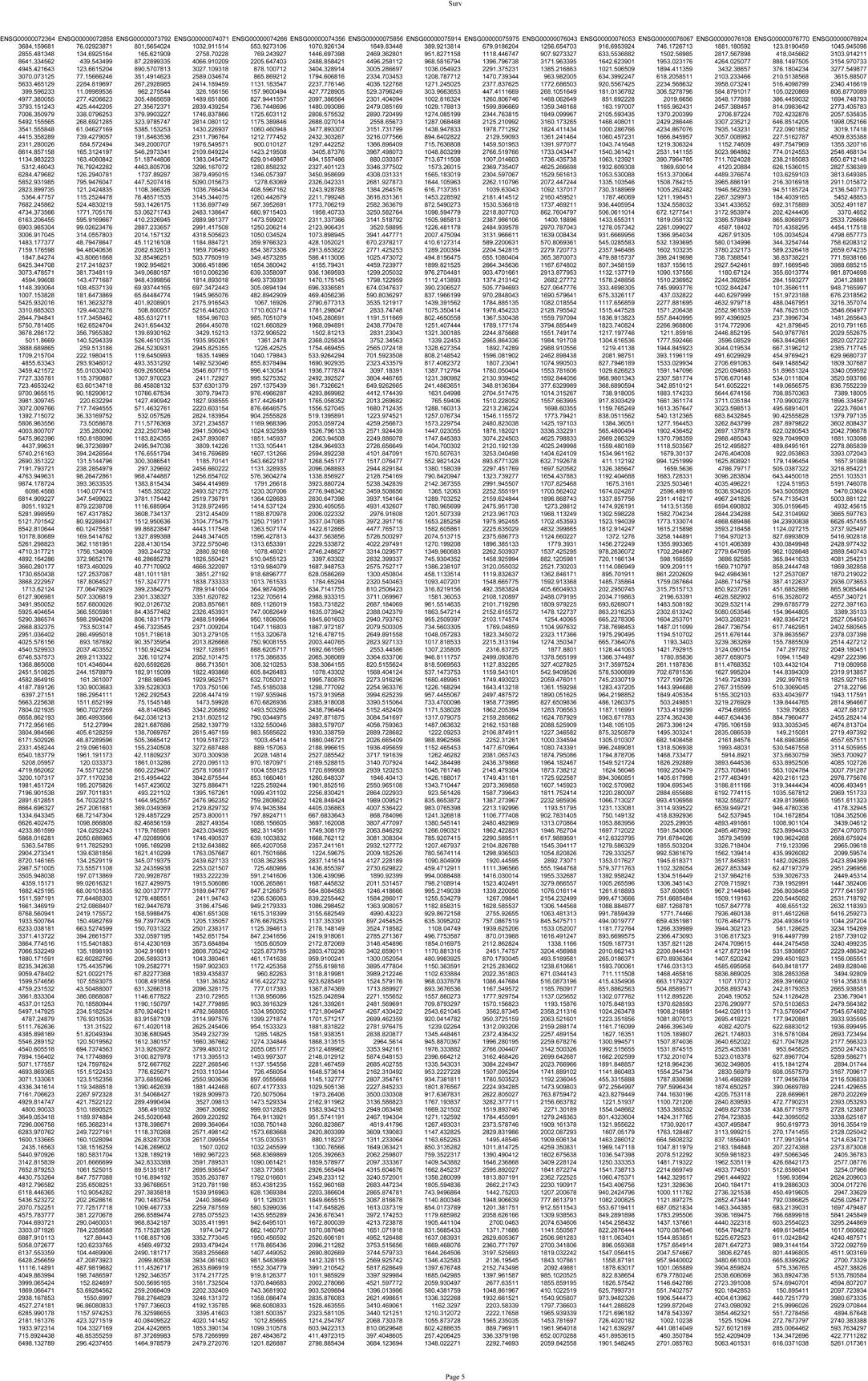
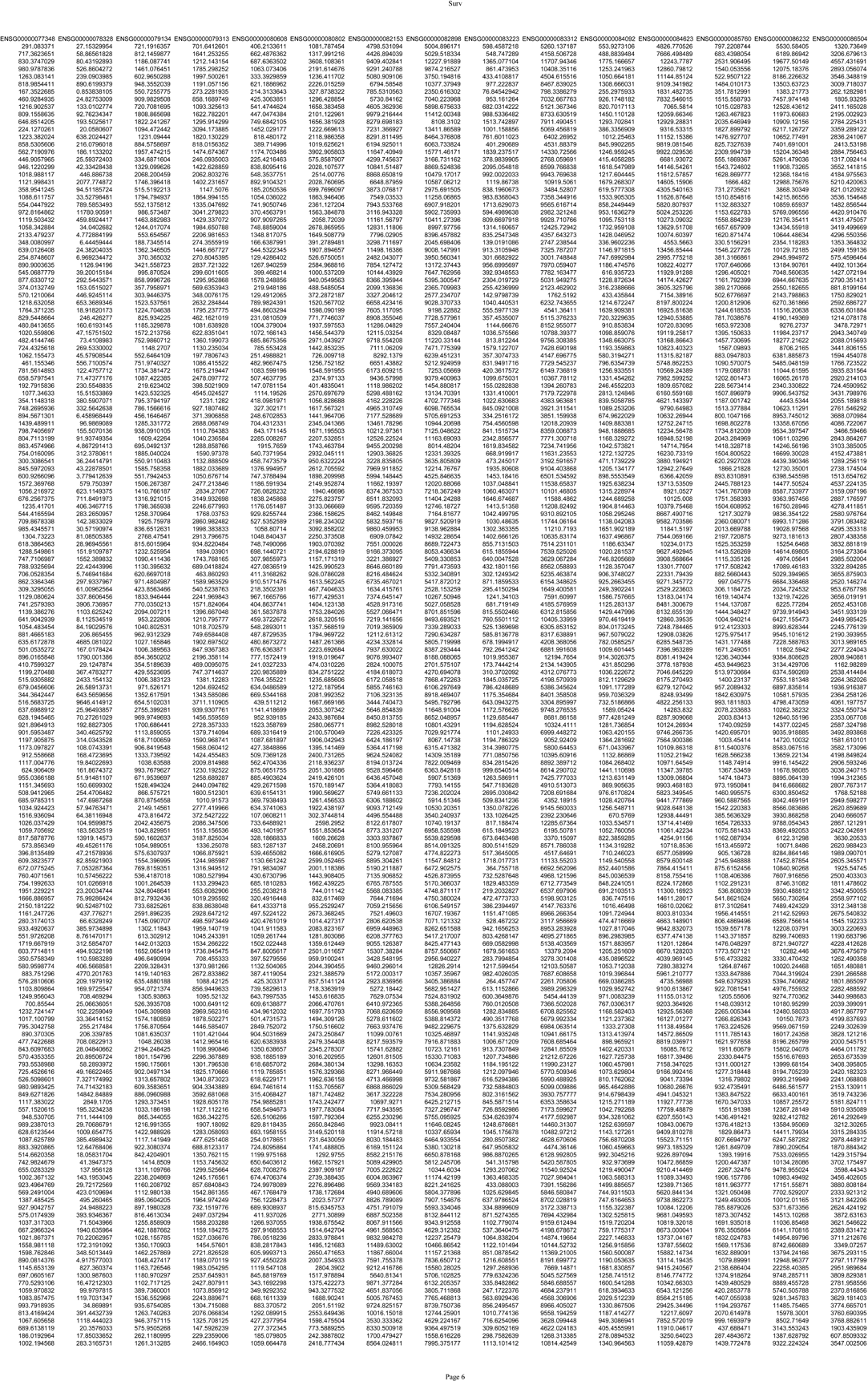
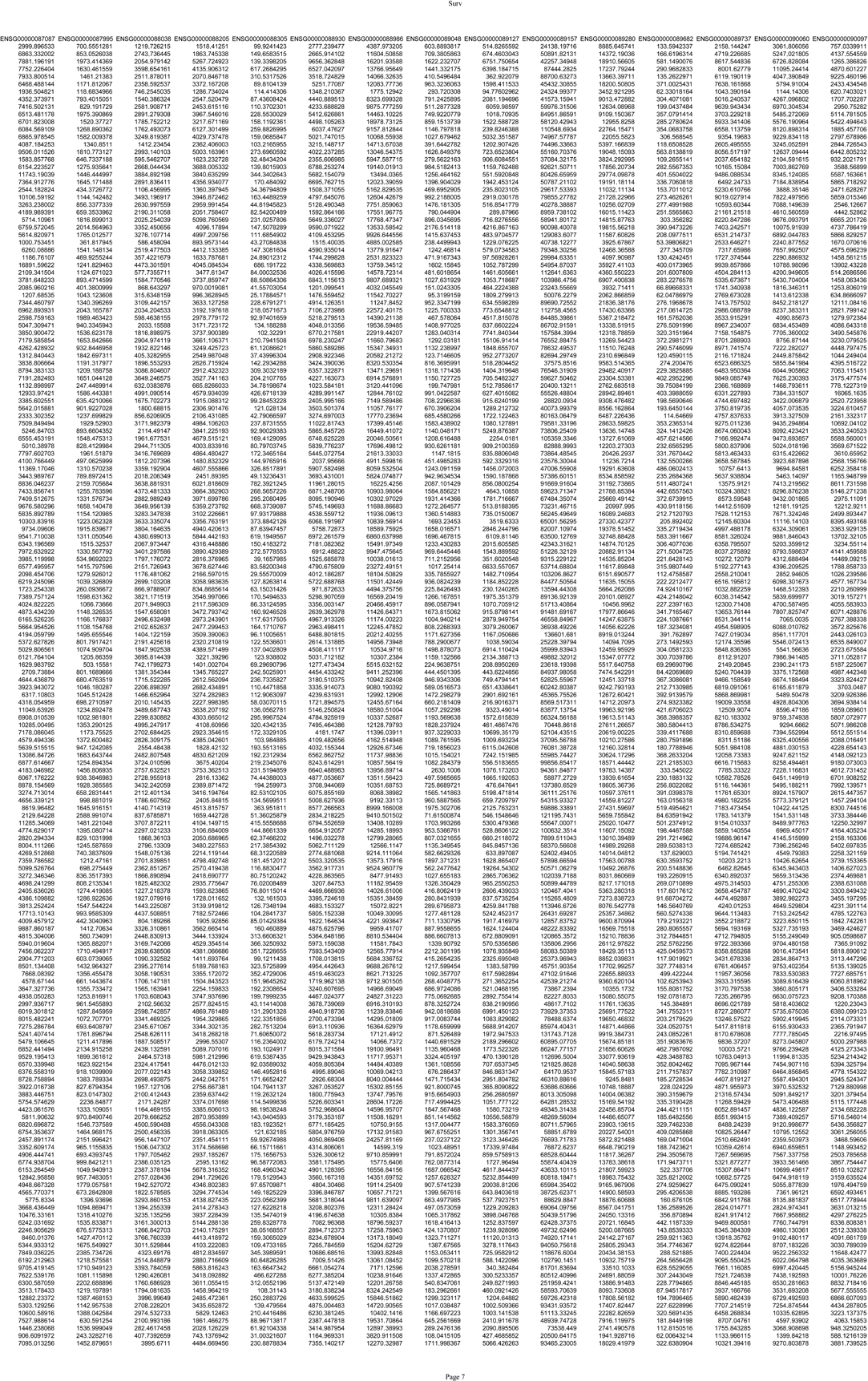
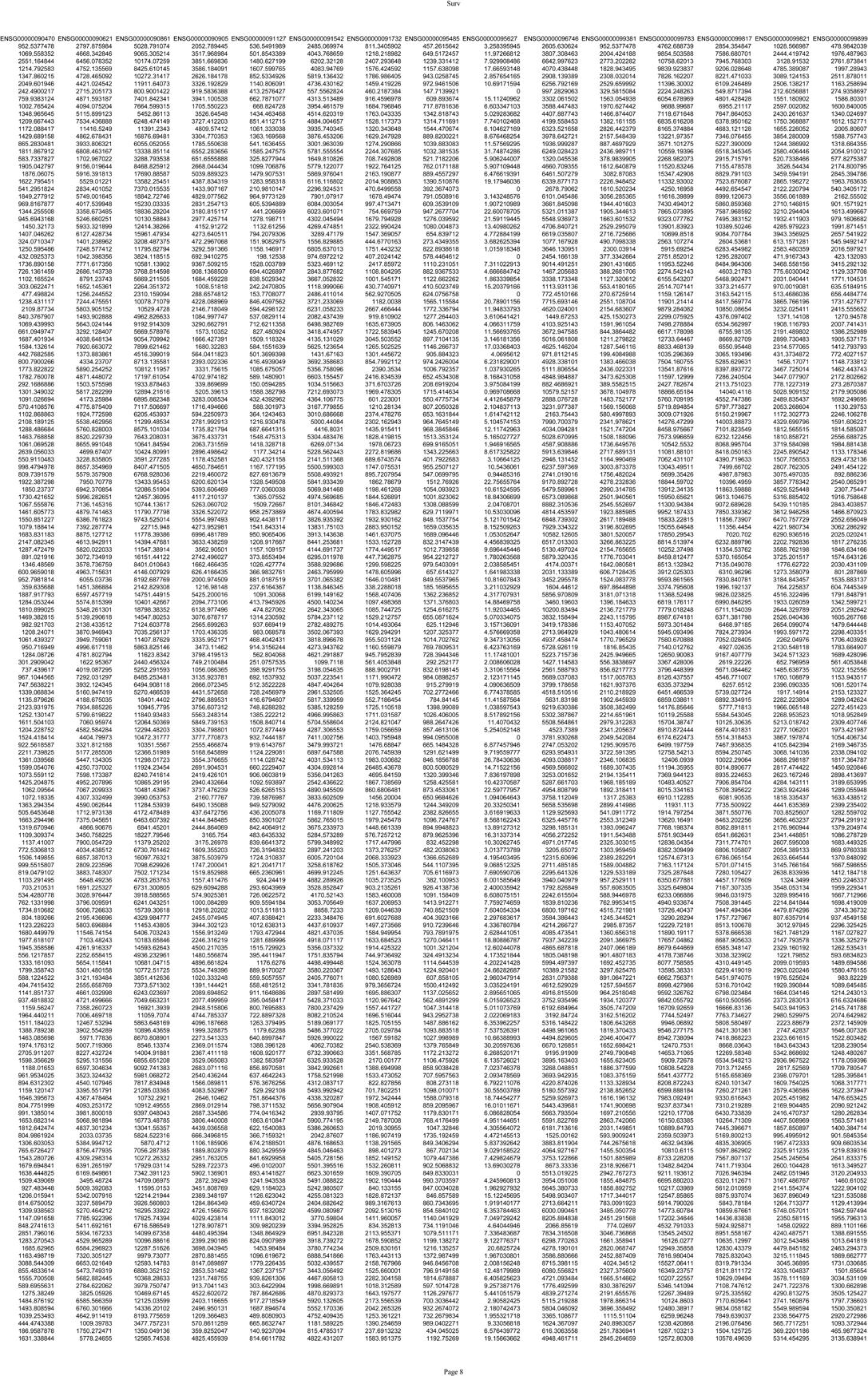
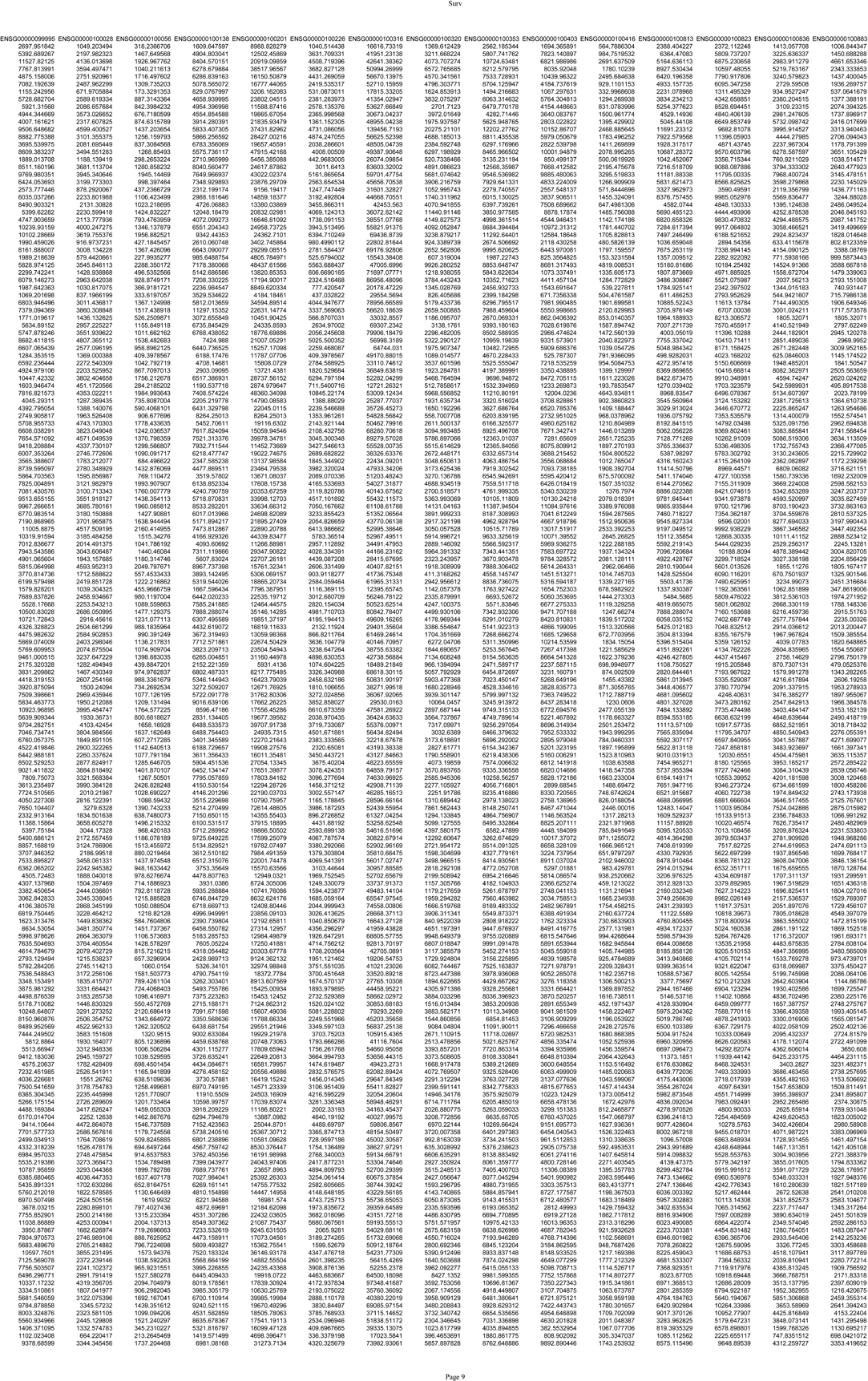
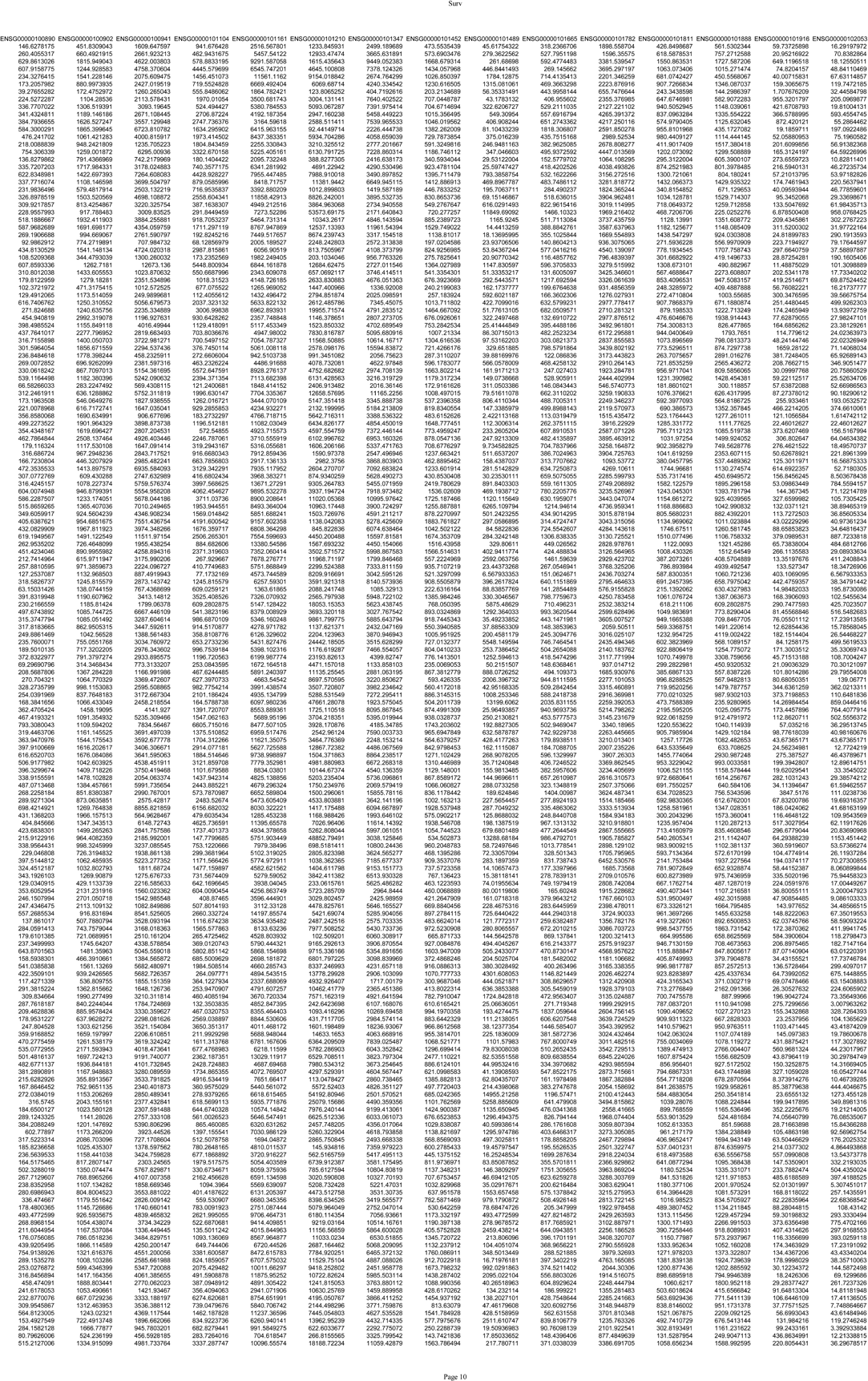
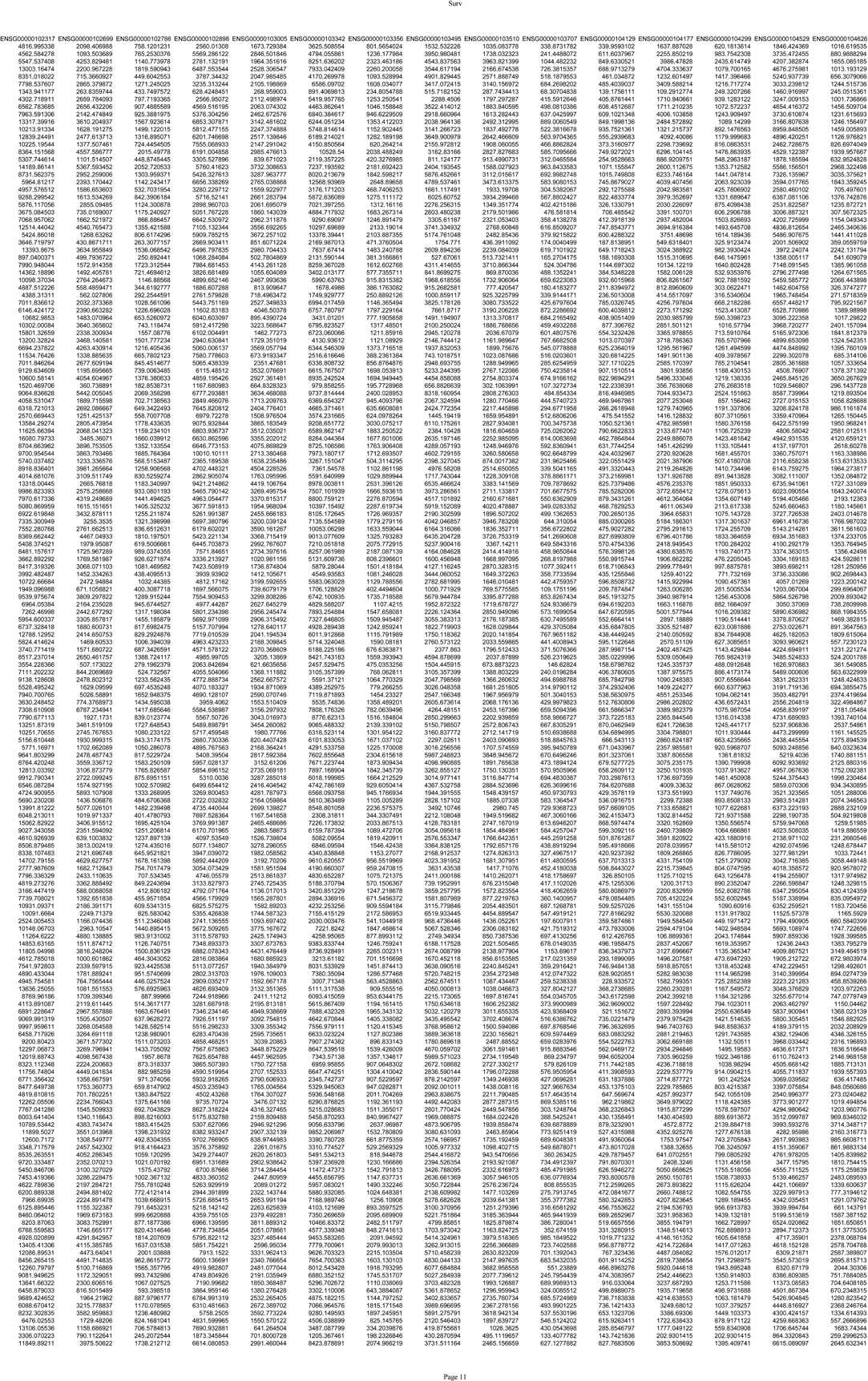
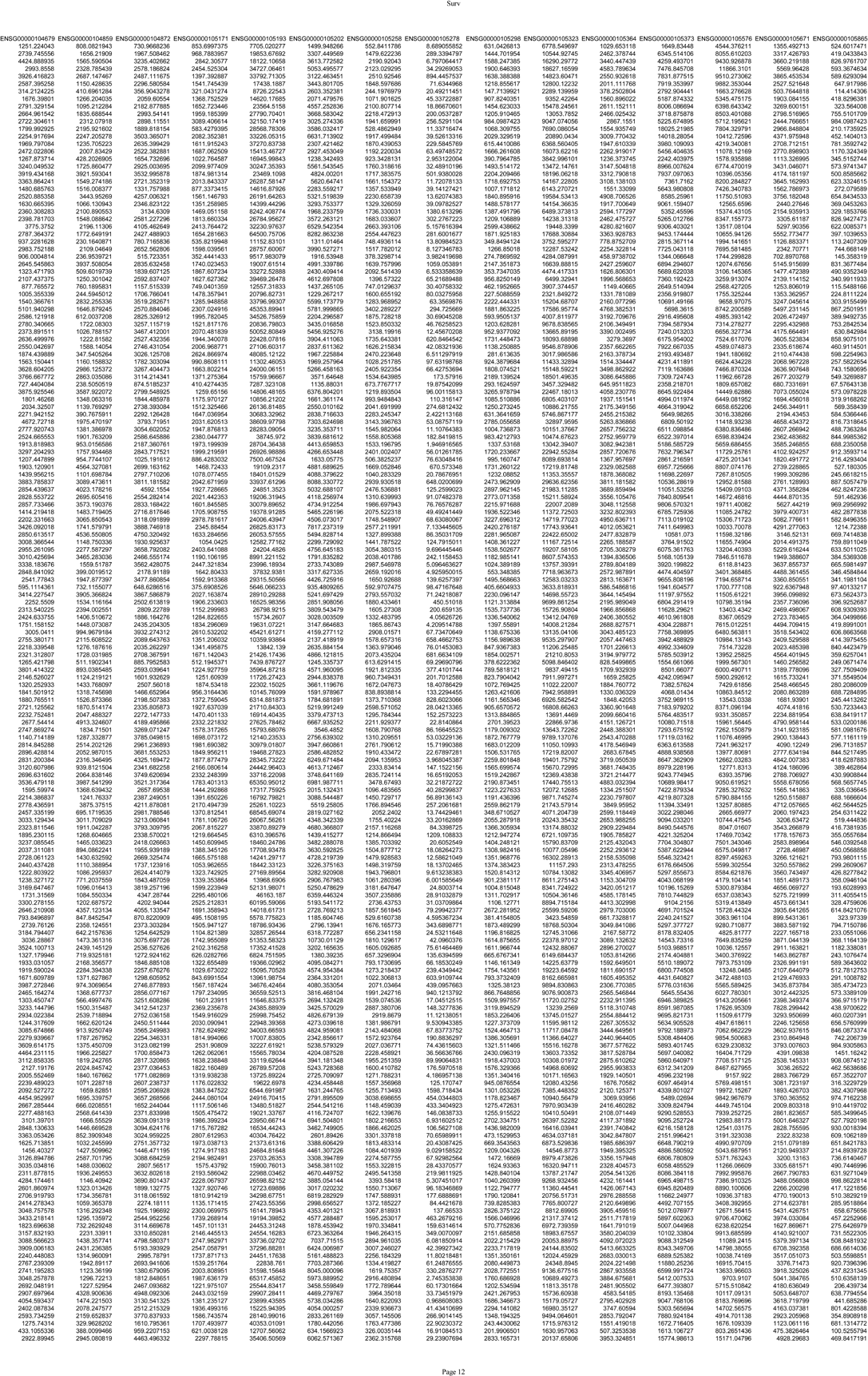
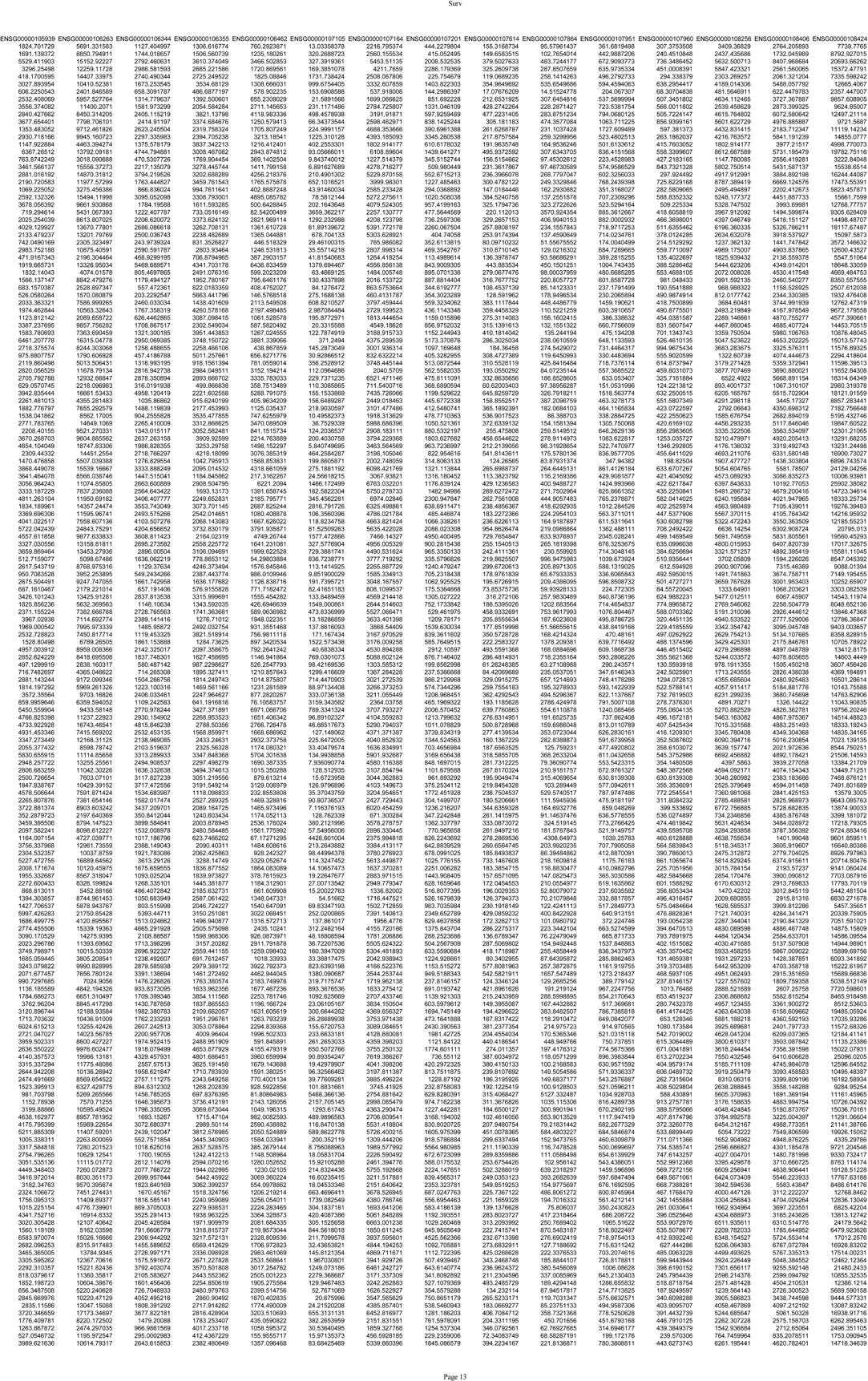
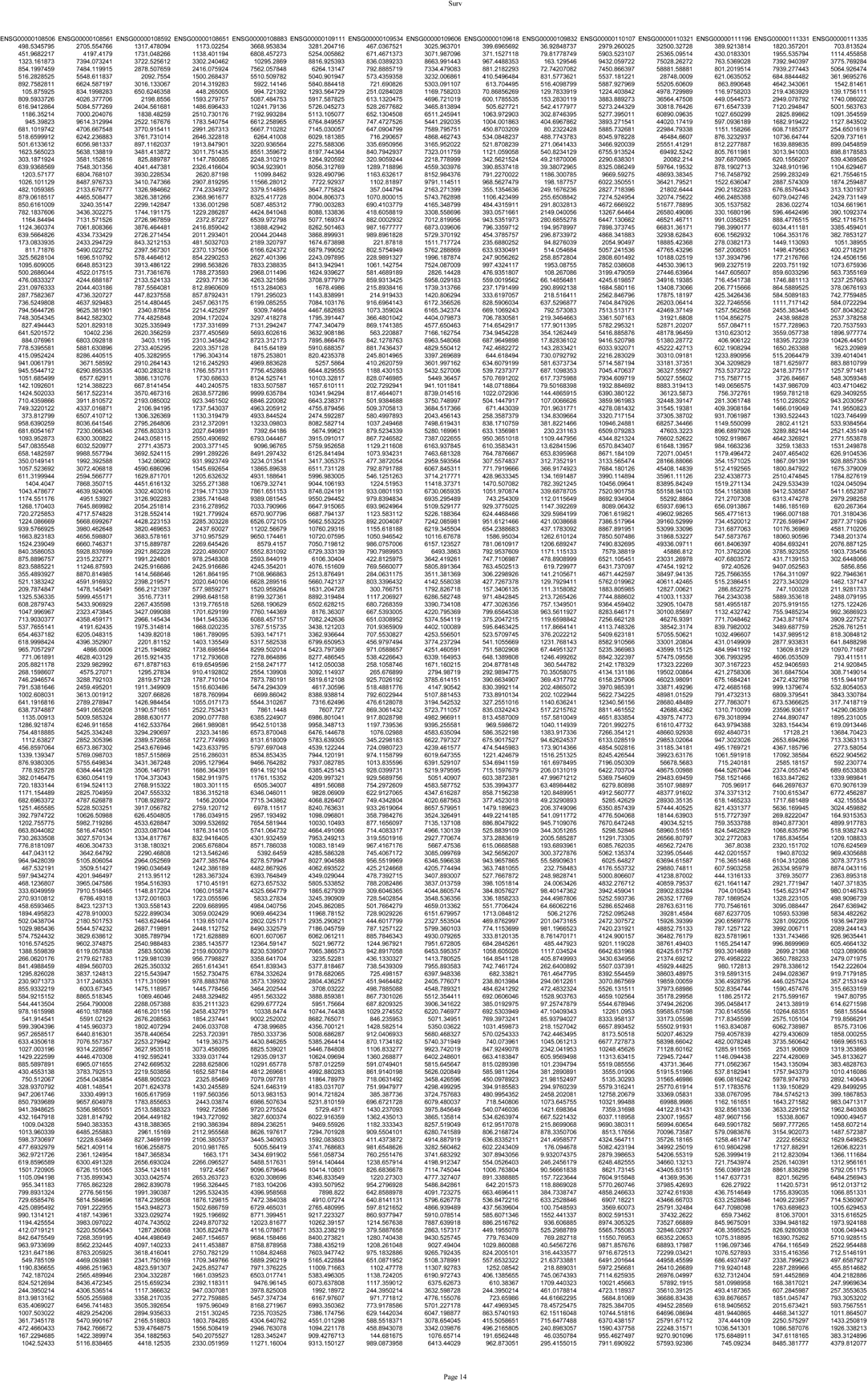
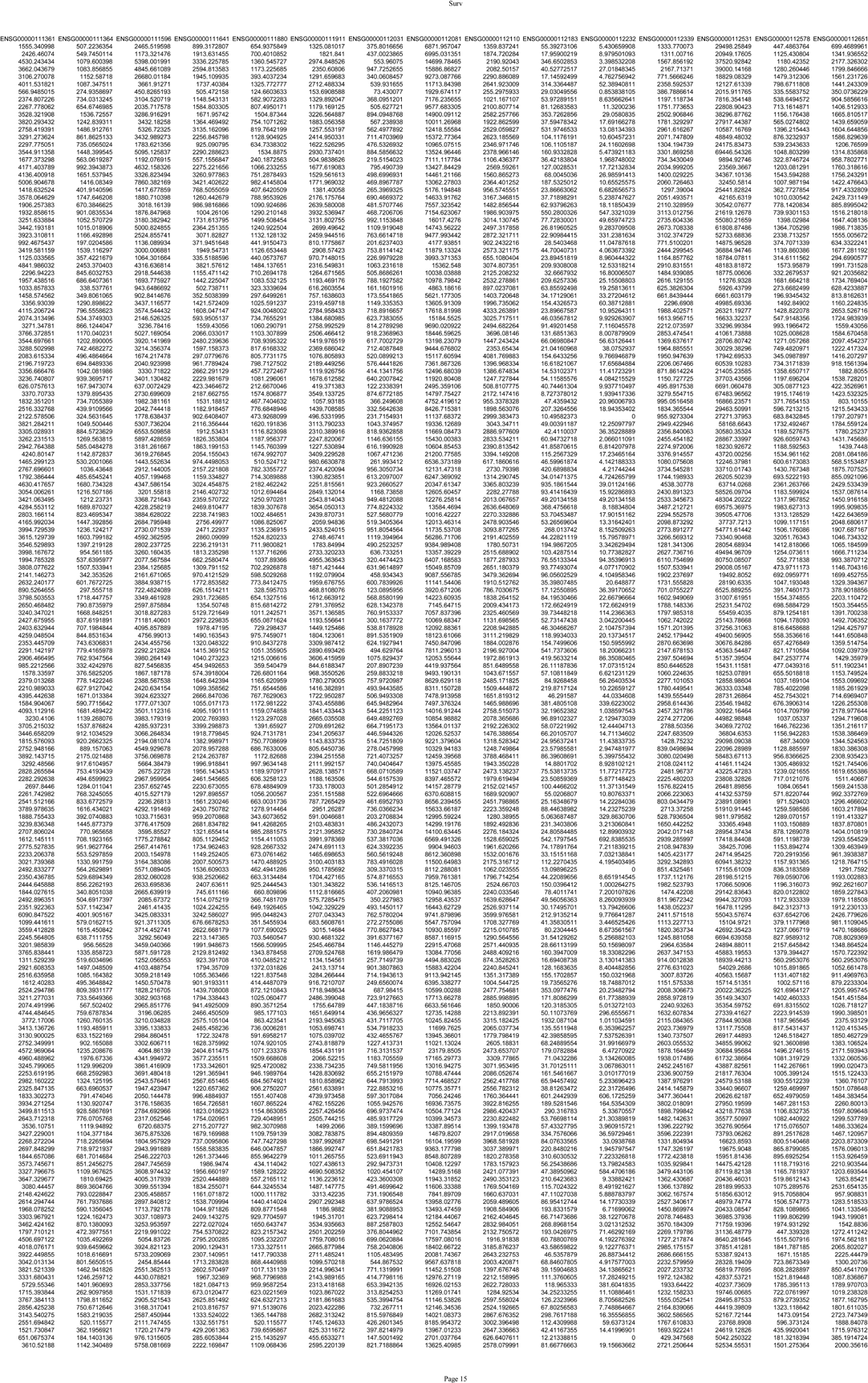
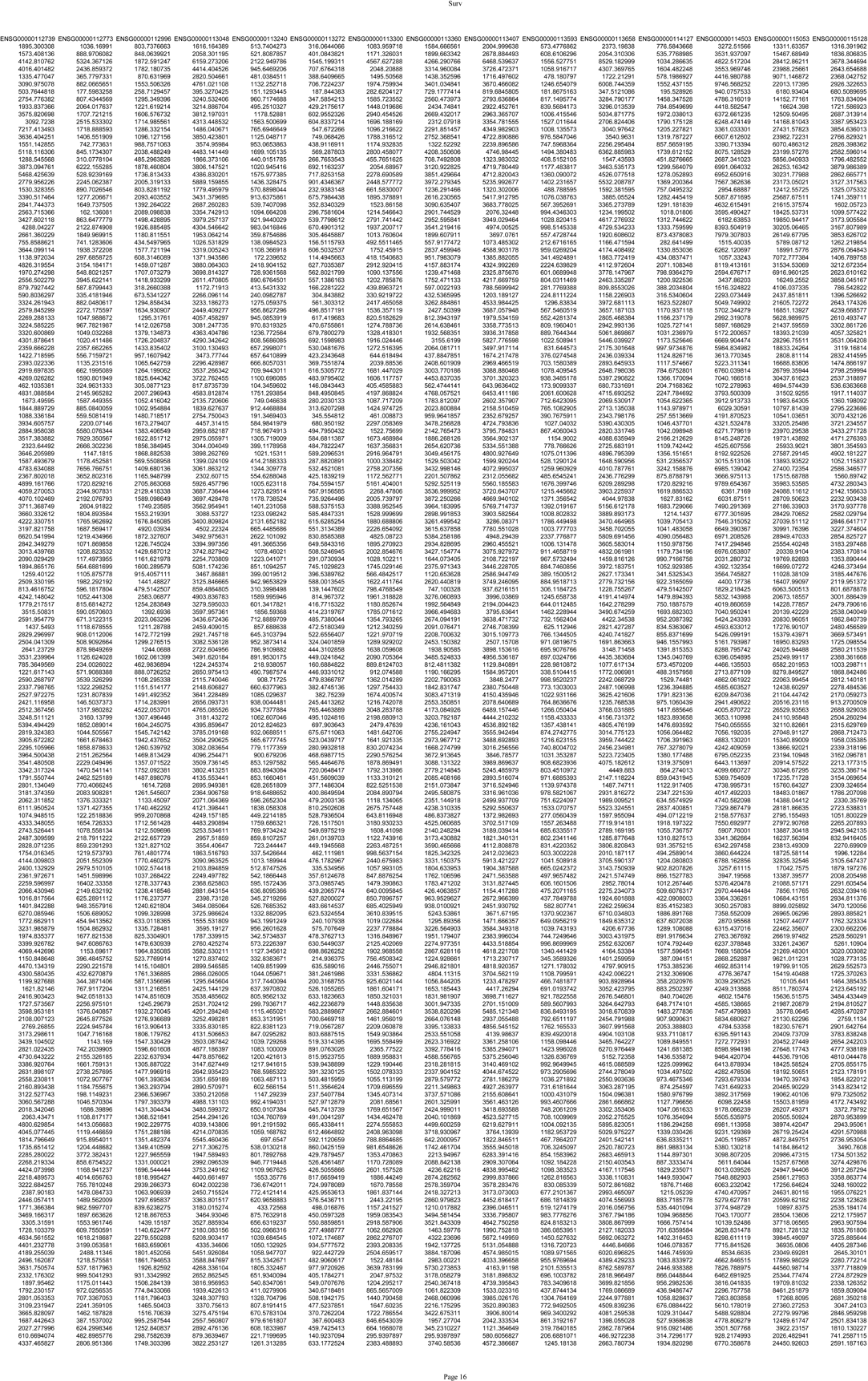
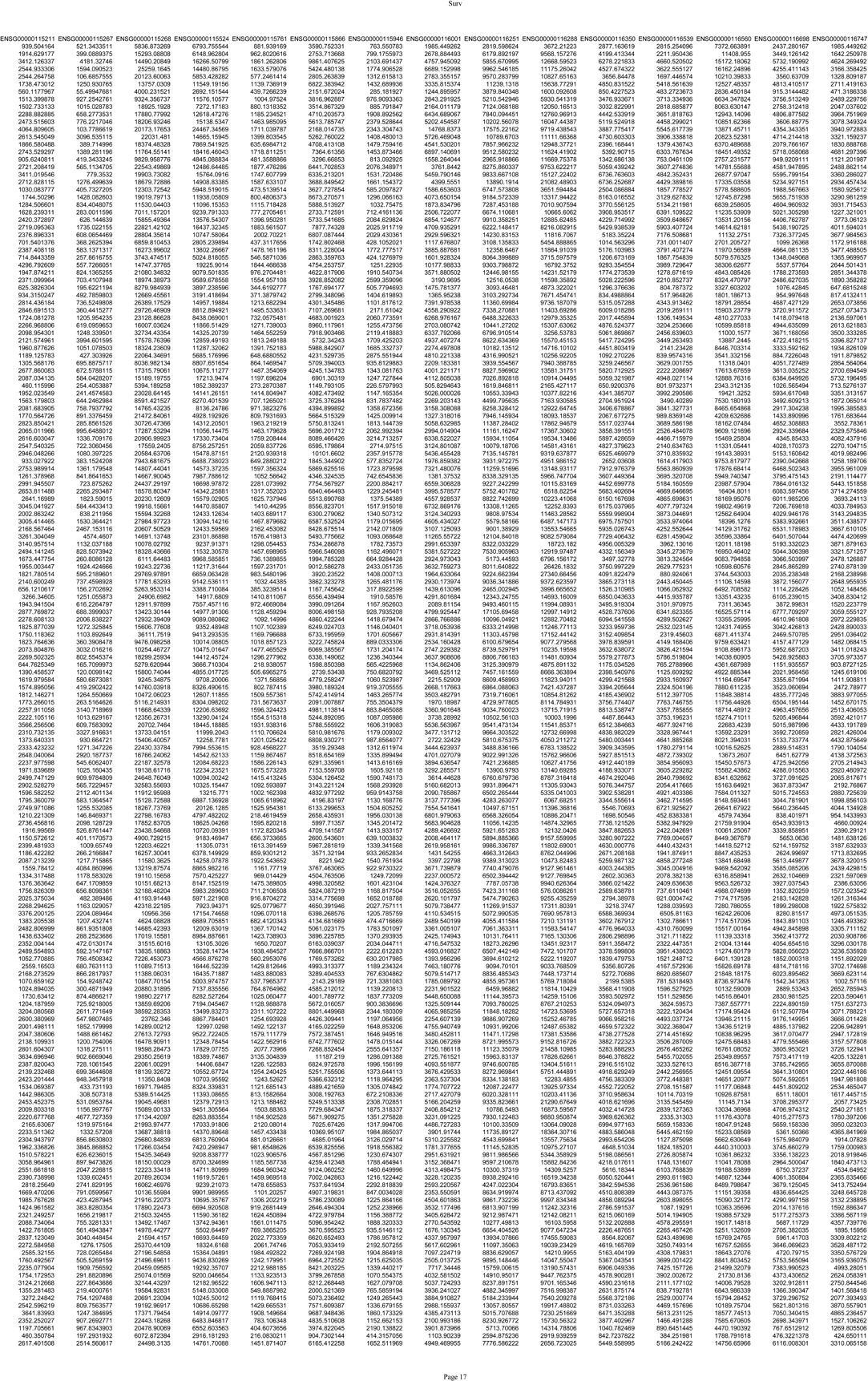
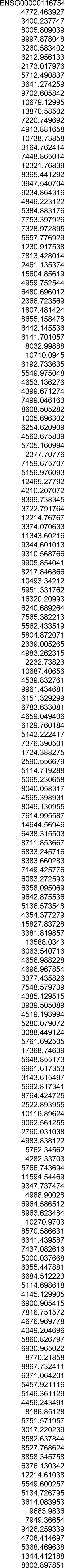
Differentially expressed RBPs used for survival analysis.

**Supplementary Table 9.**
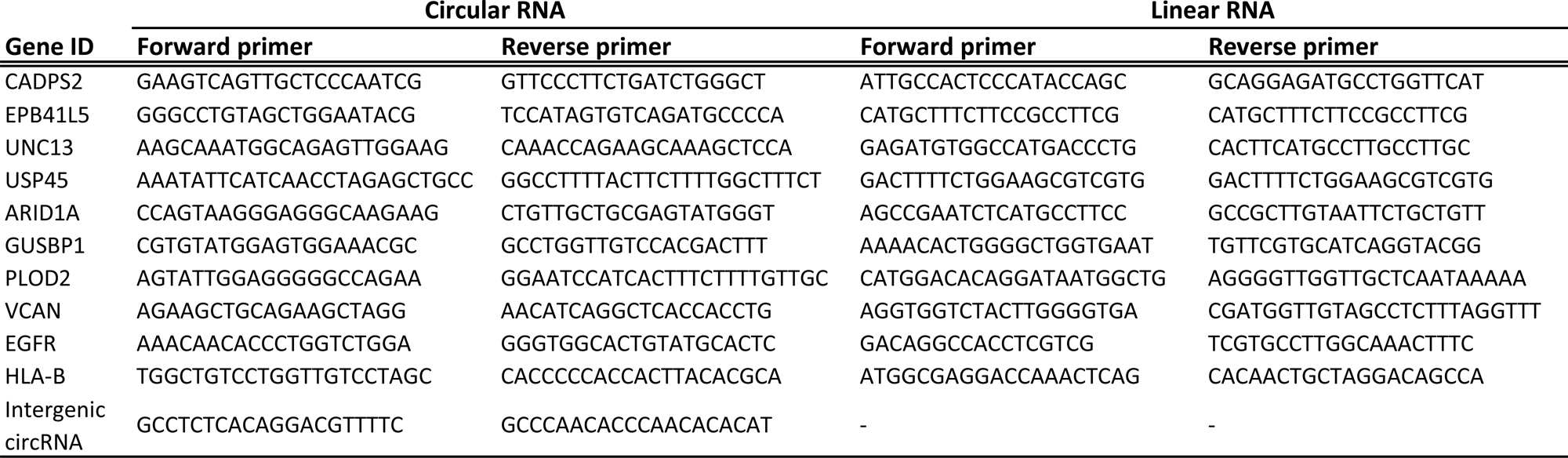
List of the primers used in the qPCR validation and RNase R treatment experiment.

## Figures legends

**Supplementary Figure 1.**
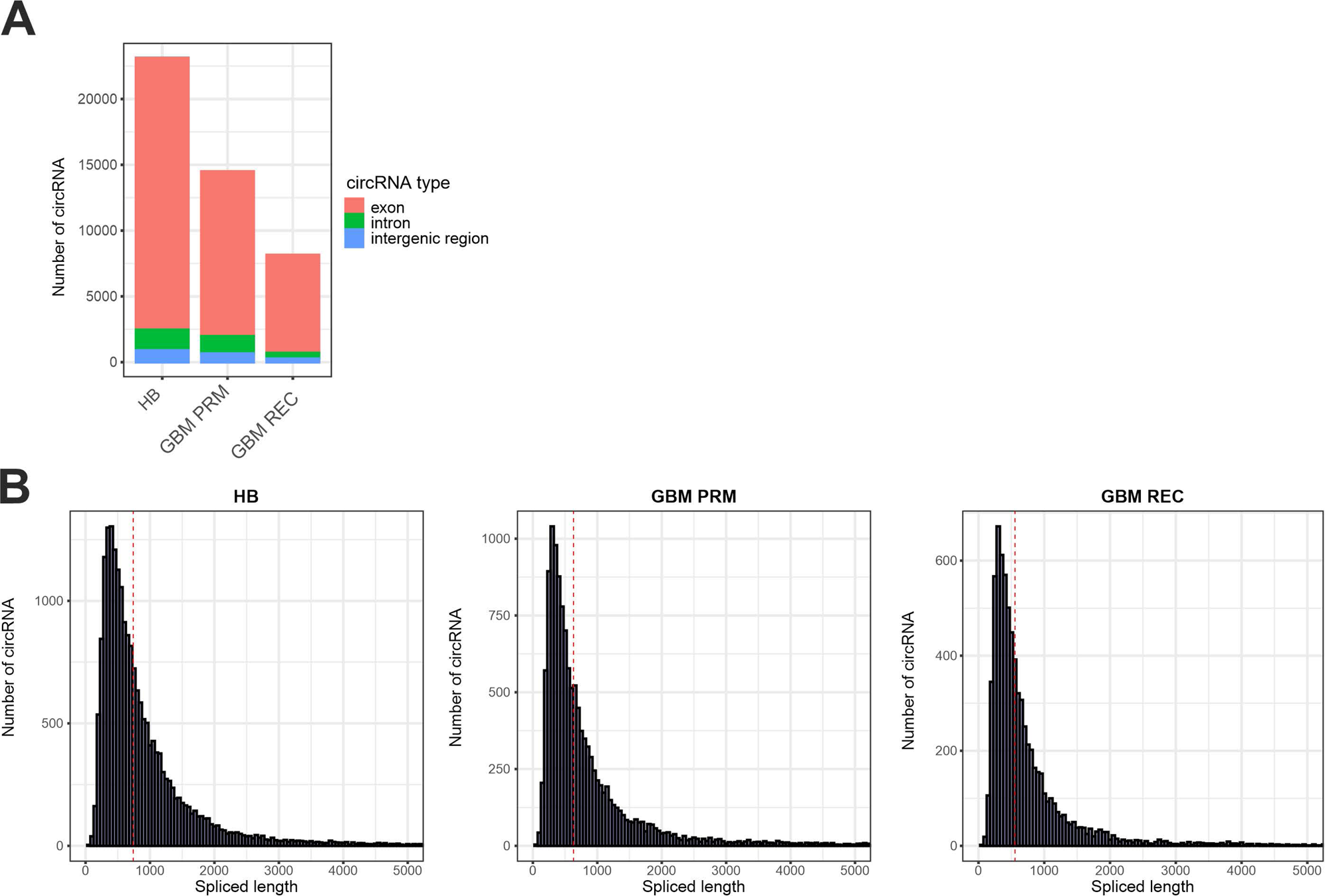
Overview of the features of circRNAs identified in GBM-PRM and GBM-REC tissues compared to HB control. **A.** Barplot presents the genomic origin of identified circRNAs from which the vast majority are derived from exons. **B.** Histogram displays a spliced length distribution of identified circRNAs for analyzed tissues.

**Supplementary Figure 2.**
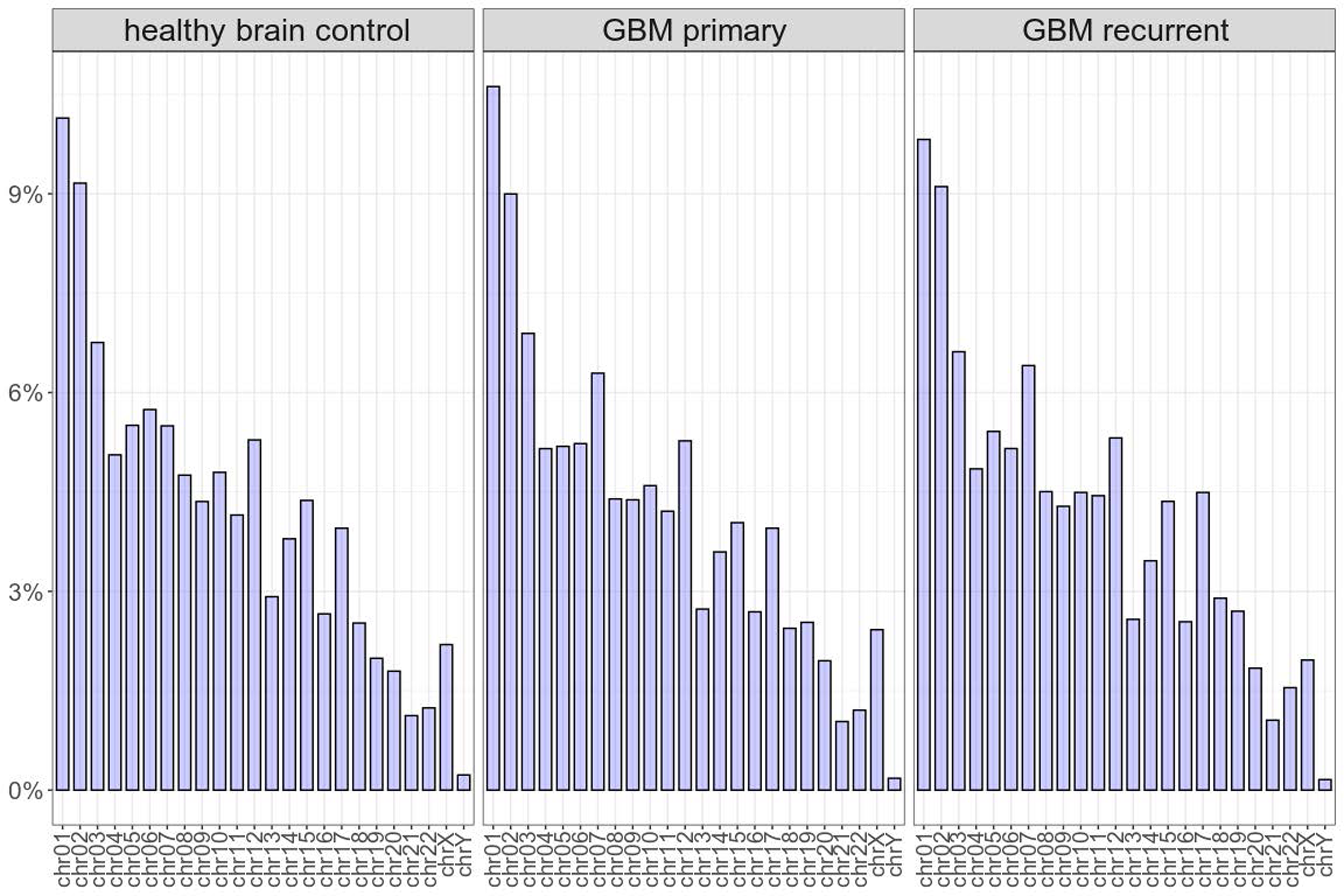
Percentage of unique circRNA per chromosome in GBM-PRM and GBM-REC in comparison to the HB control.

**Supplementary Figure 3.**
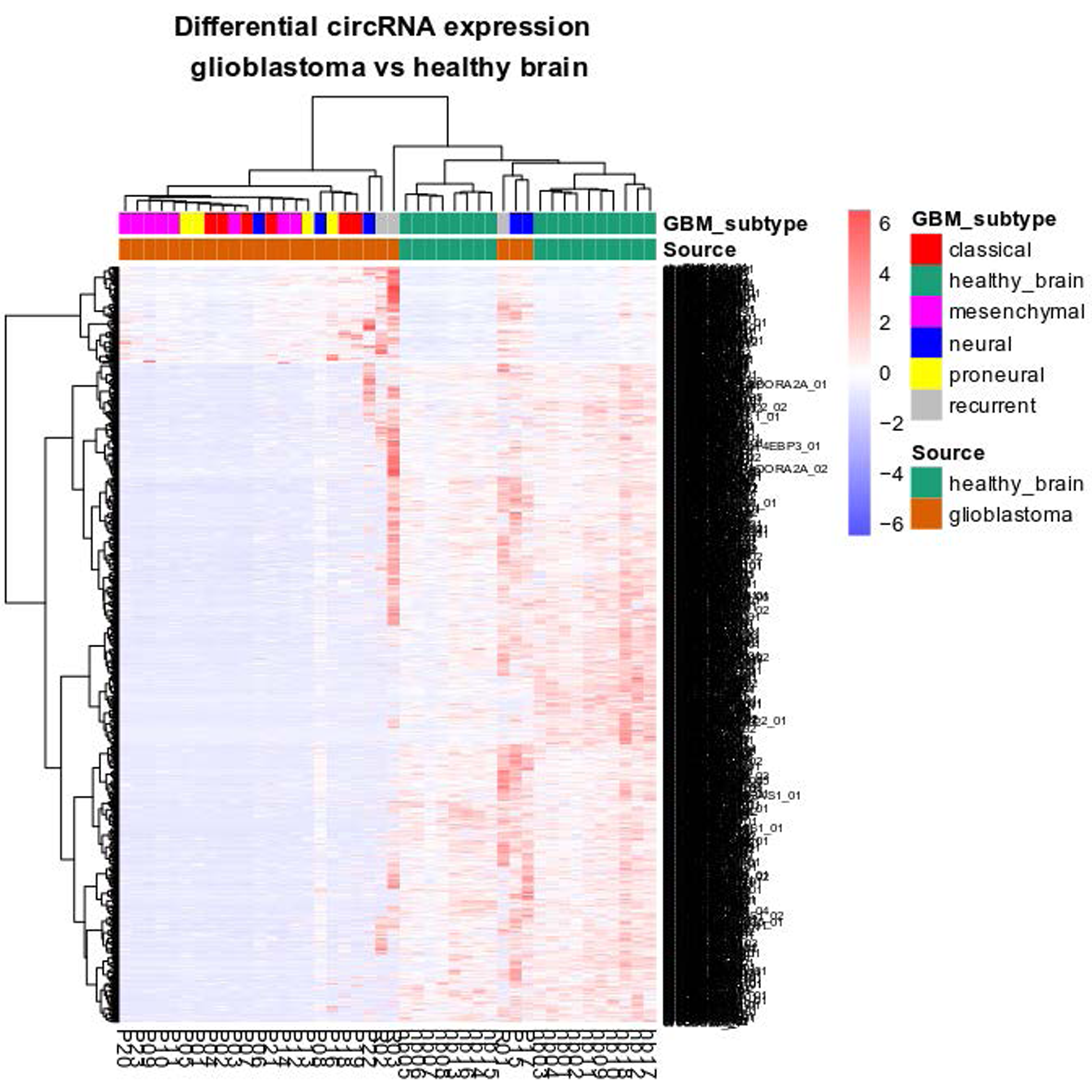
Cluster heatmap illustrating differential expression of circRNAs among GBM (GBM-PRM and GBM-REC) versus HB samples with the presence of different circRNAs in individual molecular GBM subtypes.

**Supplementary Figure 4.**
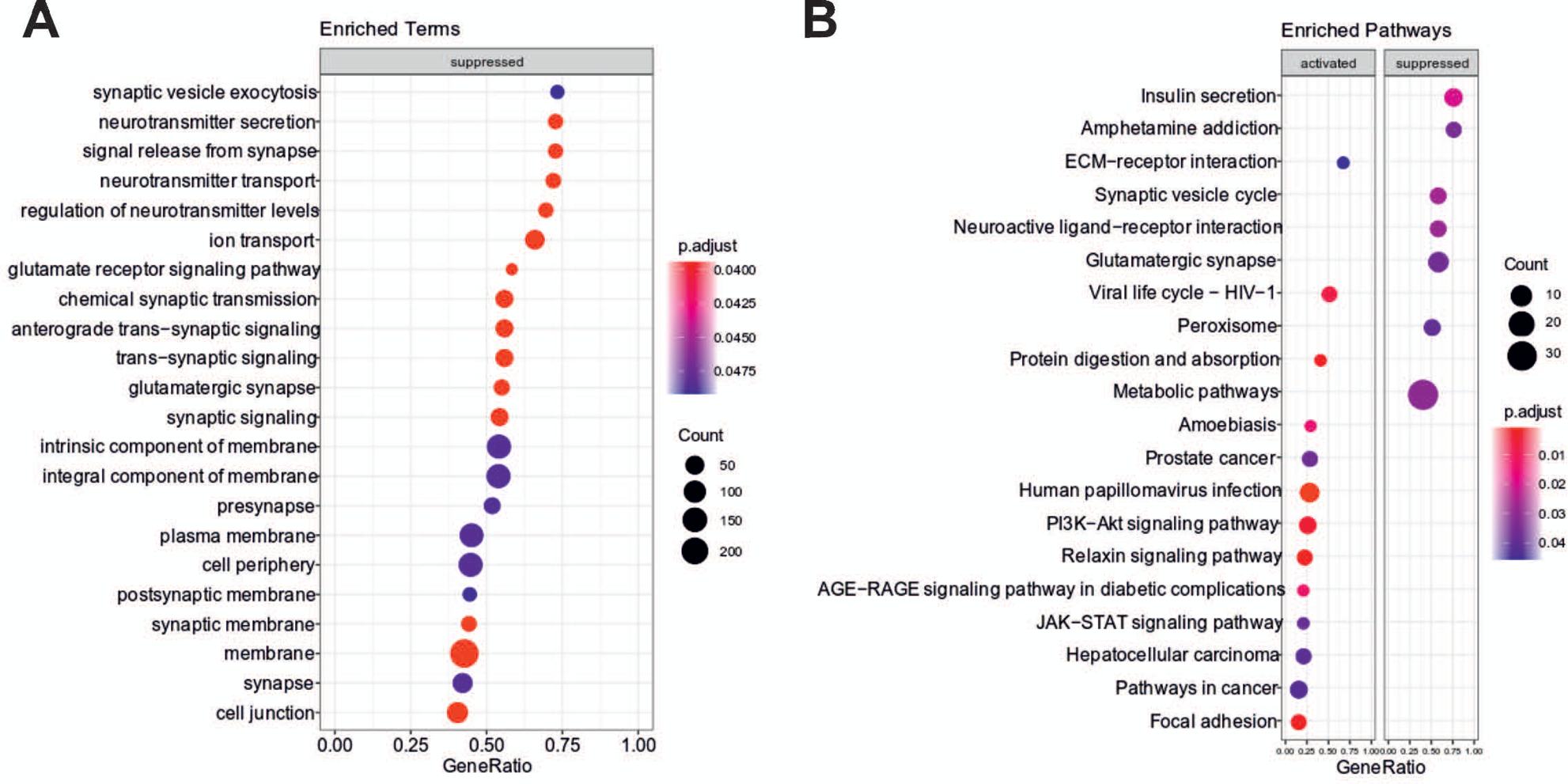
**A.** GO ontologies. Enrichment was calculated using mean circRNA log2 fold change for the particular host genes. **B.** Bubble map of KEGG enriched terms using GO ontologies. Enrichment was calculated using mean circRNA log2 fold change for the particular host genes.

**Supplementary Figure 5.**
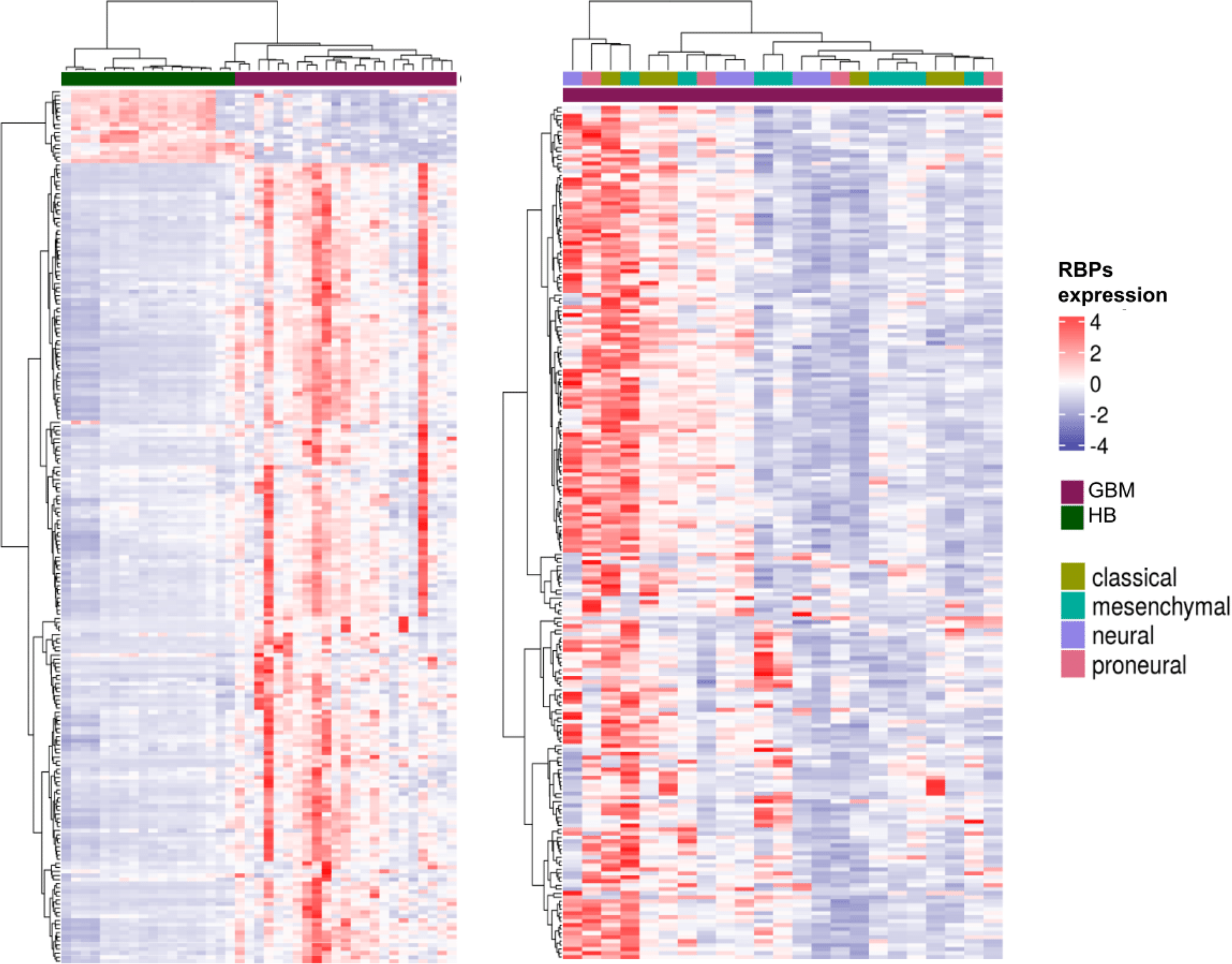
Cluster heatmap illustrating differential expression of RBPs among GBM (GBM-PRM and GBM-REC) versus HB samples. Expression of 214 RBPs can be used to define novel patients’ subgroups beyond known molecular subtypes.

**Supplementary Figure 6.**
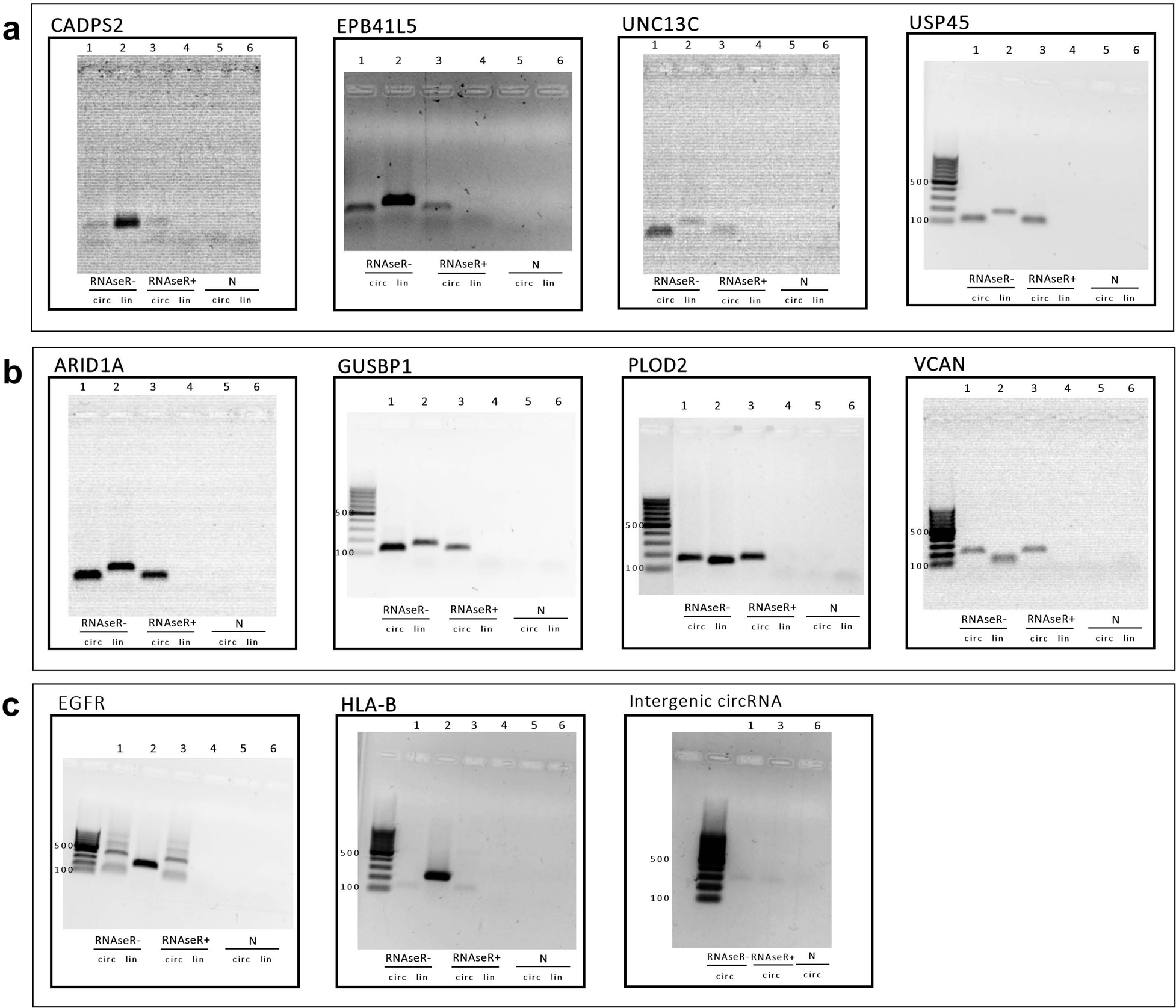
RNase R treatment of circRNAs and their linear counterparts. A-C. Bands represent PCR products on agarose gel of selected downregulated, upregulated and progression-related candidates, respectively. **1.** Band represent PCR product on agarose gel of circRNA without RNase R treatment; **2.** Band represent PCR product on agarose gel of mRNA without RNase R treatment; **3.** Band represent PCR product on agarose gel of circRNA with RNase R treatment; **4.** Band represent PCR product on agarose gel of mRNA with RNase R treatment; **5, 6.** Negative control of PCR product.

**Supplementary Figure 7.**
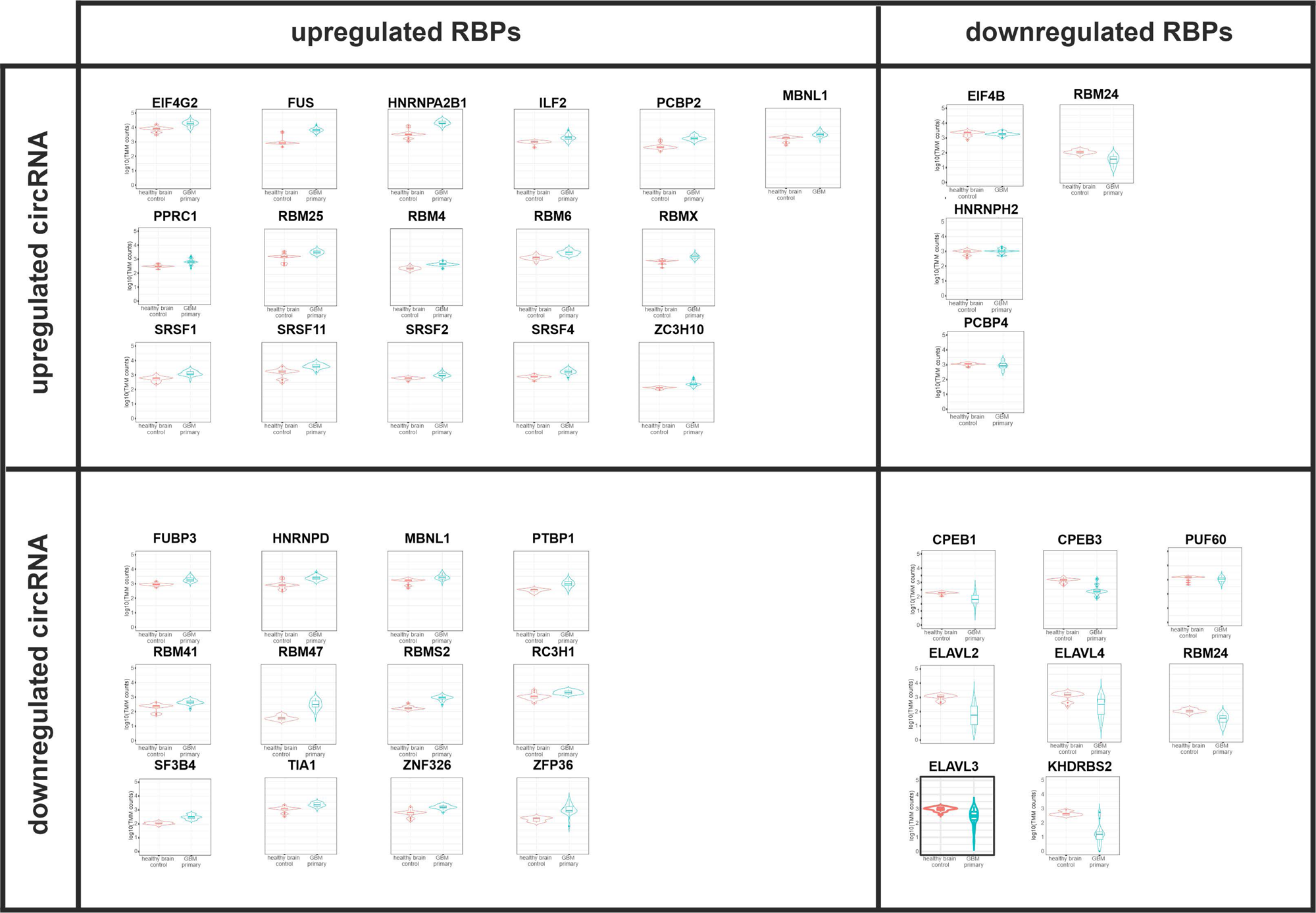
Normalized expression distribution of RBP’s transcripts in HB and GBM. **A.** Upregulated RBPs with motifs enriched in upregulated circRNAs. **B.** Downregulated RBPs with motifs enriched in upregulated circRNAs. **C.** Upregulated RBPs with motifs enriched in downregulated circRNAs. **D.** Downregulated RBPs with motifs enriched in downregulated circRNAs. Sequence logos represent motifs enriched in circRNAs.

**Supplementary Figure 8.**
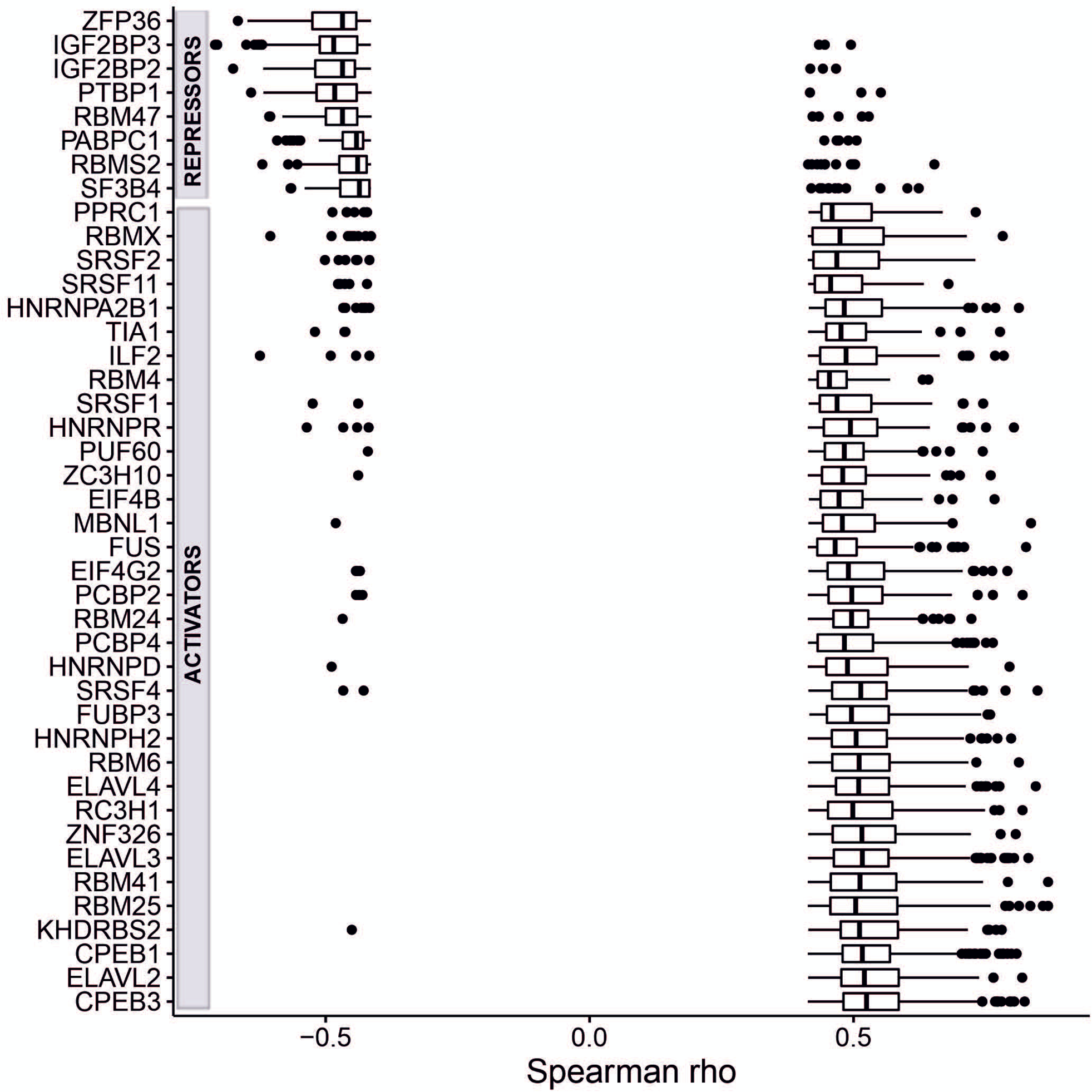
Boxplots representing correlation coefficient between RBPs with motifs enriched in circRNAs and differentially expressed cicrRNAs.

**Supplementary Figure 9.**
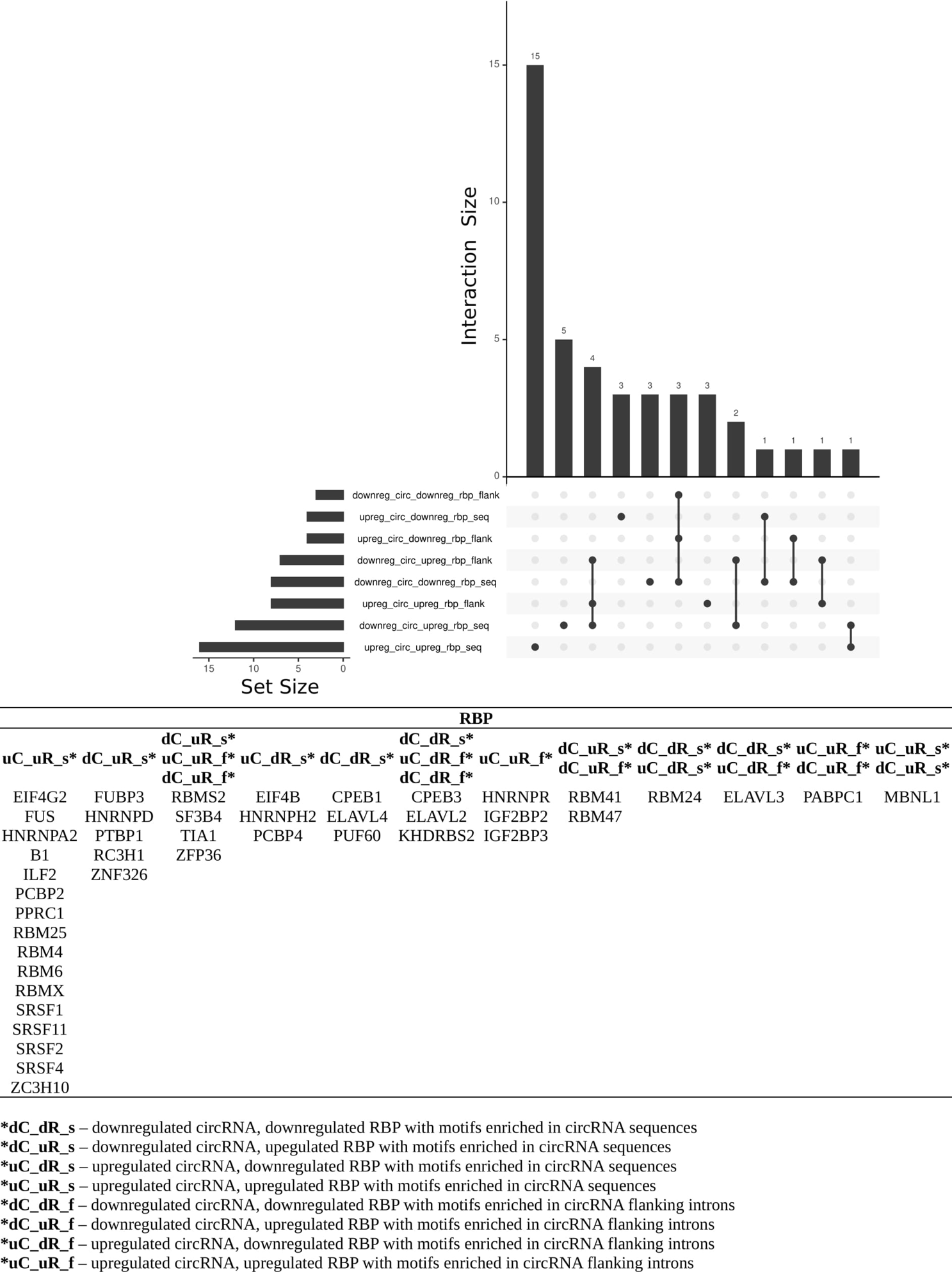
Upset plot for RBPs with binding motifs in flanking introns vs RBPs with binding motifs in circRNAs.

**Supplementary Figure 10.**
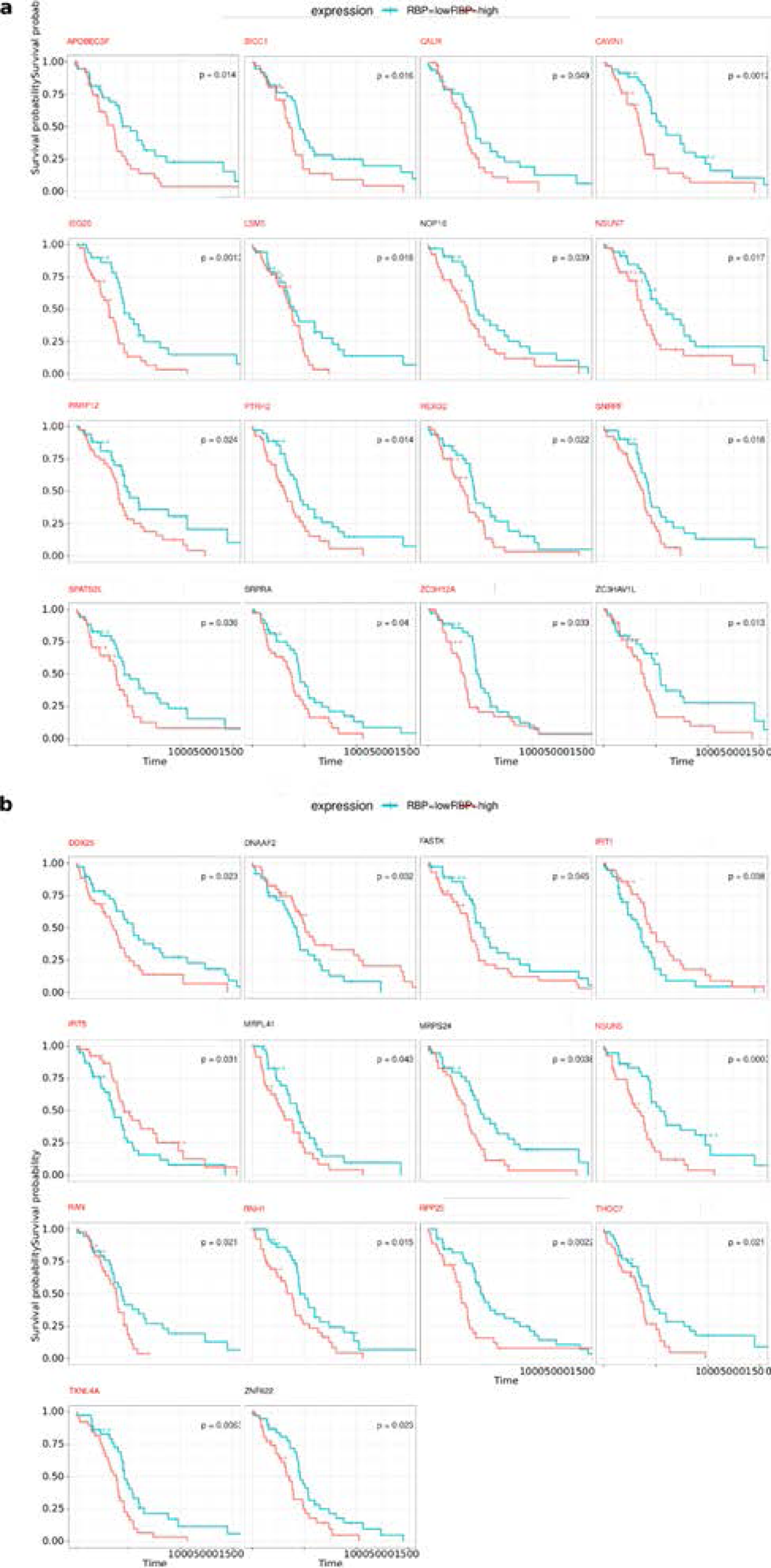
Survival analysis for differentially expressed RBPs based on TCGA glioblastoma dataset. **A.** Kaplan-Meier curves for RBPs upregulated in analyzed 23 GBM samples and significantly correlated with overall survival rate with TCGA GBM dataset. **B.** The same as A, for downregulated RBPs. RBPs highlighted red in both panels were previously shown to be brain tumor markers, correlated with disease outcome or response to treatment.

**Supplementary Figure 11.**
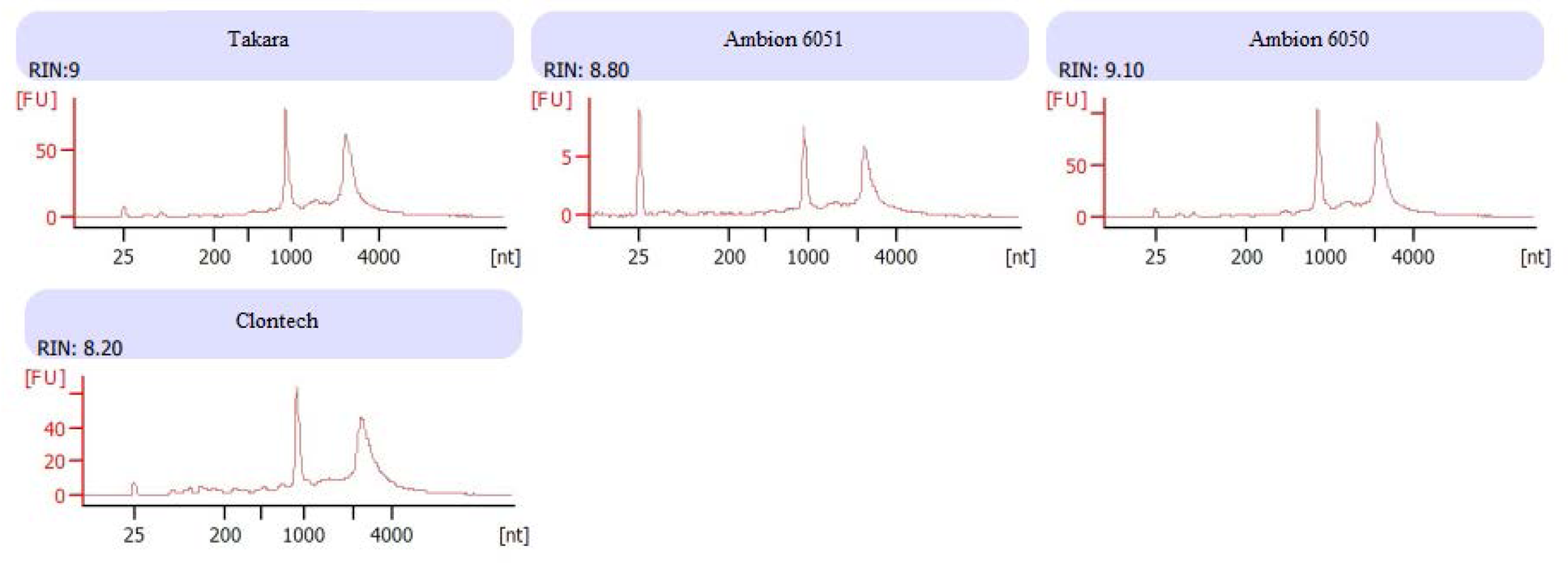
Electrophoretic separation of total RNA isolated from HB samples. The electrophoretic separation was prepared using the Agilent 2100 Bioanalyzer system. Only samples with RIN (RNA Integrity Number) value above 8 were taken into further investigation.

**Supplementary Figure 12.**
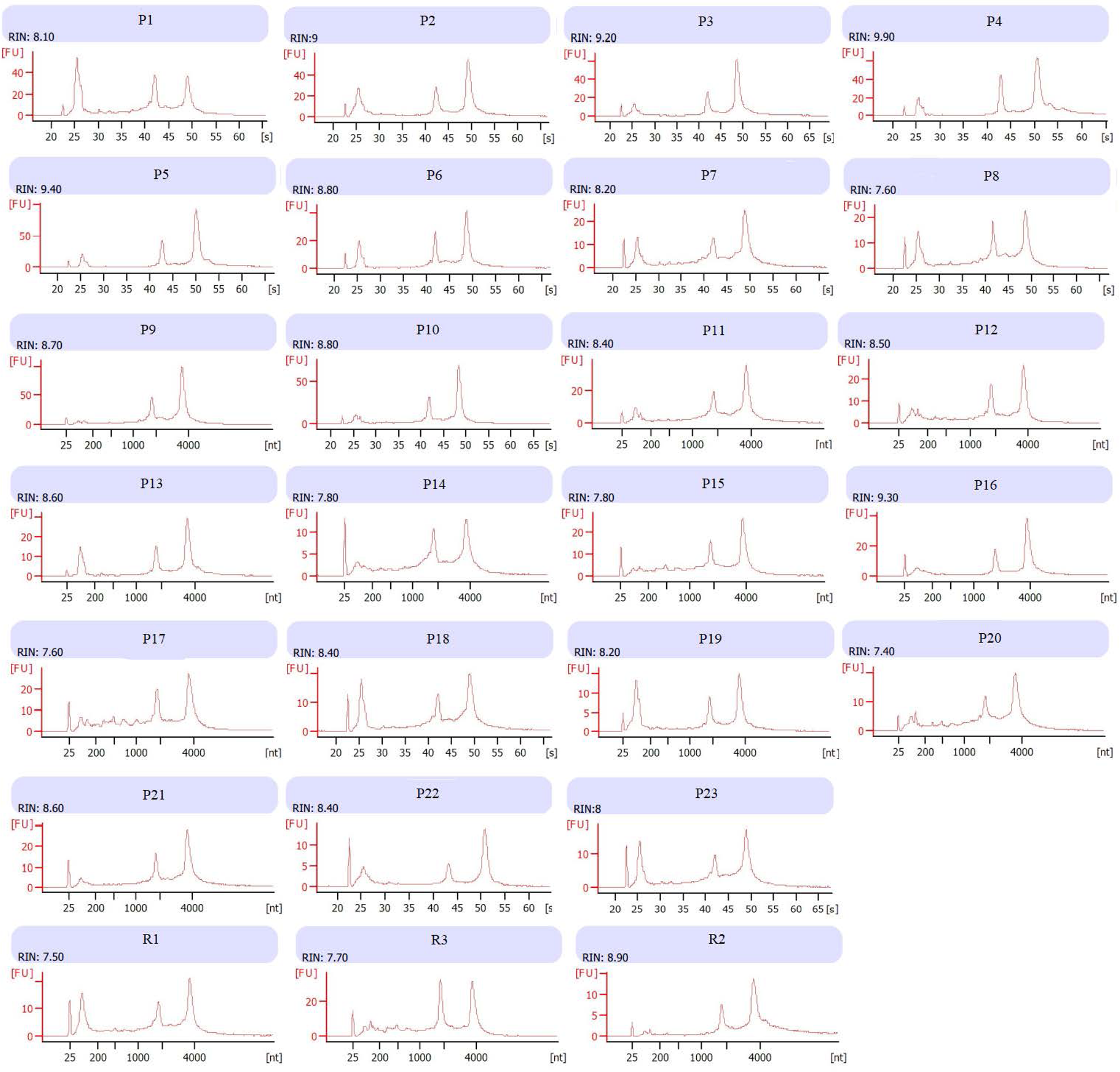
Electrophoretic separation of total RNA isolated from GBM-PRM and GBM-REC samples. Electrophoretic separation was prepared using the Agilent 2100 Bioanalyzer system. Only samples with RIN (RNA Integrity Number) value above 7 were taken into further investigation.

## Notes

### Competing Interest Statement

The authors have declared no competing interest.

